# Bioinstructive Orthogonally-crosslinked Ovoprotein Microgels for Modular Bioprinting

**DOI:** 10.64898/2026.06.17.732926

**Authors:** Suihong Liu, Vaibhav Pal, Joseph Christakiran Moses, Medine Dogan Sarikaya, Deepak Gupta, Miji Yeo, Valeriya Stepanyants, Yasar Ozer Yilmaz, Ibrahim T. Ozbolat

## Abstract

Bioprinting increasingly requires biomaterials that are not only printable, but structurally adaptive and biologically instructive. Here we establish an ovoprotein-derived microgel platform that couples intrinsic protein bioactivity with orthogonal interparticle photocrosslinking for modular bioprinting. Methacrylated ovoproteins yielded a photoresponsive protein-rich hydrogel matrix with retained proteomic complexity, tunable mechanics, and cell-regulatory biofunction. Endogenous tyrosine chemistry drove interparticle dityrosine coupling between ovoprotein microgels, producing cohesive, microporous, and intrinsically autofluorescent granular networks. The resulting systems displayed programmable rheology and broad compatibility across digital light processing, extrusion-based and aspiration-assisted bioprinting. Functionally, the ovoprotein microgel matrices attenuated sustained pro-inflammatory macrophage activation, promoted endothelial organization and host angiogenic invasion, and supported spheroid-mediated vascular morphogenesis with progressive sprouting, lumenization, branching and inosculation. It further enabled bioprinted osteogenic constructs with long-term maturation into bone-like mineralized tissues *in vitro*. These findings establish ovoprotein microgel-spheroid bioassembly as an adaptive, bioinstructive strategy for engineering vascularized and mineralized tissue constructs.

## Introduction

Bioprinting offers transformative opportunities for regenerative medicine by enabling spatially controlled deposition of cells and biomaterials into architecturally defined tissue constructs (*1*). Yet its translational potential remains fundamentally constrained by the available bioinks (*2, 3*). An ideal bioink must simultaneously satisfy competing requirements: cytocompatible processability, rheological programmability, and post-printing structural fidelity, and importantly an instructive microenvironment that directs cell fate, immune modulation, vascularization, and long-term tissue maturation (*4–6*). Protein-based bioinks are particularly compelling in this context, because their intrinsic biochemical complexity provides cell-instructive cues that are difficult to replicate in synthetic systems (*7, 8*). Despite sustained development from extracellular-matrix (ECM)-derived (*9*), animal-, marine- (*10, 11*), and plant-derived (*12, 13*) sources, protein-based bioinks frequently remain limited by source scarcity, narrow processing windows, insufficient mechanical integrity, and incomplete compatibility across multiple bioprinting modalities. These limitations highlight the need for protein-based material systems that combine intrinsic bioactivity with broad fabrication versatility across diverse bioprinting modalities.

A compelling route to versatility lies in reconceiving the material architecture of the bioink itself. Rather than treating a bioink as a continuous, homogeneous hydrogel that is cast around cells, one can instead build tissue constructs from discrete, independently fabricated material units whose composition, mechanics, and bioactivity are specified before assembly, a strategy that is inherently modular. In this bottom-up approach, microgels serve as material modules: structurally defined, compositionally tunable particles whose granular packing produces an inherently microporous and rheologically adaptable matrix that supports nutrient diffusion, cellular infiltration, and dynamic tissue remodeling (*14–17*). Multicellular spheroids function as complementary cellular modules, which are dense, self-organized aggregates with pre-established cell-cell contacts and tissue-specific biofunctions (*18, 19*), that can be independently cultured, phenotypically conditioned, and selectively incorporated into the microgel matrix as biologically coherent tissue units. The modularity of this approach lies precisely in this independence: each component is defined and optimized separately, yet assembled collectively through bioprinting to engineer heterogeneous, vascularized tissue architectures. The central unresolved challenge is to achieve interparticle cohesion within such granular assemblies without occluding the interstitial porosity that enables vascular morphogenesis, cell migration, and tissue remodeling (*20–22*). Existing strategies commonly introduce auxiliary surface functionalization, interpenetrating hydrogel phases, or post-fabrication processing steps each of which adds formulation complexity and erodes the generalizability that makes module-based biofabrication attractive in the first place (*15, 16, 23–26*).

Here, we address this challenge by establishing methacrylated ovoprotein microgels as a bioinstructive, orthogonally photocrosslinkable material platform for modular bioprinting. Derived from eggwhite, an abundant and structurally underexploited protein source, these microgels exploit the intrinsic tyrosine chemistry of ovoproteins to enable visible-light-mediated interparticle dityrosine coupling, yielding cohesive, microporous granular autofluorescent assemblies without surface engineering or secondary matrix reinforcement.

In the modular bioprinting framework introduced here, ovoprotein microgels and multicellular spheroids constitute independently defined modules whose coordinated assembly, directed by bioprinting, produces functional tissue constructs. We characterize the chemical and proteomic basis of the ovoprotein matrix after methacrylation, establish its rheological programmability and compatibility across digital light processing (DLP), extrusion-based bioprinting (EBB), and aspiration-assisted bioprinting (AAB) modalities, and demonstrate that the resulting constructs support immune modulation, endothelial organization, vascular morphogenesis, and long-term osteogenic biomineralization. These findings position orthogonally crosslinked ovoprotein microgels as a structurally adaptive, bioinstructive platform that brings the full promise of modular bottom-up bioprinting within practical reach.

## Results and Discussion

### 1. Ovoprotein-Based Photoresponsive Hydrogel

Ovoprotein derived from chicken eggwhite (EW), predominantly comprising ovalbumin-rich fractions, was chemically functionalized to impart photoresponsive crosslinking functionality. Primary amine residues within the protein backbone were selectively modified via methacrylation using methacrylic anhydride (MA), yielding methacrylated ovoprotein (EggMA), as illustrated in Fig. 1A. This covalent modification introduces pendant vinyl groups along the polypeptide chains, thereby enabling subsequent radical-mediated network formation (*27*), while largely preserving the native protein conformation and inherent bioactive motifs. The installed methacrylate functionalities rendered EggMA amenable to photoinitiated polymerization. In the presence of the water-soluble photoinitiator lithium phenyl-2,4,6-trimethylbenzoylphosphinate (LAP), exposure to visible light (λ = 405 nm) rapidly generates free radicals that initiate crosslinking of the vinyl groups, resulting in the formation of a covalently interconnected hydrogel network.

**Figure 1.**
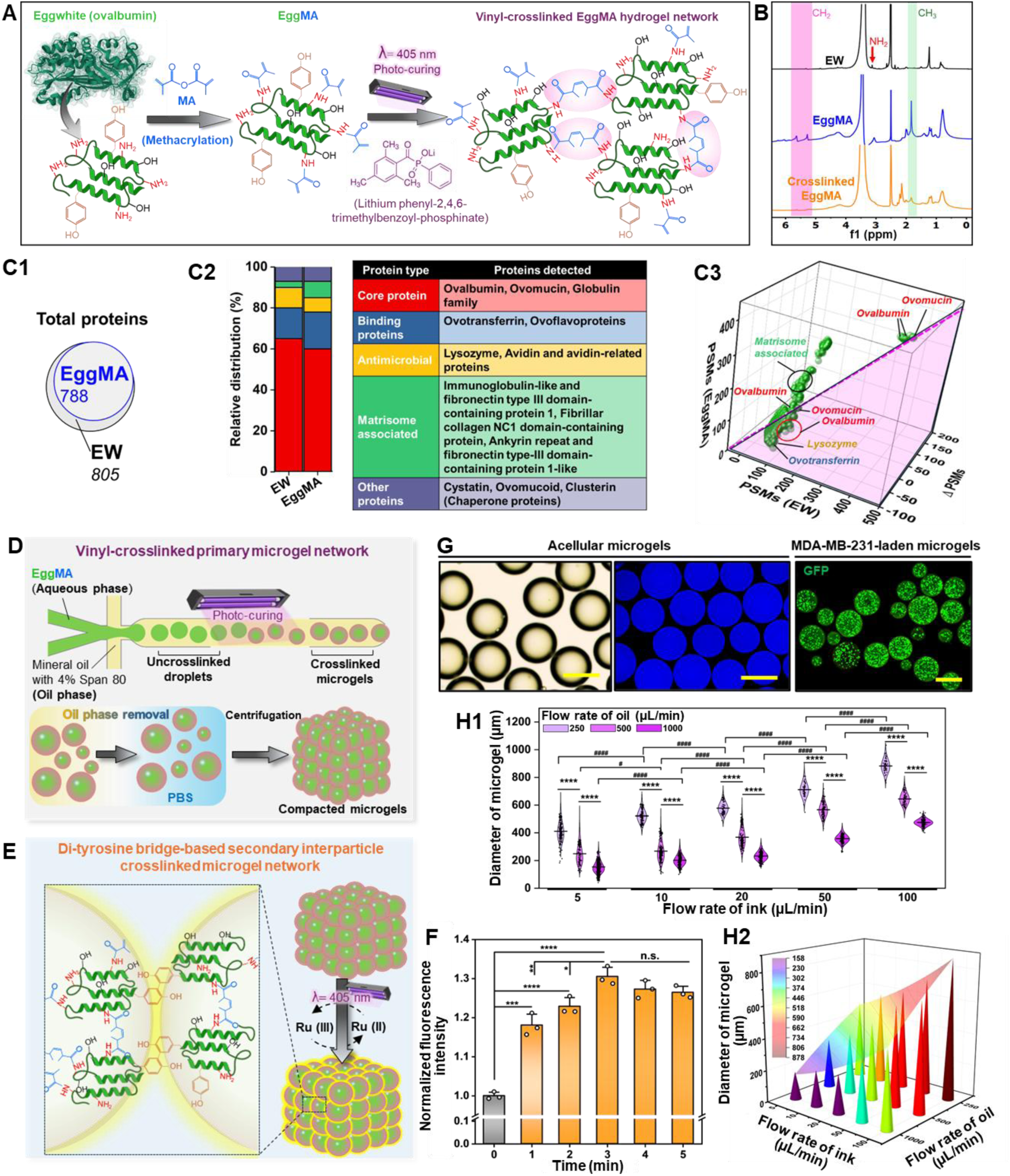
Engineering ovoprotein microgel networks through molecular functionalization and orthogonal interparticle crosslinking. A, Schematic illustrating methacrylation of eggwhite-derived ovoproteins and subsequent visible-light photocrosslinking to form the primary vinyl-crosslinked EggMA hydrogel network. **B,** ^1^H NMR spectra of EW, EggMA, and photocrosslinked EggMA hydrogel, confirming methacrylation and vinyl-crosslinked network formation. **C1–C3,** Proteomic evaluation of methacrylation effects on EggMA protein composition: **C1,** Number of proteins identified in native EW and EggMA; **C2,** Functional categorization and relative proportion of proteins retained in EggMA; **C3,** Correlation of normalized peptide-spectrum match (PSM) values between native EW and EggMA, showing preserved global protein abundance with protein-specific deviations after methacrylation. **D,** Schematic of microfluidic generation of vinyl-crosslinked EggMA microgels, followed by oil-phase removal and compaction into densely packed granular assemblies. **E,** Representative bright-field and autofluorescence images of acellular EggMA microgels and GFP^+^ MDA-MB-231-laden microgels; scale bar, 100 μm. **F1,** Diameter distribution of EggMA microgels generated under different dispersed-phase (ink) and continuous-phase (oil) flow conditions; (data were presented as mean ± standard deviation (SD), *n* ≥ 100 microgels per condition. \*\*\*\**P*<0.0001 denotes comparisons between ink flow rates at fixed oil flow rate. ^#^*P*<0.05 and ^####^*P*<0.0001 denote comparisons among oil flow rates at a fixed ink flow rate.) **F2,** 3D Mapping of microgel diameter as a function of ink and oil flow rates. **G,** Schematic of orthogonal secondary interparticle crosslinking through Ru/SPS-mediated dityrosine bond formation, yielding covalently integrated ovoprotein microgel networks. **H,** Time-dependent fluorescence analysis of dityrosine formation during secondary interparticle crosslinking, normalized to uncrosslinked microgels at 0 min (data were presented as mean ± SD, *n*=3. \**P*<0.05, \*\**P*<0.01, \*\*\**P*<0.001, and \*\*\*\**P*<0.0001, n.s., not significant).

The successful methacrylation and subsequent vinyl crosslinking capability of EggMA were validated by ^-1^H NMR spectroscopy (Fig. 1B). Compared to native EW, EggMA exhibited the emergence of characteristic resonances corresponding to methacrylate vinyl protons (–CH₂, δ ≈ 5.3–6.2 ppm) and methyl groups (–CH₃, δ ≈ 1.8 ppm), confirming the covalent incorporation of methacrylate functionalities onto the protein backbone. The accompanying attenuation of signals associated with lysine-derived primary amine environments further indicated consumption of free –NH₂ groups during functionalization. Consistently, 2,4,6-trinitrobenzenesulfonic acid (TNBS) analysis revealed a degree of functionalization of approximately 76%. Following photocrosslinking, attenuation of the vinyl proton signals further substantiates the participation of these unsaturated groups in radical-mediated network formation, collectively verifying the efficient functionalization and crosslinking reactivity of the ovoprotein-derived macromer. The biofunctionality of protein-based biomaterials is dictated by their constituent protein repertoire; therefore, a comprehensive proteomic analysis was performed to characterize the protein composition of EggMA following methacrylation (Figs. 1C1–C3). Compared to native EW, EggMA exhibited a modest reduction in the total number of identified proteins (788 versus 805, Fig. 1C1), likely attributable to partial loss, structural alteration, or reduced detectability of certain protein species during chemical functionalization. Despite this slight decrease, the global proteomic landscape remained largely preserved (Fig. 1C2), with core protein constituent-including ovalbumin, ovomucoid, and globulin families–retaining dominant representation. Functional classification further confirmed the preservation of key bioactive protein groups, including binding proteins (e.g., ovotransferrin), antimicrobial proteins (e.g., lysozyme and avidin-related proteins), and matrisome-associated proteins, with only minor variations in relative abundance, indicating that the intrinsic biofunctional diversity of EW is largely maintained in EggMA. To further interrogate the functional relevance of the retained proteome, 181 ovoproteins identified in EggMA were subjected to Reactome pathway analysis based on their human orthologs, together with gene ontology (GO) enrichment analysis, revealing conserved involvement in ECM organization, immune regulation, and protein-binding-associated biological processes (Figs. S1–S2).

Moreover, quantitative proteomic analysis was performed using spectral counting based on normalized peptide-spectrum match (PSM) values to compare protein abundance between EW and EggMA. Correlation analysis of protein abundance (Fig. 1C3) revealed a strong overall concordance between the two biomaterials, while also highlighting protein-specific deviations arising from chemical modifications. Proteins that were closely aligned along the diagonal exhibited minimal perturbation, suggesting that their structural integrity and detectability remain mostly unaffected by methacrylation. In contrast, subsets of proteins deviating from this correlation reflect differential susceptibility to modifications, where proteins enriched in accessible primary amine residues, such as ovotransferrin and lysine-rich domains, are more prone to methacrylation, potentially altering their conformation, solubility, or ionization efficiency and thereby influencing their apparent abundance in proteomic analysis. Conversely, proteins with more compact or structurally constrained architectures may exhibit reduced accessibility to reactive groups and thus retain protein abundance profiles comparable to native EW. This selective functionalization behavior underscores the heterogeneous reactivity of ovoprotein components and indicates that methacrylation is not uniformly distributed across all protein species but is instead governed by the protein structure, accessibility, and local chemical environment. Importantly, key structural and bioactive proteins, including ovalbumin and ovotransferrin, remain prominently represented after the modification, indicating that essential biological functions are preserved despite partial chemical derivatization, thereby reinforcing the capacity of EggMA to retain intrinsic bioactivity while enabling a tunable physicochemical modification.

Following functionalization, the sol–gel transition behavior and crosslinking capability of EggMA at varying concentrations (5, 6, 10, 15, and 20%, w/v) were first evaluated using vial inversion assays (Fig. S3A). Rapid and concentration-dependent gelation was observed upon photoirradiation, with increasing EggMA content yielding progressively more robust and self-supporting hydrogel constructs. In contrast, formulations at or below 5% exhibited insufficient network formation, indicative of limited crosslinking efficiency. Rheological analysis further elucidated the photoinitiated gelation kinetics, characterized by a rapid increase in storage modulus (*G′*) upon 405 nm irradiation, exceeding the loss modulus (*G″*) and signifying the formation of a solid-like network (Fig. S3B). The plateau *G′* increased with increasing EggMA concentration (Fig. S3C), consistent with enhanced crosslinking density. Frequency sweep measurements further confirmed the predominantly elastic behavior (*G′* ≫ *G″*) with minimal frequency dependence, indicative of stable, covalently crosslinked hydrogel networks. Concordantly, compressive mechanical testing revealed a pronounced concentration-dependent enhancement in stiffness, with increasing EggMA concentration yielding significantly higher compressive stress and modulus (Figs. S3D-E), further corroborating the formation of denser and mechanically reinforced hydrogel networks at elevated EggMA concentrations. Subsequently, the cytocompatibility and cellular response of functionalized EggMA hydrogel at varying concentrations (6, 10, 15, and 20% w/v) were evaluated with normal human lung fibroblasts (NHLFs) and green fluorescent protein (GFP^+^) metastatic breast cancer cells (MDA-MB-231). High viability of NHLFs seeded on and encapsulated within EggMA was maintained across all tested concentrations over time, with minimal cell death observed (Fig. S4A), as further confirmed by quantitative analysis showing >85% viability without significant differences among groups (Fig. S4B). Proliferation assays revealed progressive cell growth, with a marked increase from Day 3 to 5 across all groups (Fig. S4C), indicating that methacrylation and photocrosslinking preserve cytocompatibility. Cellular behavior exhibited a stiffness-dependent response. In cell-seeded constructs, intermediate to high concentrations (10-20%) supported robust spreading and elongation (Fig. S4A), whereas low concentrations (6%) provided insufficient structural support and limited cell extension. In contrast, an opposite trend was observed in cell-laden hydrogels, where cellular spreading progressively decreased with increasing EggMA concentration, with minimal spreading at 20% due to excessive matrix stiffness. A comparable trend was observed in MDA-MB-231-laden constructs (Fig. S5A), where cells exhibited progressive spreading and elongation at low to intermediate concentrations (6–15%), while spreading was restricted at 20%. Quantitative analysis confirmed consistently high viability (∼90%) across all groups (Fig. S5B), along with sustained proliferative capacity over time (Fig. S5C). Collectively, these results demonstrate that methacrylated ovoprotein integrates efficient photocrosslinking with preservation of intrinsic protein bioactivity, yielding EggMA hydrogels with tunable physicochemical and mechanical properties. Moreover, this ovoprotein-based photoresponsive system exhibits robust cytocompatibility across a broad concentration range while enabling stiffness-mediated regulation of cellular behavior, underscoring its potential as a bioactive and instructive platform for tissue biofabrication.

### 2. Orthogonally Crosslinked Ovoprotein Microgel Networks

Granular hydrogel assemblies have emerged as advanced biomaterials for tissue engineering, wherein their modular and intrinsically microporous architecture decouples bulk construct formation from local cellular remodeling, thereby enabling efficient mass transport, cellular infiltration, vascular integration, and dynamic cell–matrix reciprocity, while affording injectability, printability, heterogeneity, and spatiotemporal tunability (*14, 16, 23*). Utilizing the photoresponsive ovoprotein hydrogel precursor (EggMA), monodispersed microgels were generated via microfluidic emulsification (Fig. 1D), in which aqueous EggMA droplets were formed under controlled interfacial shear within an oil phase and subsequently photopolymerized to yield vinyl-crosslinked primary EggMA microgels (E-MG). Following oil-phase removal and centrifugation, freely formable, densely packed microgels assemble into shape-adaptable granular networks with inherent interstitial porosity. However, stabilization of microgel-assembled architectures generally necessitates interparticle annealing or covalent integration to achieve structural integrity (*16, 24, 28*). Notably, the intrinsic presence of tyrosine residues within ovoproteins (Fig. 1A), together with their incorporation into vinyl-crosslinked E-MG, enables a secondary interparticle crosslinking mechanism via photoinduced dityrosine bond formation (Fig. 1E). This process, mediated by tris(2,2′-bipyridyl)dichlororuthenium(II) hexahydrate/sodium persulfate (Ru/SPS) photoredox catalysis under visible light irradiation (λ = 405 nm), generates tyrosyl radicals via Ru(II)/Ru(III) redox cycling, enabling intermolecular coupling into dityrosine-based covalent crosslinking between microgels. Time-resolved fluorescence analysis revealed a progressive increase in dityrosine-associated emission intensity, confirming the time-dependent formation of covalent dityrosine linkages and validating the efficacy of the secondary interparticle crosslinking process (Fig. 1F). The fluorescence signal reached a plateau after ∼3 min, suggesting reaction completion, likely arising from depletion of available tyrosine residues or quenching of radical intermediates. The effectiveness of Ru/SPS-mediated interparticle crosslinking was further demonstrated at the construct level by forming thin E-MG layers and rectangular E-MG constructs (Supplementary Movie 1). Compared with non-crosslinked E-MG assemblies, Ru/SPS-treated E-MG exhibited markedly improved cohesion and structural stability, confirming the role of photo-induced dityrosine coupling in stabilizing the microgel network (Supplementary Movie 2). Importantly, this orthogonal crosslinking strategy reinforces network cohesion while preserving the integrity of the primary vinyl-derived network, enabling interparticle stabilization without perturbing the bulk network architecture. In contrast to conventional protein-based microgel systems, this approach obviates the need for auxiliary precursor functionalization, microgel surface engineering, multiphase or composite material design, or secondary matrix-mediated processing (*14, 16, 17, 23*).

Uniform and highly monodispersed spherical E-MG were generated using 10% EggMA, as shown in Fig. 1G. Microgel size was precisely regulated by tuning the flow rates of the dispersed EggMA phase and the continuous mineral oil phase (Fig. 1H1, Fig. S6A). Specifically, at a constant ink flow rate, increasing the flow rate of oil phase reduced microgel diameter via enhanced shear-induced droplet breakup, while decreasing the ink flow rate at fixed flow rate of oil conditions similarly yielded smaller microgels due to reduced droplet volume at formation. Across the investigated flow regime, microgel diameters were tunable over a broad range (∼100–900 μm), demonstrating robust scalability and precise control over microgel fabrication. 3D Mapping of microgel size as a function of dispersed (ink) and continuous (oil) phase flow rates (Fig. 1H2) revealed a predictable flow-dependent size-regulation profile, wherein microgel diameter increases with increasing ink flow rate and decreases with increasing oil flow rate. This behavior is consistent with classical droplet microfluidics governed by the interplay between volumetric influx and interfacial shear (*29–31*), enabling predictive tuning of microgel size through flow-rate modulation. Notably, microgels exhibited spherical morphology across all conditions, with circularity values consistently exceeding 0.98 (Fig. S6B), reflecting stable flow-focusing dynamics, uniform interfacial tension, and highly reproducible droplet generation. A comparable flow-governed scaling behavior was observed for microgels derived from 15% EggMA (Figs. S6C–E). Notably, deviations from ideal sphericity arose under regimes of high dispersed-phase flux coupled with low continuous-phase flow (e.g., 250 μL/min), where insufficient interfacial shear compromised droplet pinch-off dynamics. This imbalance led to jetting-like elongation and droplet coalescence, ultimately reducing circularity and undermining morphological fidelity.

Photoluminescence mapping unveiled a pronounced intrinsic autofluorescence signature in photocrosslinked EggMA, characterized by well-resolved excitation–emission matrices across all tested concentrations (6–15% w/v, Fig. S7), and faithfully preserved in the derived E-MG, as visualized by fluorescence microscopy using a DAPI filter set (357/44 nm excitation, 447/60 nm emission) (Fig. 1G, Figs. S6A, S6C). This baseline autofluorescence can be ascribed to intrinsic protein chromophores and clustered emissive domains within the conformationally constrained network, consistent with previous studies on albumen–gelatin-based luminescent glasses (*32*). Two emission features were resolved, including a high-energy band (∼320–340 nm emission under ∼280–300 nm excitation) from aromatic residues and a broader band (∼400–480 nm emission under ∼320–380 nm excitation) associated with initial dityrosine formation. Comparative analysis of LAP-only and orthogonally crosslinked (LAP + Ru/SPS) hydrogels revealed a marked enhancement of fluorescence upon secondary crosslinking (Fig. S8), with emission extending into the ∼500–600 nm range and a visible transition from blue to orange. This shift arises from Ru-mediated photoredox generation of tyrosyl radicals and subsequent dityrosine coupling, which increases the density of conjugated chromophores. Coupled with network-induced conformational restriction, this promotes chromophore clustering and suppresses non-radiative relaxation, thereby amplifying emission. Together, these results establish orthogonally crosslinked ovoprotein networks as an intrinsically fluorescent biomaterial that reinforces the structural integrity of microgel-based microporous scaffolds and provides a built-in, label-free optical modality for real-time interrogation of scaffold architecture.

E-MG assemblies with tunable packing densities were established via controlled centrifugations, enabling systematic modulation of the resulting microporous architecture (Fig. S9). The porous network is visualized using intrinsic autofluorescence of E-MG in combination with fluorescein isothiocyanate (FITC)–dextran infiltration, captured by multiphoton microscopy (Fig. S10), and further reconstructed into 3D architectures using Imaris (Fig. S9A). Both assemblies were porous, with low packing forming larger voids and high packing yielding more compact, less interconnected networks. Quantitative assessment of porosity, derived from both 2D projections and 3D volumetric reconstructions, showed consistent trends (Figs. S9B–C), with significantly higher porosity of E-MG in the low-packing group (∼10–15%) compared to the high-packing counterpart (∼5–7%) (*\*\*\*P* < 0.001, *\*P* < 0.05). Moreover, the cytocompatibility of E-MG was further validated using both NHLFs and MDA-MB-231 cells (Fig. S11). In cell-seeded configurations, MDA-MB-231 and NHLFs adhered to microgel surfaces and exhibited progressive spreading, forming interconnected networks with extensive surface coverage after 7 days (Figs. S11A, S11C). In parallel, MDA-MB-231-laden E-MG exhibited a homogeneous initial cell distribution (Fig. 1G), followed by pronounced proliferation and colonization throughout the microgel volume by Day 7 (Fig. S11B). Quantitative analysis indicated high cell viability (>90%) and substantial cellular coverage ratio on microgels (>80%) across all conditions (Figs. S11D-E), indicating a permissive microenvironment for cell survival and active remodeling. Overall, orthogonally crosslinked ovoprotein E-MG networks offer a structurally robust and architecturally tunable platform with integrated optical and biological functions for tissue biofabrication.

### 3. Bioprinting of Ovoprotein-based Bulk Hydrogel and Microgel Systems

The versatility of the photocrosslinkable ovoprotein-based systems was rigorously interrogated across diverse bioprinting modalities to establish their potential as a versatile bioink and for functional supporting bath for tissue biofabrication. Prior to bioprinting, rheological properties of bulk EggMA hydrogel (E-HG), microgel assemblies (E-MG) with low and high packing densities, and the hybrid formulation E-HG@MG, prepared by mixing EggMA hydrogel and microgels at a 1:1 (v/v) ratio, were systematically evaluated to delineate their flow and deformation responses under bioprinting-relevant conditions. All formulations exhibited pronounced non-Newtonian shear-thinning behavior, with viscosity decreasing monotonically over several orders of magnitude with increasing shear rate (Fig. 2A), indicative of progressive microstructural disentanglement and lubrication-mediated flow under shear, thereby enabling efficient extrusion through confined geometries. Microgel-based systems (E-MG and E-HG@MG) displayed substantially elevated viscosity profiles compared to E-HG, reflecting the emergence of a jammed, particulate-dominated regime governed by interparticle contacts and frictional dissipation (*33–35*). Under oscillatory step-shear perturbations, these systems demonstrated rapid viscosity recovery following high-shear fluidization (Fig. 2B), evidencing pronounced thixotropic behavior of microgel networks, a prerequisite for maintaining filament continuity and integrity during, and after extrusion. Oscillatory strain sweeps further revealed a predominantly elastic response (*G′* ≫ *G″*) within the linear viscoelastic regime, followed by a sharp yielding transition at critical strain amplitudes (Fig. 2C). Notably, microgel-containing formulations exhibited enhanced *G′* and extended yield points (*G′* = *G″*), consistent with mechanically robust, load-bearing networks arising from densely packed and interlocked microgel assemblies. For instance, high-packing E-MG displayed higher moduli than low-packing counterparts (Fig. 2C). Yield stress measurements (Fig. 2D) further corroborated the enhanced resistance to flow in microgel-containing systems relative to the bulk hydrogel, reflecting the increasing contribution of interparticle interactions and packing-induced jamming. This behavior is consistent with the emergence of a stress-bearing, arrested state characteristic of soft granular materials, enabling structural stability under applied stress while permitting flow upon yielding (*36*). In conjunction with rapid structural recovery under cyclic shear (Fig. 2B), these rheological features indicate dynamic network reconfiguration and reversible interparticle interactions, characteristic of self-healing behavior, thereby supporting bioprinting modalities that require transient fluidization under localized stress and rapid stabilization of deposited structures.

**Figure 2.**
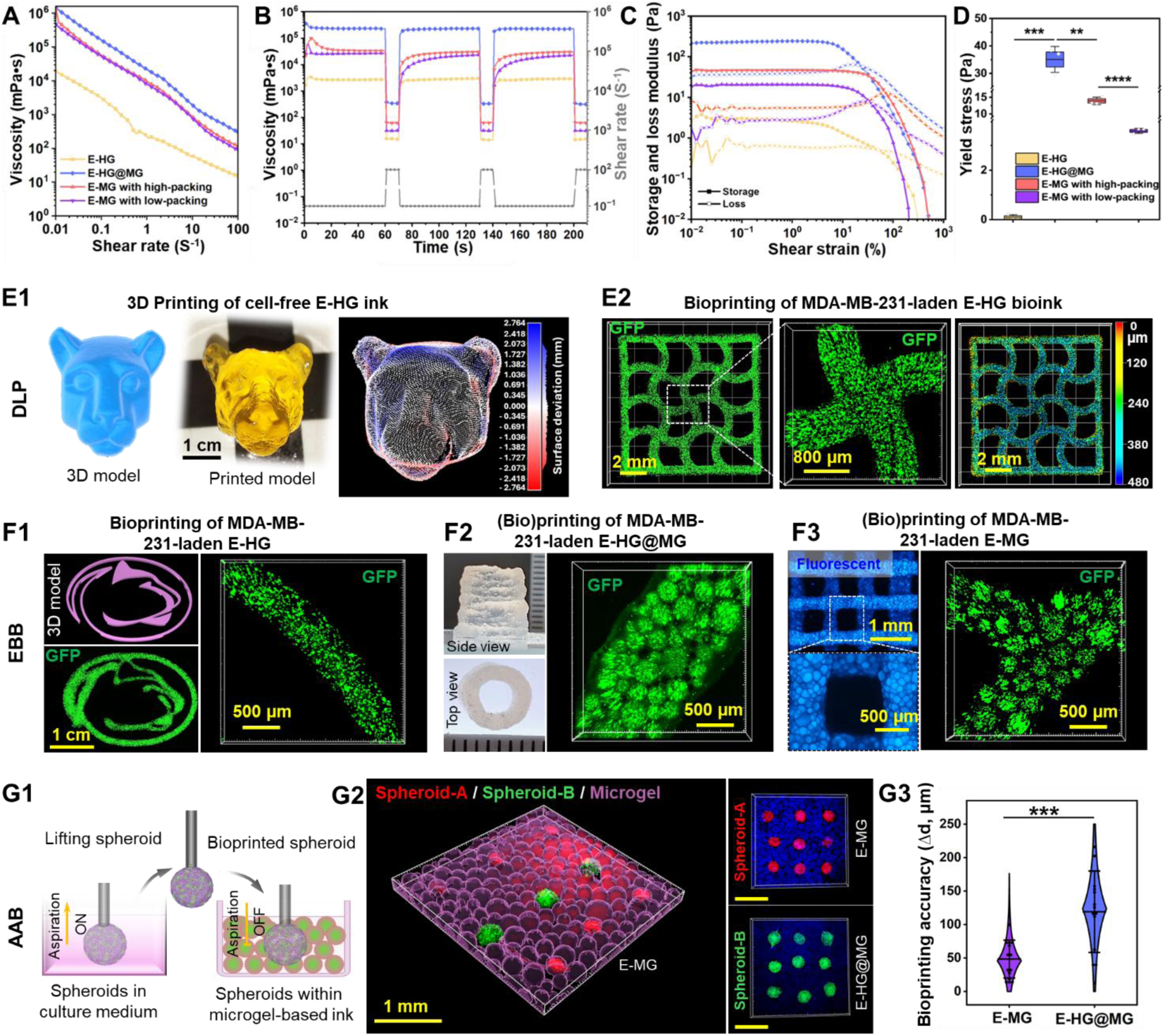
Rheological programming and multimodal bioprinting of ovoprotein hydrogel and microgel (bio)inks. A, Shear-rate-dependent viscosity profiles of E-HG, E-HG@MG, and E-MG with low- and high-packing densities. **B,** Cyclic step-shear recovery analysis showing reversible viscosity recovery after high-shear fluidization. **C,** Oscillatory strain sweep showing *G′* and *G″* and yielding behavior. **D,** Yield stress of different ovoprotein-based bioink formulations. (Data were presented as mean ± SD (*n* = 3); *\*\*P* < 0.01, *\*\*\*P* < 0.001 and *\*\*\*\*P* < 0.0001.). **E1,** Digital light processing (DLP) of acellular E-HG into a centimeter-scale Penn State University Nittany Lion head model, with point-cloud surface deviation analysis. Scale bar, 1 cm. **E2,** DLP of GFP^+^ MDA-MB-231-laden E-HG constructs with depth-coded confocal projection, showing homogeneous cell distribution and viability. Scale bar, 2 mm, 800 μm. **F1,** Extrusion-based bioprinting (EBB) of GFP^+^ MDA-MB-231-laden E-HG, showing shape fidelity in a bioprinted PSU logo and homogeneous cell distribution within printed filaments. **F2,** EBB of acellular and GFP^+^ MDA-MB-231-laden E-HG@MG bioink, showing fabrication of a hollow tubular construct and a cell-laden filament. **F3,** EBB of acellular and GFP^+^ MDA-MB-231-laden E-MG bioink, showing intrinsic autofluorescence of the printed microgel grid and viable cell distribution within the E-MG construct. **G1,** Schematic of aspiration-assisted bioprinting (AAB) for spatial deposition of spheroids within microgel-based matrices. **G2,** Representative 3D reconstruction and fluorescence images showing patterned spheroid (spheroid-A: hMSC spheroids, red; spheroid-B: vascular spheroids, green) populations within E-MG and E-HG@MG constructs. Scale bar, 1 mm. **G3,** Quantification of bioprinting accuracy in E-MG and E-HG@MG matrices (data were presented as mean ± SD, *n*=14; *\*\*\*P* < 0.001.)

Building upon these rheological characteristics, the printability and versatility of the ovoprotein-based systems were further interrogated across multiple bioprinting modalities (Figs. 2E–G). The photocrosslinkable E-HG demonstrated excellent compatibility with DLP, enabling rapid fabrication of complex geometries with high structural fidelity, including centimeter-scale constructs such as the Penn State University (PSU) Nittany Lion head (Fig. 2E1 and Supplementary Movie 3). Quantitative surface deviation analysis, based on point-cloud distance mapping between the printed construct and its digital model, revealed minimal geometric deviation, confirming high printing accuracy and reconstruction of fine architectural features. In parallel, GFP^+^ MDA-MB-231-laden E-HG constructs were successfully bioprinted with DLP (Fig. 2E2), with depth-coded confocal projection revealing uniform 3D cell distribution throughout the bioprinted constructs, indicative of uniform cell encapsulation during photopolymerization. These results demonstrate that the E-HG system enables high-resolution bioprinting with precise geometric conformity, uniform cell encapsulation, and scalability for rapid fabrication of cell-laden constructs. When tested for EBB, GFP^+^ MDA-MB-231-laden E-HG supported continuous filament deposition and fabrication of a PSU logo architecture (Fig. 2F1); however, its relatively low viscosity and limited yield stress constrained structural fidelity and stability in larger-scale constructs. In contrast, microgel-based systems (E-HG@MG and E-MG) exhibited enhanced printability and shape fidelity during extrusion. The E-HG@MG formulation enabled the fabrication of stable, centimeter-scale acellular hollow tubular architectures (Fig. 2F2), and GFP^+^ MDA-MB-231-laden filament deposition, where MDA-MB-231 cells preferentially encapsulated within the microgel phase of the hybrid bioink (Fig. 2F2). Notably, high-packed, acellular E-MG exhibited superior print fidelity, enabling the fabrication of well-defined grid architectures with intrinsic blue autofluorescence (Fig. 2F3), while preserving interconnected microporosity and robust interparticle cohesion, characteristic of a jammed granular network (Fig. S12). In contrast, low-packed, GFP^+^ MDA-MB-231-laden E-MG demonstrated favorable extrusion behavior, with cells retained at high local density within individual microgels after deposition (Fig. 2F3), indicative of a permissive microenvironment for cell encapsulation and protection during processing.

Beyond conventional bioprinting modalities, microgel-based systems were further interrogated using AAB, an emerging technique that enables the precise manipulation and spatial organization of spheroids, organoids, and microtissues as high-density building blocks for bottom-up tissue assembly (*37, 38*). Leveraging the yield-stress behavior and rapid self-recovery of the microgel-based matrices, both E-HG@MG and low-packing E-MG served as supportive, reconfigurable environments for spheroid placement (Figs. 2G1-G3). These matrices enabled localized fluidization during spheroid deposition, followed by rapid structural recovery to spatially stabilize deposited spheroid. Two distinct spheroid populations (hMSC spheroids, red; vascular spheroids, green) were patterned with high spatial fidelity and maintained discrete positioning within the E-MG matrix without undesired displacement or fusion (Fig. 2G2 and Supplementary Movie 4) and independently patterned within E-MG and E-HG@MG matrices, respectively. Quantitative analysis (Fig. 2G3) of positional accuracy revealed sub-50 μm deviation for E-MG matrices, significantly outperforming E-HG@MG, attributable to the enhanced confinement and positional stability imparted by the jammed microgel network. Importantly, the combination of a yield-stress support matrix and high-density cellular building blocks enables precise bottom-up assembly of multicellular constructs, highlighting the suitability of the ovoprotein microgel system for scalable and architecturally defined tissue fabrication.

### 4. Immunocompatibility of Ovoprotein Microgel Matrices

Given the xenogeneic origin of ovoproteins derived from avian sources, evaluating their immunological response is essential, as interspecies disparities in protein structure, epitope presentation, and post-translational modifications may potentiate unintended immune activation upon interaction with human cells (*39, 40*). Such responses are particularly relevant for implanted or cell-laden constructs, where biomaterial–immune cell interactions shape inflammation resolution, matrix remodeling, vascular ingrowth, and host tissue infiltration, thereby determining regenerative performance (*39, 41*). Among immune effectors, macrophages play a pivotal role in mediating biomaterial-associated host responses, functioning as highly plastic regulators that integrate biochemical and biophysical cues from the microenvironment to direct inflammation, matrix remodeling, and tissue regeneration.

Macrophage polarization exists along a dynamic spectrum, broadly characterized by pro-inflammatory (M1-like) and pro-regenerative (M2-like) phenotypes, with transitional and hybrid states reflecting functional plasticity in response to environmental stimuli (*42, 43*). Therefore, delineating the polarization landscape induced by ovoprotein-based matrices is essential for assessing their immunocompatibility and bioinstructive potential. To this end, we systematically interrogated the immunomodulatory properties of E-HG, E-MG, and E-HG@MG using human monocytic leukemia cell line (THP-1)-derived macrophages. Prior to matrix evaluation, a rigorous flow-cytometric workflow was established to resolve matrix-dependent macrophage phenotypes (Fig. S13). Parent events were gated to exclude debris, followed by doublet discrimination and selection of the CD68⁺ pan-macrophage population. Within this gate, polarization states were resolved using CCR7 as a pro-inflammatory marker and CD163/CD206 as reparative markers. The control conditions validated the assay stringency: unstimulated M0 macrophages remained largely negative for these markers (Figs. S13B1-B3), whereas interferon-gamma (IFN-γ) treatment generated highly enriched CCR7⁺ populations (>98%, Figs. S13C1-C3) and interleukin-4 (IL-4) treatment yielded strongly CD163⁺/CD206⁺ populations (>95%, Figs. S13D1-D3), thereby defining the M1- and M2-polarized reference states.

Against this benchmark, ovoprotein-based matrices elicited markedly different and time-evolving polarization trajectories (Fig. 3). E-HG was initially strongly M1-biased, with predominantly CCR7⁺ macrophages at Day 3 in both the CCR7/CD163 and CCR7/CD206 analyses (>90%, Fig. 3A1). By Day 7, the bulk hydrogel remained largely CCR7-retentive in the CCR7/CD163 gate (81.03%), with only a modest emergence of a CCR7⁺/CD163⁺ hybrid population (12.85%). In contrast, the CCR7/CD206 analysis revealed a striking shift toward a CCR7⁺/CD206⁺ hybrid state (99.25%), while single-positive CD206⁺ cells remained negligible, suggesting progression from a purely inflammatory phenotype toward a plastic, transitional macrophage state rather than full M2 polarization. In contrast, E-MG elicited a more plastic macrophage response on Day 3, with reduced CCR7-only fractions (≤80% in the CCR7/CD163 and CCR7/CD206 gates) accompanied by the emergence of CCR7⁺/CD163⁺ (11.31%) and CCR7⁺/CD206⁺ (8.35%) subpopulations (Fig. 3B1). By Day 7, this phenotypic shift became substantially more pronounced: the CCR7⁺/CD163⁺ fraction increased to 80.57%, whereas the CCR7-only population in the CCR7/CD163 gate declined to 11.27%. In parallel, the CCR7/CD206 analysis showed a reduction in the CCR7-only fraction to 65.80% together with expansion of the CCR7⁺/CD206⁺ population to 20.54%. These data indicate that E-MG did not sustain a purely inflammatory macrophage state, but instead promotes transition toward a hybrid, remodeling-associated phenotype with enhanced functional plasticity. The most reparative immunophenotype was observed in E-HG@MG. At Day 3, this hybrid matrix already exhibited a heterogeneous macrophage landscape (Fig. 3C1), comprising appreciable single-positive CD163⁺ (17.75%) and CD206⁺ (19.35%) populations together with CCR7⁺/CD163⁺ (15.75%) and a dominant CCR7⁺/CD206⁺ hybrid fraction (52.71%). By Day 7, this distribution shifted further toward reparative polarization, with marked enrichment of CD163⁺ and CD206⁺ populations (59.21% and 65.54%, respectively), accompanied by a substantial reduction in the CCR7-only fractions to 21.13% and 7.35% (Fig. 3C1). These phenotypic trends were further corroborated by immunofluorescence imaging of retrieved macrophages (Fig. 3A2–C2). Collectively, the bulk E-HG maintained a predominantly pro-inflammatory phenotype, the ovoprotein microgel-containing matrices progressively redirected macrophages toward hybrid and reparative states with E-MG favoring phenotypic plasticity and E-HG@MG exhibiting the most pronounced late-stage M2-like polarization.

**Figure 3.**
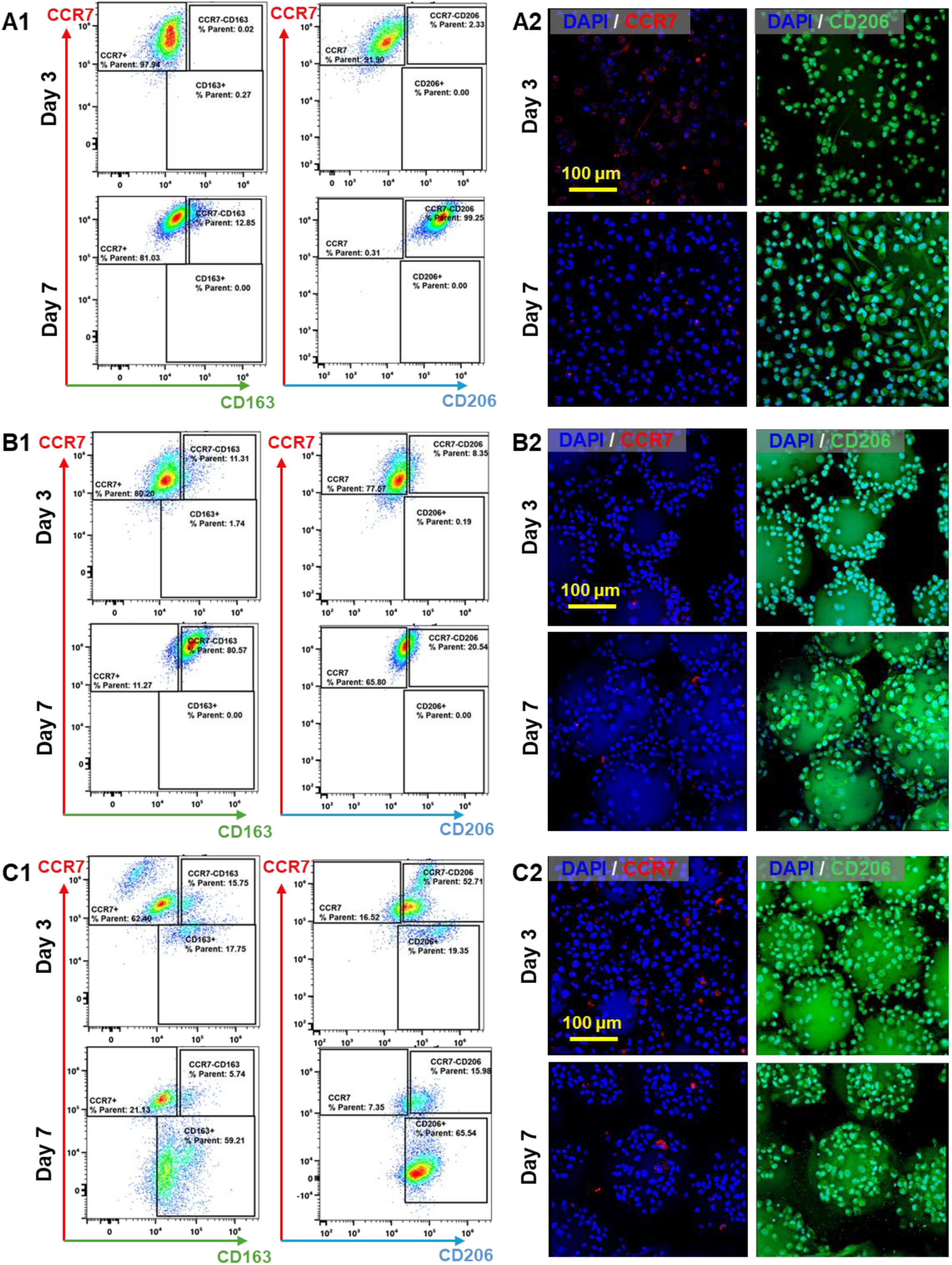
Immunomodulatory response of ovoprotein hydrogel and microgel bioink formulations in naïve M0 macrophages. Immunophenotypic analysis of THP-1-derived macrophages retrieved from E-HG (**A1-A2**), E-MG (**B1-B2**), and E-HG@MG (**C1-C2**) constructs at Days 3 and 7. **A1–C1,** Flow cytometric profiles showing distributions of CCR7⁺, CD163⁺, and CD206⁺ populations. **A2–C2,** Corresponding immunofluorescence images of retrieved macrophages stained for CCR7 or CD206, with nuclei counterstained by DAPI. Scale bar, 100 μm.

In addition, the widely used methacrylated gelatin (GelMA) bulk hydrogel (GelMA-HG) and hybrid GelMA-HG@MG, prepared by mixing GelMA hydrogel and microgel at 1:1 (v/v) ratio, were included as benchmark controls and exhibited a markedly divergent immunophenotypic response relative to the ovoprotein-based systems (Figs. S14 and S16). GelMA-HG remained overwhelmingly CCR7-dominant throughout culture, with the CCR7⁺ fraction increasing from 74.25% at Day 3 (Figs. S14A1-A3) to 98.26% at Day 7 (Figs. S14C1-C3) in the CCR7/CD163 analysis, while single-positive CD163⁺ and CD206⁺ populations remained essentially undetectable. GelMA-HG@MG showed only limited attenuation of this pro-inflammatory profile: although a modest CCR7⁺/CD163⁺ hybrid population emerged by Day 7 (21.09%), macrophages remained predominantly CCR7⁺ in both analyses (75.54% and 94.57%), with negligible single-positive CD163⁺ (0.29%) and CD206⁺ (0.04%) fractions (Figs. S14 B1-B3, D1-D3). Notably, a pure GelMA microgel matrix was not pursued, owing to its limited interparticle crosslinking capacity and the resulting inability to establish a mechanically stable granular construct. Together, these results indicate that the immunomodulatory behavior observed in the ovoprotein systems is not a generic feature of methacrylated protein matrices, but rather a matrix-specific response associated with the EggMA microgel platform and its capacity to redirect macrophages away from sustained pro-inflammatory activation toward hybrid and reparative polarization states.

This immunophenotypic divergence is mechanistically concordant with the coupled biophysical and biochemical features of the ovoprotein microgel microenvironment. Macrophage phenotype is highly sensitive to biomaterial architecture, including porosity, interfacial geometry, confinement, and matrix organization, all of which can bias inflammatory versus reparative programs (*43–45*). Relative to bulk E-HG, the open, microporous, and dynamically remodelable E-MG and E-HG@MG matrices likely create a less confinement-dominated but more immunoregulatory microenvironment, where interfacial geometry, groove–ridge-like microtopography (*46*), local mechanical cues, and improved cell–matrix accessibility favor macrophage phenotypic diversification over persistent CCR7-dominant activation (*47*). Moreover, the retained EggMA protein repertoire, including ovalbumin, ovomucin-related proteins, and lysozyme C (Fig. S1C), together with enrichment of cytokine signaling, IL-4/IL-13 signaling, ECM organization, and membrane–ECM interaction pathways (Fig. S2), indicates that the ovoprotein matrix is immunologically active rather than inert. In particular, lysozyme is recognized as a regulator of innate immune signaling and inflammatory resolution (*48*), whereas mucin-associated proteins are closely associated with immune-interface regulation (*49*). Overall, these architectural and biochemical attributes likely account for the reduced persistence of CCR7-dominant states and the enhanced emergence of hybrid and reparative macrophage phenotypes in the ovoprotein microgel matrices, thereby establishing a more immunologically permissive microenvironment than bulk hydrogel or GelMA-based controls. These findings position ovoprotein microgel assemblies as immunomodulatory biomaterials capable of coupling structural design with favorable host-response regulation.

### 5. Ovoprotein Microgel Assemblies Orchestrate Vascularization

Ovoprotein-derived microgel assemblies, by virtue of their intrinsic bioactivity and microporous architecture, were investigated for their ability to direct endothelial behavior and orchestrate vascular morphogenesis across multiple biological scales. At the bulk hydrogel matrix level, EggMA supported the formation of interconnected capillary-like networks using the endothelial tube formation assay, outperforming conventional matrices such as GelMA in terms of cellular coverage, vessel length and junction density (Fig. S17). Immunostaining confirmed robust CD31 expression and cytoskeletal organization, indicative of stabilized endothelial junctions and functional network formation (Fig. S18). These results suggest that ovoprotein-based hydrogel provides a permissive and bioinstructive microenvironment for endothelial self-assembly. At the individual microgel level, HUVECs cultured on E-MG microgels maintained high viability (>90%, Figs. S19A–B) and exhibited progressive spreading, migration, and colonization, ultimately forming continuous multicellular networks across adjacent microgels (Figs. S19C–E). Consistently, robust CD31 expression was observed throughout the E-MG constructs (Fig. S20), accompanied by well-defined intercellular junctions, while quantitative analysis revealed a time-dependent increase in both cellular coverage on microgels and CD31 fluorescence intensity (Fig. S20B), indicative of active endothelial remodeling and progressive functional maturation within the microgel microenvironment.

Building upon these observations, vascular spheroids were bioprinted within E-MG and E-HG@MG using AAB (Fig. 4A). The subsequent endothelial dynamics and vascular morphogenesis within these biofabricated constructs, composed of ovoprotein microgel-based matrices and high-density cellular building blocks, were systematically interrogated over the course of culture (Fig. 4B). Notably, vascular spheroids embedded within both E-MG and E-HG@MG exhibited sustained growth, proliferative expansion, and progressive outgrowth into the surrounding microgel environment, accompanied by gradual cellular dispersion and spheroid fusion over culture (Figs. S21–S22). Cells emanating from adjacent spheroids (distance: 800 μm) underwent pronounced spreading, migration, and inter-spheroid bridging, indicative of coordinated cellular communication and early network integration (Fig. S23). Importantly, the intrinsically porous E-MG matrix further supported extensive cell spreading, migration, and spatial colonization throughout large-scale constructs (Fig. S24), underscoring the capacity of the granular microenvironment to accommodate long-range cellular invasion and multicellular organization.

**Figure 4.**
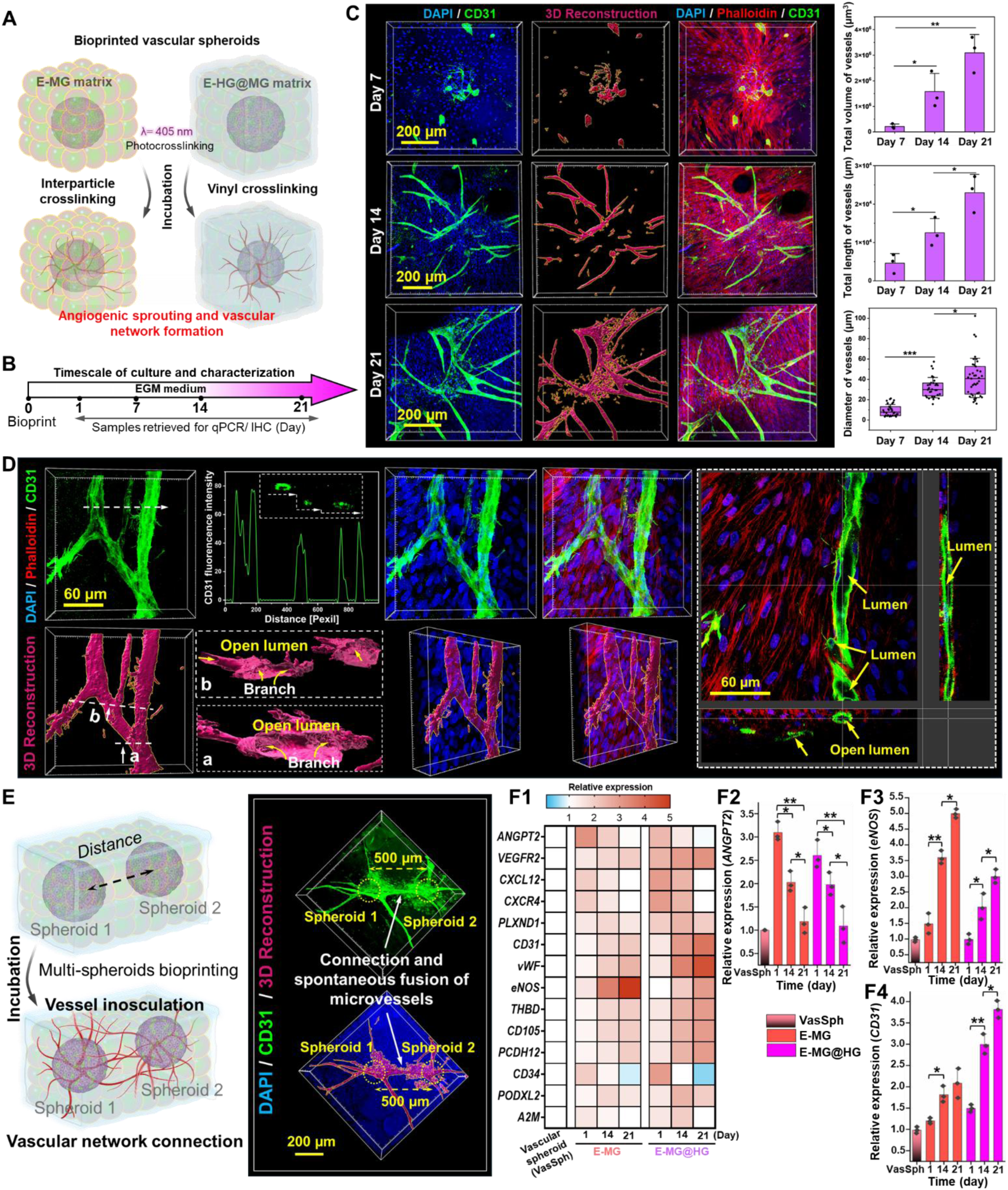
Ovoprotein microgel-based matrices direct spheroid-mediated vascular morphogenesis. A, Schematic of AAB of vascular spheroids into E-MG and E-HG@MG matrices, followed by interparticle or bulk photocrosslinking to form spheroid-loaded constructs. **B,** Culture and characterization timeline for vascular morphogenesis analysis. **C,** Representative confocal images of DAPI (blue), CD31 (green) and phalloidin (red) staining, together with 3D reconstruction of CD31 fluorescence via Imaris, showing time-dependent endothelial sprouting and vascular network formation from bioprinted vascular spheroids within microgel-based matrices. Quantification shows total vessel volume and length, as well as the diameter distribution of formed vessels over time (data were presented as mean ± SD, *n*=3; *\*P* < 0.05, *\*\*P* < 0.01, and *\*\*\*P* < 0.001.). **D,** High-resolution confocal imaging, 3D reconstruction, fluorescence intensity profiling and orthogonal sectioning of CD31⁺ vessel-like structures within E-HG@MG constructs at Day 21, revealing hierarchical branching architectures and patent luminal compartments. **E,** Schematic and representative 3D images showing multi-spheroid bioprinting and spontaneous microvascular interconnection between spatially separated spheroids (distance=500 μm). **F1–F4,** Heatmap and quantitative qPCR analysis of angiogenesis-associated gene expression in vascular spheroids cultured within E-MG and E-HG@MG matrices over time. **F1,** Heatmap showing temporal expression profiles of angiogenesis-, endothelial maturation- and vascular remodelling-associated genes. **F2–F4,** Relative expression of representative vascular markers, including *ANGPT2* (**F2**), *eNOS* (**F3**) and *CD31* (**F4**). Data were presented as mean ± SD (*n*=3); *\*P* < 0.05, *\*\*P* < 0.01 and *\*\*\*P* < 0.001.

Following spatial organization, vascular spheroids exhibited pronounced angiogenic sprouting and progressive microvascular self-organization within the ovoprotein microgel matrices. Time-resolved imaging revealed sustained endothelial sprouting, radial invasion, and the emergence of interconnected vascular networks with increasing branch complexity over culture (Fig. 4C). Quantitative analysis further confirmed this temporal evolution, showing significant increases in total vessel volume, cumulative vessel length and mean vessel diameter, consistent with progressive microvascular network expansion and remodelling (Fig. 4C). Moreover, 3D reconstruction of CD31^+^ vascular networks further delineated lumenized microvascular architectures with patent luminal compartments and hierarchically arborized branching patterns, consistent with native vascular morphogenesis rather than passive cellular redistribution or nonspecific outgrowth (Fig. 4D and Supplementary Movie 5). Cross-sectional CD31 fluorescence profiling, together with orthogonal section views, further verified the presence of patent open luminal compartments across vessels of varying calibers, including within bifurcating microvascular structures (Fig. 4D). Notably, spatially separated vascular spheroids within ovoprotein microgel matrices exhibited directed endothelial outgrowth toward one another, culminating in spontaneous microvascular interconnection and vascular inosculation across predefined spatial gaps (500 μm, Fig. 4E). This phenomenon was further observed in multi-spheroid configurations, in which three independently patterned spheroids established complex inter-spheroid microvascular anastomoses over separations of up to ∼1000 μm, ultimately yielding an integrated and hierarchically interconnected vascular network (Fig. S25). These microvascular networks were supported in both E-MG and E-HG@MG matrices (Fig. S26), with comparable vessel caliber and CD31 fluorescence intensity, indicating that both microenvironments effectively sustain endothelial organization and vascular maturation. E-HG@MG appeared to promote more elaborate branching and interconnection, suggestive of a denser and more reticulated vascular architecture. Notably, at the construct scale, extensive cellular migration, spatial colonization, and widespread microvascular network formation were observed throughout the entire E-MG matrix (Fig. S27), underscoring the capacity of the ovoprotein microgel system to support vascular integration across large volumes.

Consistent with the progressive endothelial outgrowth, lumenization, branching, and inter-spheroid inosculation observed morphologically, transcriptional profiling further substantiated a time-dependent pro-angiogenic maturation program within the ovoprotein microgel matrices (Figs. 4F1-F4). Relative to free-standing vascular spheroid controls, both E-MG and E-HG@MG induced upregulation of genes associated with endothelial activation, vascular remodeling, and network stabilization, including *ANGPT2, VEGFR2, CXCL12, CXCR4, CD31, vWF,* and *eNOS*. The progressive increase in *CD31* expression paralleled the emergence of CD31^+^ branched microvascular networks (Fig. 4C), whereas elevated *eNOS* and *vWF* are consistent with endothelial functional maturation and acquisition of a more stabilized vascular phenotype. Concurrent activation of the *CXCL12/CXCR4* axis further supports chemotactic guidance and coordinated endothelial migration, in agreement with the directed sprouting and long-range interconnection observed between spatially separated spheroids (Figs. 4C-E, Figs. S25, S27). Likewise, the increased expression of *ANGPT2* and *VEGFR2* supports ongoing angiogenic remodeling and endothelial plasticity during network elaboration. These transcriptional responses are mechanistically concordant with the intrinsic molecular composition of EggMA, which retains multiple proteins and mapped human orthologs with angiogenesis-associated relevance (Fig. S1), including ovotransferrin-, clusterin-, fibronectin-, and collagen-related components, together with pathway enrichment in VEGF signaling, ECM organization, non-integrin membrane–ECM interactions, and vascular wall-associated cell surface interactions (Fig S2). Together, these results indicate that the ovoprotein microgel microenvironment is not merely permissive for vascular outgrowth, but actively bioinstructive, coupling its microporous physical architecture with an intrinsically pro-angiogenic protein milieu to promote endothelial invasion, vessel assembly, and progressive microvascular network maturation.

Beyond vasculogenesis, host-derived angiogenic invasion is a prerequisite for effective tissue integration and regenerative performance (*50, 51*). We therefore interrogated the pro-angiogenic capacity of ovoprotein microgel assemblies using the *in ovo* chorioallantoic membrane (CAM) assay (Fig. S28A). After 7 days of implantation, gross imaging revealed pronounced host vascular recruitment toward the implanted constructs, with vessels infiltrating and ramifying throughout the scaffold interior, particularly in the microgel-based groups (Fig. S28B), whereas vascular ingrowth into bulk E-HG remained comparatively limited, consistent with its lower intrinsic porosity relative to the microgel-based matrices. Corresponding histological assessment further confirmed vessel infiltration throughout the E-MG and E-HG@MG scaffolds, with vascular structures traversing the interstitial spaces between adjacent microgels, indicating that the granular architecture provides permissive conduits for host vascular ingrowth (Figs. S28C-D). Moreover, qualitative and quantitative lectin expression analyses further delineated the spatial distribution of host vasculature (Fig. S29), demonstrating that unlike bulk E-HG where vascular signal remained largely restricted to the construct periphery, E-MG and E-HG@MG supported deeper vascular penetration and more pervasive vessel engagement within the scaffold interior (Figs. S29A2-A3). In particular, low-packing E-MG displayed more pronounced vessel extension along microgel interfaces and within interparticle void spaces, underscoring the importance of microporous topology in directing angiogenic invasion. Collectively, these findings establish ovoprotein microgel assemblies as pro-vascular biomaterial systems that not only support endothelial self-organization *in vitro* but also actively promote host-driven angiogenic infiltration *in ovo*, thereby enabling vascular integration across multiple biological contexts.

### 6. Ovoprotein Microgel Assemblies Promote Spheroid-Mediated Biomineralization

Biomineralization is a defining process in bone development and regeneration, governing the deposition of mineralized ECM required for mechanical competence and long-term tissue function (*52, 53*). Protein-based biomaterials are particularly attractive for this purpose because their intrinsic bioactivity and functional group diversity can regulate stem-cell behavior, matrix remodeling, and mineral deposition (*7, 54*). Accordingly, ovoprotein microgel assemblies were investigated as a bioactive and microporous platform for human bone marrow mesenchymal stem cell (hMSC) spheroid-mediated osteogenesis and progressive biomineralization. As illustrated in Fig. 5A, hMSC spheroids were spatially patterned within ovoprotein microgel matrices via AAB, enabling flexible and scalable fabrication of osteogenic spheroid–microgel constructs. These constructs were subsequently cultured and systematically characterized over a defined temporal framework (Fig. 5B) to interrogate how the microgel-based environment regulates hMSC behavior and progressive biomineralization. Notably, hMSC spheroids within microgel matrices underwent sustained cellular egress into the surrounding granular matrix, with progressive cell spreading, microgel-to-microgel bridging, and expansion of a construct-spanning multicellular network over culture (Figs. 5C-D). This process progressed into pronounced construct-scale colonization by Days 21–28, at which stage the initially discrete spheroid-derived outgrowths had converged into a dense and spatially continuous multicellular network permeating porous E-MG constructs. Such progressive occupancy of the interstitial microgel space indicates that the ovoprotein granular matrix supports not only spheroid retention and outgrowth, but also cell-mediated bioassembly, whereby multicellular networks propagate through the microporous architecture and further integrate the interparticle crosslinked constructs. Therefore, ovoprotein microgels serve as both structural building blocks and permissive substrates for tissue-level consolidation, enabling the transition from a particle-assembled scaffold to a biologically unified, cell-remodeled construct.

**Figure 5.**
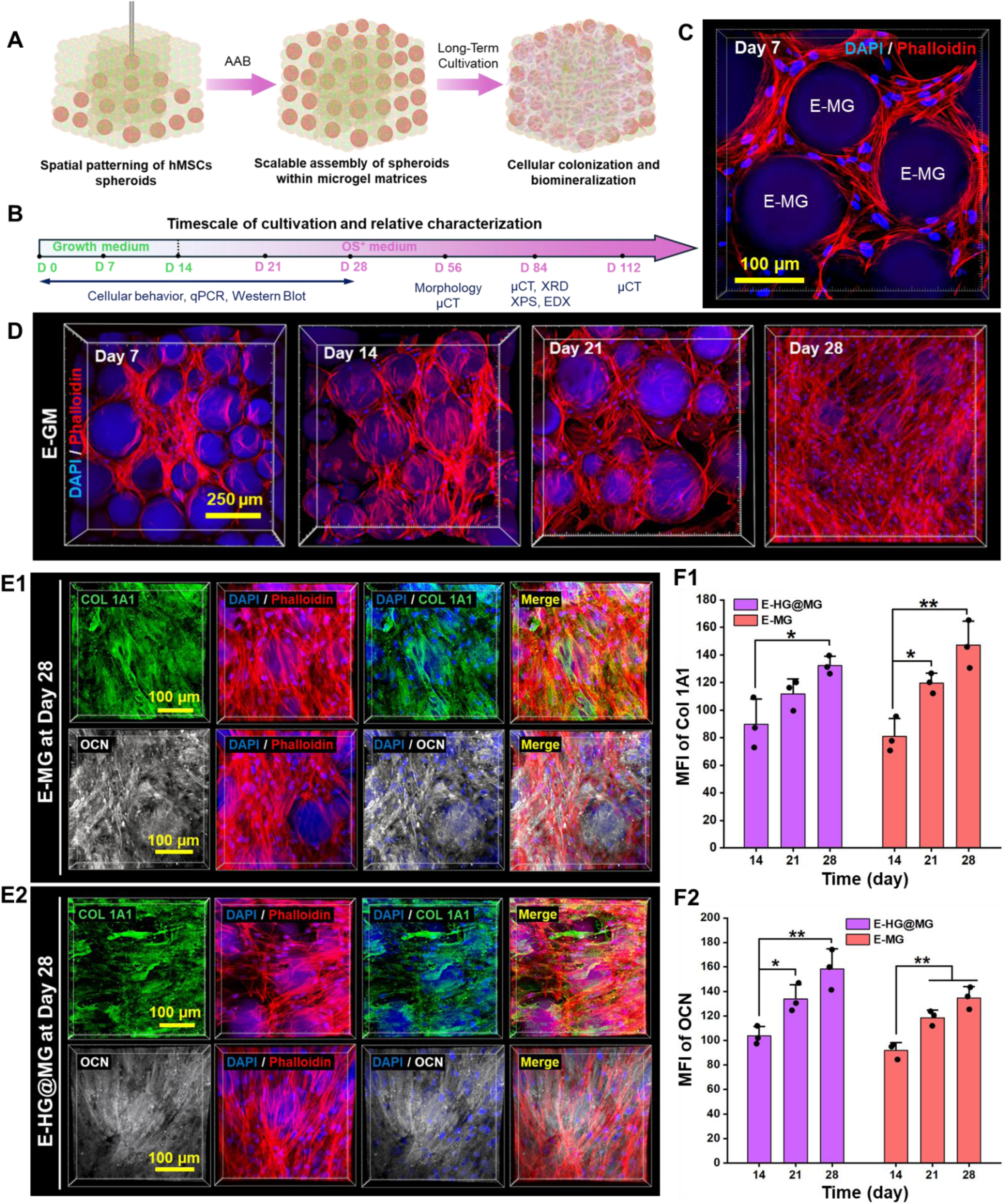
Scalable bioassembly of ovoprotein microgel–spheroid constructs enables osteogenic colonization and matrix elaboration. A, Schematic of AAB-mediated spatial patterning of hMSC spheroids within E-MG matrices for scalable assembly of spheroid–microgel constructs and long-term osteogenesis. **B,** Culture and characterization timeline for assessing cellular colonization, osteogenic differentiation and biomineralization. **C,** Representative high-resolution confocal image of the spheroid–microgel interface at Day 7, showing hMSCs stained for nuclei (DAPI, blue) and F-actin (phalloidin, red), together with surrounding ovoprotein microgels visualized by intrinsic autofluorescence, highlighting cell–microgel interactions within the bioassembled construct. **D,** Time-resolved confocal images showing spheroid-derived cellular egress, intermicrogel migration and progressive construct-wide colonization within E-MG matrices from Days 7 to 28. **E1-E2,** Immunofluorescence images of COL1A1 (green) and OCN (white) deposition in E-MG (**E1**) and E-HG@MG (**E2**) constructs at Day 28, with DAPI-labelled nuclei (blue) and phalloidin-labelled F-actin (red). **F1-F2,** Quantification of COL1A1 (**F1**) and OCN (**F2**) fluorescence intensity over culture. Data were presented as mean ± SD (*n* = 3); *\*P* < 0.05 and *\*\*P* < 0.01.

Concomitantly, osteogenic matrix elaboration progressed throughout the culture, as evidenced by the temporal accumulation of both early and late bone-associated markers within the constructs. Immunostaining revealed progressively intensified COL1A1 deposition from Days 14 to 28 in both E-MG and E-HG@MG, consistent with sustained collagenous matrix assembly during osteogenic maturation (Figs. S30–S31). In parallel, osteocalcin (OCN) deposition became increasingly pronounced over time (Figs. S32–S34), indicating progressive transition toward a more mature mineralizing phenotype. By Day 28, both constructs exhibited abundant COL1A1^+^ and OCN^+^ ECM throughout the cellularized regions (Figs. 5E1-E2), while longer-term culture further confirmed continued OCN-rich matrix accumulation in E-MG at Day 56 (Fig. S34). Quantitative analysis corroborated these trends, showing time-dependent increase in fluorescence intensities of both COL1A1 and OCN in E-MG and E-HG@MG (Figs. 5F1-F2). Furthermore, these phenotypic trends were substantiated at the transcriptional level by a coordinated upregulation of genes associated with osteogenic commitment, matrix production, remodeling, and mineralization (Fig. 6A). Relative to free-standing hMSC spheroid controls, E-MG and E-HG@MG elicited distinct but convergent osteogenic trajectories (Fig. 6A1). In E-MG, *COL1A1* expression increased rapidly from Day 1 and remained elevated through Days 14–28 (Fig. 6A2), consistent with early and sustained collagenous matrix elaboration and tissue consolidation, in agreement with the progressive COL1A1 deposition observed by immunostaining (Figs. 5E–F). This matrix-rich program was accompanied by progressive *RUNX2/SP7* activation and later enrichment of *BGLAP*, *DMP1*, and *PHEX*, indicative of maturation toward a mineralization-competent phenotype. In contrast, E-HG@MG exhibited a comparatively stronger late-stage remodeling response, marked by more pronounced *MMP2* and *MMP9* upregulation at Days 14–28 (Figs. 6A3-A4), suggesting active matrix turnover during prolonged osteogenesis. Persistent *COL5A1/COL6A1* expression in both groups further supports ongoing fibrillar and pericellular matrix organization, whereas late *SMPD3* enrichment is consistent with advancing skeletal maturation. Together, these data indicate that E-MG preferentially supports an early, matrix-deposition–dominant osteogenic program, whereas E-HG@MG promotes a more remodeling-intensive osteogenic microenvironment at later stages. This divergence is likely rooted in the distinct structural milieus experienced by the spheroids within the two matrices. In the hybrid E-HG@MG network, containing a continuous hydrogel phase, presents a comparatively constrained microenvironment that limits extensive cell migration and construct-wide colonization while favoring progression toward late-stage ECM remodeling; by contrast, the interparticle-crosslinked, highly porous E-MG scaffold provides a more permissive granular niche for cellular spreading, proliferation, migration, and spatial colonization, thereby sustaining a matrix-deposition–dominant phase before advanced remodeling.

**Figure 6.**
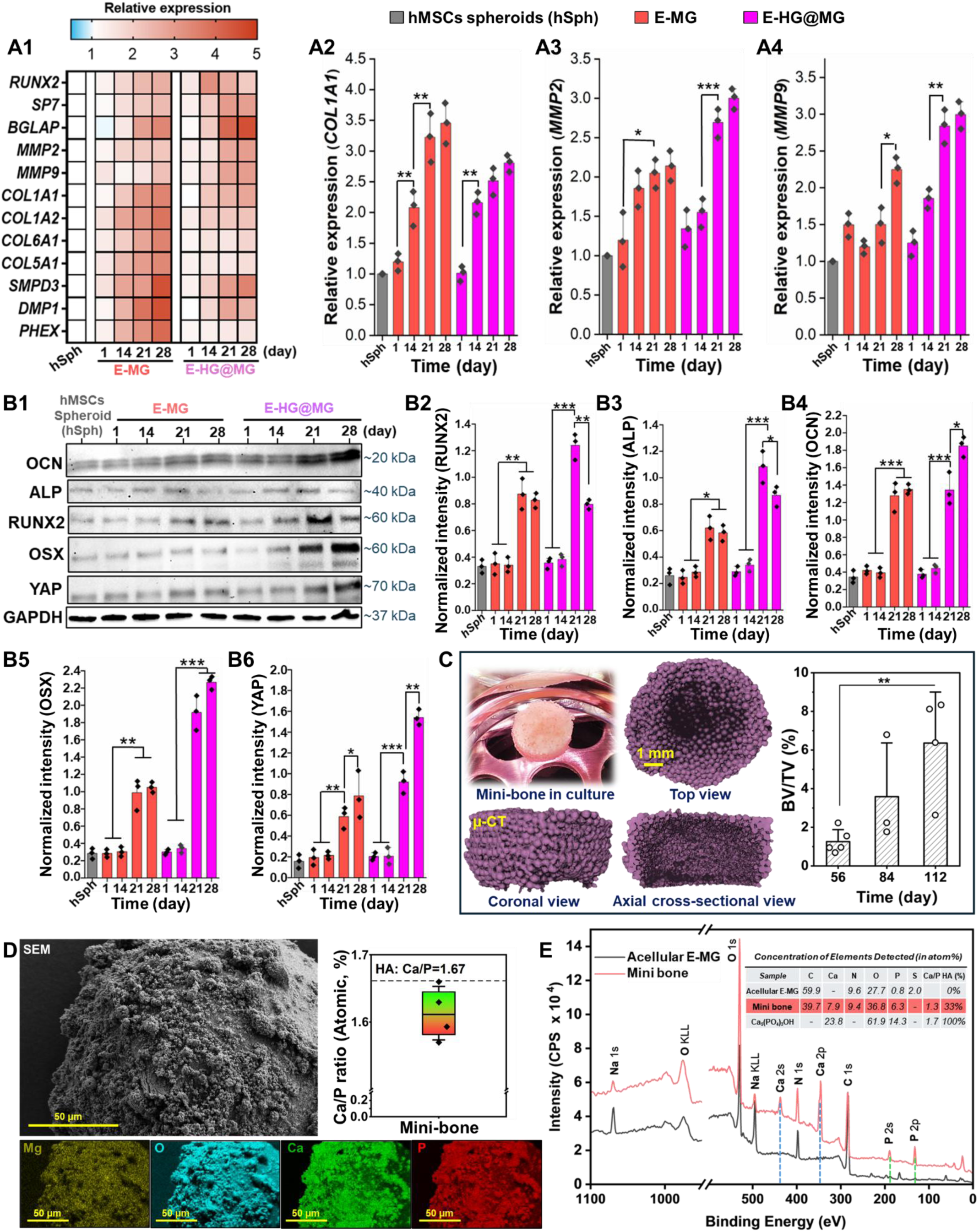
Ovoprotein microgel–spheroid constructs drive osteogenic maturation and multiscale biomineralization. A1-A4, Heatmap and qPCR analysis of osteogenesis-, matrix-remodelling- and mineralization-associated gene expression in hMSC spheroids within E-MG and E-HG@MG constructs during culture. **A1,** Heatmap showing temporal expression profiles of osteogenic differentiation, ECM deposition, matrix remodelling and mineralization-related genes. Relative expression of representative markers, including *COL1A1* (**A2**), *MMP2* (**A3**) and *MMP9* (**A4**). Data were presented as mean ± SD (*n*=3); *\*P* < 0.05, *\*\*P* < 0.01 and *\*\*\*P* < 0.001. **B1-B6,** Western blot analysis of osteogenic and mechanotransductive protein expression in hMSC spheroid-loaded E-MG and E-HG@MG constructs during culture. **B1,** Representative immunoblots for OCN, ALP, RUNX2, OSX and YAP, with GAPDH as the loading control. Densitometric quantification of RUNX2 (**B2**), ALP (**B3**), OCN (**B4**), OSX (**B5**) and YAP (**B6**) expression normalized to GAPDH. Data were presented as mean ± SD (*n*=3); *\*P* < 0.05, *\*\*P* < 0.01 and *\*\*\*P* < 0.001. **C,** Long-term culture of bioprinted mini-bones, including a representative gross image at Day 112 and μCT-based 3D reconstructions of mineralized tissue rendered in purple. Mineralized regions were segmented using a grayscale threshold of 61–225, revealing progressive mineral tissue formation over time. Quantitative analysis of bone volume fraction (BV/TV, %) further confirmed the temporal increase in mineralized tissue content. Data were presented as mean ± SD (*n*≥3); *\*\*P* < 0.01. **D,** SEM imaging and elemental mapping of mineralized mini-bone constructs after 84 days of culture, revealing mineral-rich surface deposits and localized enrichment of Mg, O, Ca and P within the microgel architecture. The measured Ca/P atomic ratio approached that of hydroxyapatite. Data were presented as mean ± SD (*n*=4). **E,** XPS spectra and elemental composition analysis of spheroid-laden mini-bones after 84 days of culture compared with cell-free E-MG matrices, showing enriched Ca 2p and P 2p signals consistent with bone-like mineral formation.

These matrix-dependent osteogenic trajectories were further substantiated at the protein level by western blot analysis of lineage- and function-associated markers (Fig. 6B1). Both microgel systems elicited time-dependent increases in OCN, ALP, RUNX2, and OSX (Figs. 6B2-B5), confirming progressive osteogenic maturation, while the concurrent elevation of YAP indicates sustained mechanotransductive engagement with the evolving microenvironment (Fig. 6B6). Notably, E-HG@MG exhibited more pronounced late-stage expression of RUNX2, ALP, OCN, OSX, and YAP, consistent with a remodeling-intensive and maturation-oriented osteogenic program, whereas E-MG maintained a stronger matrix-elaborative phenotype, concordant with the transcriptional signatures and higher COL1A1 and OCN deposition observed in Fig. 5F and Fig. 6A. This divergence is mechanistically concordant with the distinct structural milieus imposed by the two matrices and further aligns with the intrinsic molecular composition of EggMA. Proteomic mapping identified retained ovoprotein constituents and human ortholog-associated functions with osteoregulatory relevance, including ovotransferrin/transferrin-related proteins, fibronectin-associated proteins, and collagen/fibrillar matrix-associated proteins (Fig. S1A). Such components are not merely structural: ovotransferrin has been shown to promote bone formation (*55*), whereas fibronectin- and collagen-enriched matrices potentiate osteogenic commitment through integrin-dependent adhesion, MAPK/ERK signaling, and RUNX2-associated transcription (*56–58*). Consistently, pathway enrichment highlighted ECM organization, non-integrin membrane–ECM interactions, receptor tyrosine kinase signaling, MAPK signaling, and broader tissue-development programs (Fig. S2). These findings suggest that the ovoprotein-based matrix provides a bioinstructive protein milieu that couples adhesion, mechanotransduction, matrix remodeling, and mineralization during spheroid-mediated osteogenesis.

Building upon these findings, bioprinted osteogenic constructs were maintained in long-term osteogenic culture for up to 112 days to interrogate biomineralization across multiple structural scales. Progressive densification of the constructs was evident by micro computed tomography (μCT), which revealed time-dependent increases in mineralized tissue volume together with marked expansion of the mineralized area from Day 56 to 112 (Fig. 6C, Fig. S36, and Supplementary Movie 6). The resulting “mini-bone” constructs exhibited increasingly consolidated internal architecture in top, coronal, and axial views, consistent with sustained mineral accrual and volumetric maturation over prolonged culture. Notably, mineral deposition was initially concentrated in the construct periphery and progressively advanced inward with time, whereas only discrete mineralized particulates were observed in the central region. This spatial pattern likely reflects diffusion limitations under static culture, where nutrient and osteogenic factor availability were greater at the construct surface, while the central inorganic core might correspond to the originally bioprinted spheroid-rich domain. This underscores the importance of incorporating dynamic perfusion culture, even in highly porous microgel constructs, to enhance mass transport and promote more homogeneous mineral maturation throughout the fabricated mini-bones. Quantitative analyses of both BV/TV (%, Fig. 6C) and mineralized volume fraction (Fig. S36B) further confirmed significant time-dependent mineral accumulation, indicating that the spheroid–microgel system supports sustained osteogenic progression beyond early-stage matrix deposition. At the microscale, SEM imaging of mini-bones after 84 days of culture revealed dense mineralized deposits coating and bridging the construct surface, while elemental mapping demonstrated marked enrichment of calcium and phosphorus throughout the matured tissues (Fig. 6D), in clear contrast to the acellular microgel controls, which lacked comparable mineral-associated elemental signatures (Fig. S37). Importantly, the measured Ca/P atomic ratio (1.61 ± 0.04) approached that of hydroxyapatite (Ca/P = 1.67) (*59*), supporting the formation of a bone-relevant apatite-like mineral phase rather than nonspecific ionic precipitation. This interpretation was further substantiated by X-ray photoelectron spectroscopy (XPS), which showed clear emergence and enrichment of Ca 2p and P 2p signals in the mini-bone (Day 84) relative to acellular E-MG controls (Fig. 6E), together with elemental compositions consistent with biologically mineralized matrices. X-ray diffraction (XRD) analysis further identified crystalline phases assignable to hydroxyapatite-related calcium phosphate and calcium magnesium phosphate species, confirming progressive mineral crystallization within the constructs (Fig. S38). Mechanistically, these mineralization outcomes are consistent with both the intrinsic biochemical composition of EggMA and the established role of proteinaceous matrices in biomineralization. Proteins can nucleate and organize mineral growth by concentrating ions, providing interfacial binding motifs, and stabilizing growing crystal phases, while matrix proteins and cell-secreted osteogenic ECM further direct crystal maturation (*60–62*). Moreover, the retained ovoprotein components (Fig. S1), including ovotransferrin/ transferrin-related proteins and fibronectin- and collagen-associated functions, are particularly relevant. For instance, ovotransferrin has been reported to stimulate osteoblast proliferation and differentiation, whereas fibronectin- and collagen-rich matrices are well known to potentiate osteogenic commitment and matrix mineralization through adhesion-mediated signaling and organized ECM assembly (*55, 63, 64*). Collectively, these multiscale cellular, molecular, and mineralization outcomes establish ovoprotein microgel assemblies as a bioinstructive matrix that synergizes with hMSC spheroids to drive progressive biomineralization. By integrating a protein-rich, osteoregulatory microenvironment with a microporous and structurally adaptive granular architecture, this system supports osteogenic maturation, matrix remodeling, and long-term bone-like tissue formation.

## Conclusion and Outlook

In summary, we developed a photoresponsive ovoprotein-derived biomaterial platform that integrates bulk hydrogels, orthogonally crosslinked microgel assemblies, and hybrid bioinks for scalable tissue biofabrication. Methacrylation of EW-derived ovoproteins yielded EggMA, a protein-rich matrix that retained intrinsic biochemical complexity while enabling visible-light crosslinking, tunable mechanics, and concentration-dependent regulation of cell behavior, with its proteomic composition and associated biological functions systematically delineated. We further established an orthogonal crosslinking strategy to generate ovoprotein microgel assemblies through interparticle dityrosine coupling, endowing the system with structural robustness, microporous architecture, and intrinsic autofluorescence for label-free optical interrogation. These properties collectively enabled broad compatibility across DLP, EBB, and AAB bioprinting modalities, thereby expanding its utility in tissue biofabrication. More broadly, the abundance and low cost of ovoproteins position this platform as a fertile basis for future advances in biomaterials. Promising directions include the development of photocrosslinkable ovoprotein bioinks directly driven by dityrosine coupling, the design of mechanically reinforced ovoprotein hydrogels through tunable dual-network architectures, the extension of these materials to light-based and particularly volumetric bioprinting for rapid and scalable biofabrication (*65, 66*), and the exploitation of their intrinsic autofluorescence for label-free detection and monitoring of material stability, degradation, tissue maturation, and other applications (*32*).

Beyond processability, the ovoprotein microgel-based environment actively modulated key regenerative processes. The microgel-based matrices attenuated sustained pro-inflammatory macrophage activation *in vitro*, promoted endothelial organization and host angiogenic infiltration, and enabled precise spheroid-based assembly of vascularized constructs with progressive lumenization, branching, and inosculation. Moreover, when combined with osteogenic spheroids, the ovoprotein microenvironment supported extensive cellular colonization, osteogenic matrix elaboration, coordinated activation of osteogenic and remodeling-associated programs, and long-term biomineral maturation toward bone-like mineralized tissue. Collectively, these findings position ovoprotein microgel assemblies as an instructive and structurally adaptive biomaterial platform that couples intrinsic protein function with modular biofabrication. To meet the growing demand for multifunctional and scalable biofabrication, future work should extend this strategy toward perfusion-enabled culture, multicellular interface engineering, and *in vivo* regenerative validation.

## Supporting information

Description of Additional Supplementary Movies

## Acknowledgements

The authors acknowledge support from the National Institute of Biomedical Imaging and Bioengineering (NIBIB) awards R01EB034566 (I.T.O.) and R01EB036245 (I.T.O.), and the National Institute of Dental and Craniofacial Research (NIDCR) award R01DE0335200 (I.T.O.). The authors thank Mr. Logan Haugh (Penn State) for assistance in microgel preparation and Ms. Ilayda Namli (Penn State) for assistance with multiphoton microscopy. The authors acknowledge Dr. Ethan Gerhard from the Department of Biomedical Engineering at Penn State for assistance with photoluminescence spectroscopy, and Dr. Thomas Neuberger and Mr. Sean Gullette from the High Field MRI Facility at Penn State for support with MRI and Micro-CT measurements. The authors also acknowledge use of the Pennsylvania State University Materials Characterization Lab Core Facility, Materials Research Institute, University Park, PA (RRID:SCR_012386), and thank Dr. Nichole M. Wonderling and Dr. Anthony Raymond Richardella for assistance with XRD measurements, and Dr. Jeff Shallenberger for support with XPS measurements and analysis. The authors further thank Mr. Scott L. Kephart of the Poultry Education and Research Center at Penn State University for kindly providing the chicken eggs used in this work.

## Author contributions

**S.L.** conceptualized the entire study, designed and performed experiments and analyses, and wrote the manuscript. **V.P.** and **J.C.M.** designed and performed the experiments and analyses. **M.D.S.**, **D.G.**, **M.Y.**, **V.S.**, and **Y.O.Y.** conducted experiments and analyses. **I.T.O.** conceptualized and supervised the study and acquired funding. All authors reviewed and approved the manuscript.

## Data availability

All data supporting the results of this study are available within the Article and its Supplementary Information. Additional data are available from the corresponding author upon request.

## 4. Materials and methods

### 4.1 Synthesis of the Ovoprotein-Based Photoresponsive Hydrogel

Fresh organic chicken eggs (*Gallus gallus* domesticus, Hy-Line W-36 strain; Hy-Line International, USA) were kindly provided by Poultry Education and Research Center at PSU (University Park, PA, USA). The eggs were rinsed with deionized water and surface-sterilized with 70% ethanol. The Ovoprotein, eggwhite (EW, also referred to as ovalbumin) was carefully separated under sterile conditions and gently stirred to obtain a homogeneous mixture. Subsequently, EW was functionalized through reaction with methacrylic anhydride (MA, ≥94%, Sigma-Aldrich) to confer photoresponsive functionality. In brief, 50 mL of EW was mixed with 50 mL of Dulbecco’s phosphate-buffered saline (DPBS; Corning, NY) and stirred at 4 °C for 1 h. The mixture was centrifuged at 8,000 rpm for 5 min to eliminate air bubbles and sediment solid amyloid-like fibrils, commonly existed in eggwhite. The resulting supernatant was collected and adjusted to pH 9.0 using 1 M NaOH. Afterwords, 2.5 mL of MA was added incrementally in six portions, and the reaction was carried out under continuous stirring at 4 °C for 5 h, with pH rigorously maintained at 9 throughout the process. Following the reaction, the mixture was centrifuged at 8,000 rpm for 5 min, and the supernatant was collected, dialyzed using a 6–8 kDa cut-off membrane, and freeze-dried. The resulting functionalized eggwhite protein (designated EggMA) powder was sterilized using an ethylene oxide gas sterilizer (Anprolene AN74i, Andersen Products, USA) and stored at 4 °C for subsequent use and long-term preservation.

### 4.2. Chemical Characterization of EggMA

To verify the successful methacrylation of eggwhite proteins, proton nuclear magnetic resonance (^1^H NMR) spectroscopy was conducted. The lyophilized EggMA powder was dissolved in Dimethyl sulfoxide-d6 (DMSO; 6%, w/v) at 22 °C and transferred into an NMR tube. Spectra were acquired using a Bruker AVIII-HD-500 spectrometer (Bruker, Billerica, MA, USA) equipped with a Bruker Ascend 500 MHz magnet. The lyophilized unmodified EW protein served as the control. Methacrylation was assessed by comparing the spectra of native EW and EggMA, focusing on changes in the characteristic proton signals associated with lysine residues and the appearance of methacrylate-associated vinyl proton signals. Photocrosslinked EggMA was additionally analyzed and compared with uncrosslinked EggMA to evaluate consumption of vinyl groups during network formation.

Changes in free primary amine groups following methacrylation were further assessed using a 2,4,6-trinitrobenzenesulfonic acid (TNBS) assay. Briefly, native EW and EggMA were separately dissolved in 0.1 M sodium bicarbonate buffer at a concentration of 5 mg/mL (w/v). For each sample, 0.5 mL of protein solution was mixed with 0.5 mL of 0.1% (w/v) TNBS solution and incubated for 2 h. The reaction was quenched by adding 0.25 mL of 1 M hydrochloric acid (HCl) and 0.5 mL of sodium dodecyl sulfate (SDS, 10%, w/v) solution. Absorbance was subsequently measured at 335 nm using a microplate reader (Infinite 200 PRO, Tecan Group Ltd., Switzerland).

### 4.3. Proteomic analysis of EggMA

Protein extracts from EW and EggMA were prepared by reducing with dithiothreitol (DTT, 10 mM, 30 min, 56 °C), alkylating with iodoacetamide (55 mM, 30 min, dark), and digesting overnight with sequencing-grade trypsin (1:50, w/w) at 37 °C. All reagents were sourced from Thermo Pierce, USA. Peptides were separated on an EASY-nLC system and introduced into a Thermo-Scientific Orbitrap Eclipse Tribrid mass spectrometer operating in data-dependent acquisition (DDA) mode using a modified Universal Method. Full MS1 scans were acquired in the Orbitrap at 120,000 resolution over an m/z 150–1500 range, with a precursor tolerance of ±10 ppm. The normalized AGC target was 250%, and only precursor ions with charge states 2–6 were selected for fragmentation. Dynamic exclusion was set to 60 s to minimize repeated sampling. Spectra were searched using Sequest HT in Proteome Discoverer 2.5 (Thermo-Fisher Scientific) against the *Gallus gallus* UniProt database with a reverse decoy approach. Search parameters included trypsin specificity (≤ 2 missed cleavages), carbamidomethylation (C, fixed), oxidation (M, variable), and methacrylate adducts (+68 Da on Lys, variable). Precursor and fragment tolerances were set to ± 10 ppm and ± 0.02 Da, respectively. Protein identifications were filtered at < 1% and *p*-values for protein abundance ratios were computed using a background-based t-test and adjusted for multiple comparisons using the Benjamini–Hochberg method to control the false discovery rate (FDR). Quantification was performed by spectral counting (PSM-based quantitation), and normalized PSM values were compared between native and methacrylated groups. Only proteins with ≥ 2 unique peptides were considered. Identified *Gallus gallus* proteins were mapped to human orthologs using UniProt and Reactome annotations, and pathway enrichment was performed using ReactomePA and clusterProfiler packages in R.

### 4.4. Characterization of the Photoresponsive Behavior of EggMA

EggMA hydrogel precursor solutions with concentrations of 5, 6, 10, 15, and 20% (w/v) were prepared by dissolving the appropriate amount of EggMA powder into DPBS. The photoinitiator lithium phenyl(2,4,6-trimethylbenzoyl)phosphinate (LAP, ≥95%, Sigma-Aldrich) was added at a final concentration of 5 mg/mL under light-protected conditions to prevent premature photoinitiation. The photocrosslinking behavior of EggMA hydrogels were determined using a rheometer (MCR 302, Anton Paar, Austria) equipped with a 405 nm ultraviolet (UV) light source. All measurements were carried out on three replicate samples using a 25-mm parallel-plate configuration, with the temperature maintained at 22 °C through a Peltier-controlled system. A time sweep test was conducted in which the samples were preconditioned for 3 min at 22 °C, followed by exposure to 405 nm UV light for 2 min to initiate photopolymerization. The evolution of *G′* and *G″* was continuously recorded over time. Subsequently, a frequency sweep test was carried out to assess the viscoelastic behavior of the crosslinked EggMA hydrogels, applying a constant strain amplitude of 1% while varying the angular frequency from 0.1 to 100 rad/s. Moreover, the compressive mechanical properties of crosslinked EggMA hydrogel (cylindrical specimens, 7.5 mm in diameter and 1.8 mm in height) were evaluated using the same rheometer under a uniaxial compression mode at a constant compression rate of 0.01 mm/s. The stress–strain curves were recorded, and the compressive modulus was calculated from the linear region of the curve. Four replicates were tested for each group.

To visually assess the photocrosslinking behavior and gel formation, 1 mL of the prepared EggMA precursor solutions was transferred into transparent borosilicate glass reagent vials with screw caps. The samples were irradiated with 405 nm UV light for 2 min to initiate photopolymerization. After exposure, the vials were inverted to qualitatively evaluate hydrogel formation and flow resistance. Representative images of EggMA formulations were captured before and after irradiation to visually confirm the sol–gel transition induced by photocrosslinking.

To evaluate the intrinsic autofluorescence of EggMA hydrogels, precursor solutions at 6, 10, and 15% (w/v) EggMA containing 5 mg/mL LAP were cast into the polylactic acid (PLA) molds (10 mm × 5 mm × 2 mm) fabricated using a fused deposition modelling (FDM) printer (Bambu Lab P1S, Bambu Lab, USA), and photopolymerized under 405 nm light for 2 min. The photoluminescence spectra of the crosslinked hydrogels were measured using a FluoroMax-4 spectrofluorometer (Horiba Scientific, Edison, NJ) equipped with a solid-state sample holder. Excitation and emission scans were recorded with a 10-nm step interval to assess fluorescence intensity and spectral profiles. To further assess the influence of Ru/SPS-mediated secondary crosslinking on fluorescence behavior, the EggMA hydrogels were subjected to dityrosine crosslinking as described in Section 4.8 and subsequently analyzed. In parallel, photocrosslinked EggMA hydrogels prepared with Ru/SPS premixed in the precursor solution, and uncrosslinked EggMA hydrogel solution without LAP and Ru/SPS, were included as control groups.

### 4.5. Cytocompatibility and Cellular Response of EggMA

GFP^+^ and GFP^-^ MDA-MB-231 metastatic breast cancer cell line (passage 8–15, were kindly provided by Dr. Danny Welch, from the University of Kansas, Kansas City, KS) and NHLFs (passage 4-8, purchased from Lonza in Houston, Texas) were used. Both cell lines were cultured in Dulbecco’s Modified Eagle Medium (DMEM, Sigma-Aldrich) medium supplemented with 10% (v/v) fetal bovine serum (FBS) (Life Technologies, NY, USA), 1 mm Glutamine (Life Technologies), and 1% (v/v) penicillin-streptomycin (P/S, Life Technologies). First, NHLF were seeded onto crosslinked EggMA at varying concentrations (6, 10, 15, and 20%, w/v) with a uniform cell density of 1 × 10⁵ cells/mL. Subsequently, NHLFs were encapsulated within EggMA at a concentration of 1 × 10⁶ cells/mL. The resulting NHLFs-laden EggMA bioinks were extruded into well plates using an INKREDIBLE+ bioprinter (CELLINK, Sweden) and photo-crosslinked under 405 nm UV light for 2 min. Following the same procedure, GFP^+^ MDA-MB-231 cells were encapsulated within EggMA hydrogels at a concentration of 3.5 × 10⁶ cells/mL to fabricate GFP^+^ MDA-MB-231-laden EggMA constructs. The constructs were cultured at 37 °C in a humidified atmosphere containing 5% CO₂, and cell viability and morphology were assessed at designated time points. Moreover, proliferation of NHLFs and GFP^-^ MDA-MB-231 cells on EggMA scaffolds were evaluated using a Cell Counting Kit-8 assay (CCK-8, Sigma-Aldrich) according to the manufacturer’s protocol, as described previously. In short, 60 µL of EggMA hydrogels were coated and photo-crosslinked within 96-well plates, followed by seeding 5 × 10^3^ cells per well and cultured for 1, 3, and 5 days. Cells cultured in 2D without EggMA served as a control group. After incubation with 10 µL of CCK-8 reagent for 2 h, the absorbance of each well was measured to evaluate the metabolic activity of cells using an Infinite 200 PRO microplate reader at a wavelength of 450 nm.

### 4.6. Fabrication and Characterization of EggMA Microgels

*Fabrication of EggMA Microgels:* EggMA microgels were fabricated using an in-house developed T-junction coaxial microfluidic system, as described previously (*30, 67, 68*). In brief, the setup consisted of an orthogonal flow-focusing nozzle comprising a 15G outer needle (inner diameter:1370 µm) for the continuous oil phase and a 21G inner needle (inner diameter: 510 µm) for the dispersed aqueous phase. Two programmable syringe pumps (1000-US/SyringeONE, New Era Pump System, NY) were independently connected to the inner and outer needles of the coaxial microfluidic device to enable automated and controlled extrusion. The continuous phase consisted of light mineral oil containing 4% (v/v) Span-80, which was sterilized using a 0.22-µm filter system (Cell Treat, China). The dispersed aqueous phase consisted of either cell-laden or acellular EggMA hydrogel (bio)inks. Microdroplets were generated through interfacial shear between the continuous oil phase and the dispersed aqueous phase, followed by photo-crosslinking under 405 nm light. The crosslinked microgels were subsequently collected into 50 mL centrifuge tubes and washed three times with DPBS to remove residual oil and surfactant before further processing.

*Optimization and Characterization of EggMA Microgels:* 10 and 15% (w/v) EggMA hydrogel precursor solutions were employed for microgel fabrication, guided by the physicochemical and cellular characterization results presented in Sections 4.4 and 4.5. A one-factor-at-a-time experimental design was employed to systematically investigate the effects of oil and EggMA aqueous phase flow rates on microgel size and morphology, with oil flow rates of 250, 500, and 1000 μL/min and EggMA flow rates of 5, 10, 20, 50, and 100 μL/min. The microdroplets generated were photocrosslinked under identical conditions, and the resulting microgels were washed prior to further characterization. Microgel morphology was examined using optical microscopy (EVOS® XL Core, Fisher Scientific) and fluorescence microscopy (EVOS® FL Auto, Fisher Scientific) with 405 nm excitation. The size distribution and circularity of individual microgels were analyzed using ImageJ software (Fiji, National Institutes of Health, NIH), with over 100 microgels measured for each group. The dependence of microgel diameter on the flow rates of the oil and EggMA phases was evaluated using Origin 2025 software (OriginLab Corporation, Northampton, MA, USA).

*Cellular Response of EggMA Microgels:* GFP^+^ MDA-MB-231 and NHLFs were separately seeded onto 10% (w/v) EggMA microgels at a controlled density of 1 × 10⁶ cells/mL. Briefly, 100 μL of prepared microgels were loaded into 48-well suspension culture plates, followed by the addition of 200 μL of cell suspension per well for cell adhesion. Separately, GFP^+^ MDA-MB-231-laden EggMA microgel (10%, w/v) were prepared at a cell density of 3.5 × 10⁶ cells/mL. All samples were maintained under standard culture conditions. Cellular viability and morphology were assessed using an LSM 880 Zeiss confocal microscope (Thornwood, NY) with LIVE/DEAD and nuclear/cytoskeleton staining, employing Z-stack imaging for 3D analysis. The extent of cell coverage on the microgel surface was evaluated and quantified using ImageJ based on the acquired fluorescent images.

### 4.7. Preparation and Characterization of Microgel-based (Bio)inks

*Preparation of EggMA Microgel-based (Bio)inks:* 1mM Tris(2,2′-bipyridyl)dichlororuthenium(II) (Ru, Sigma-Aldrich) and 10mM sodium persulfate (SPS, ≥ 98%, Sigma-Aldrich) solutions were freshly combined at a 1:1 (v/v) ratio to generate the Ru/SPS photoinitiator system. The Ru/SPS solution was promptly added into the prepared cell-laden or acellular EggMA microgels at a 1:1 (v/v) volume ratio, gently mixed by pipetting to ensure uniform distribution, and incubated for 5 min at room temperature (RT). Afterwards, the mixture was centrifuged, and the supernatant was carefully removed to obtain only EggMA microgel-based (designated as E-MG) (bio)inks for subsequent bioprinting or use as a supporting matrix. Two distinct microgel compaction levels were prepared by centrifugation at 2,000 rpm (low packing) and 13,000 rpm (high packing) for 5 min. In addition, the prepared cell-laden or acellular EggMA microgels were gently mixed with the EggMA bulk hydrogel (10% w/v, containing 5 mg/mL LAP) at a 1:1 (v/v) volume ratio, followed by centrifugation at 2,000 rpm for 5 min. The supernatant was then removed to obtain the EggMA bulk hydrogel–microgel hybrid (designated as E-HG@MG) bioinks. The EggMA bulk hydrogel (10% w/v) with 5 mg/mL LAP (designated as E-HG) server as a microgel-free control group.

*Rheological Behavior of EggMA Microgel-based Inks:* Rheological properties of the prepared E-MG (low- and high-packing), E-HG@MG, and E-HG inks were evaluated using Anton Paar rheometer. Briefly, a flow sweep test was performed to characterize their shear-thinning behavior over a shear rate range of 0.01–100 s⁻¹. Subsequently, a recovery sweep test was conducted to evaluate the self-healing capability of these inks after exposure to high shear stress. The test consisted of three consecutive cycles, alternating between a low shear rate of 0.1 s⁻¹ for 60 s and a high shear rate of 100 s⁻¹ for 10 s, during which the relative viscosity was continuously recorded. Moreover, an amplitude sweep test was performed at a fixed angular frequency of 1 rad/s, with the shear strain varied from 10^⁻2^ to 10^3^ %, to determine the linear viscoelastic region (LVR) and the yield point of these inks.

*Dityrosine Crosslinking Behavior:* To assess the dityrosine crosslinking capability of the EW protein and the interparticle crosslinking behavior of EggMA microgels, the dityrosine content was quantitatively evaluated over different crosslinking times. Briefly, the dityrosine crosslinking property of EW protein was first evaluated. Freeze-dried EW powder was dissolved to prepare 6, 10, and 15% (w/v) solutions in DPBS containing 0.05% Ru and 0.5% SPS under light-protected conditions. Aliquots of 100 μL of each solution were added to 96-well plates in quintuplicate and exposed to 405 nm UV light for 0, 10, 30, 60, 120, and 300 s. The fluorescence emission of each sample, corresponding to the dityrosine groups, was measured at 415 nm with excitation at 320 nm using Infinite 200 PRO microplate reader. Similarly, the interparticle crosslinking behavior of microgels was evaluated. Prepared E-MG (10%, w/v) were mixed with freshly prepared Ru/SPS photoinitiator at a 1:1 ratio (v/v) for 5 min as described above, followed by centrifugation at 2,000 rpm for 5 min to remove the supernatant. Aliquots of 100 μL of microgels were then transferred into 96-well plates, centrifuged again to remove air bubbles and promote particle–particle contact, and exposed to 405 nm UV light for 0, 60, 120, 180, 240, and 300 s. Fluorescence emission corresponding to dityrosine formation was measured under the same excitation/emission settings described above.

*Pore Size and Porosity of EggMA Microgel-based Scaffolds:* The pore structure and porosity of the fabricated E-MG scaffolds with low-packing and high-packing were assessed using a fluorescent dye infiltration assay. Briefly, the scaffolds were kept in a 3D-printed mold (7.5 mm in diameter and 1.8 mm in height) with one side covered by a glass slide, and immersed in a 5 mg/mL solution of high-molecular-weight fluorescein isothiocyanate (FITC)-dextran (2 MDa, Sigma-Aldrich) solution. The scaffolds were allowed to infiltrate for 5 min on a shaker to ensure complete filling of all pores. Fluorescent images were then acquired using a multiphoton fluorescence microscopy (Leica SP8 DIVE, Germany). The FITC–dextran within the interstitial pore spaces was visualized using 488 nm excitation, while the intrinsically autofluorescent microgels were imaged using the DAPI channel. 2D porosity and pore architecture were analyzed using ImageJ software (National Institutes of Health, Bethesda, MD, USA), and 3D reconstruction and volumetric porosity analysis were performed using Imaris software (version 9.0, Oxford Instruments, Abingdon, UK).

### 4.8. 3D Bioprinting of EggMA Hydrogel and Microgels

*Digital Light Processing (DLP):* Prepared E-HG (bio)inks were utilized for DLP using a commercial DLP printer (Mars 4 DLP, ELEGOO, China). The PSU Nittany Lion head model (L × W × H =33.5 × 32 × 35 mm) was printed with a layer thickness of 0.1 mm. The printed constructs were then imaged using a 14.1 T micro-MRI system (Bruker, Germany) to reconstruct its 3D geometry following a previously described protocol (*67*). Shape fidelity was assessed by comparing the reconstructed 3D model with respect to the original computer-aided design (CAD) model using CloudCompare (version 2.0, open-source software, France). Surface deviation analysis was visualized as a 3D heat map. Moreover, a cell-laden wave-shaped structure (L × W × H = 12 × 12 × 2 mm), were bioprinted using GFP^+^ MDA-MB-231-laden E-HG (3.5 × 10⁶ cells/mL). The spatial distribution and viability of GFP^+^ MDA-MB-231 cells within printed constructs were subsequently assessed using a LSM 880 Zeiss confocal microscopy.

*Extrusion-based Bioprinting (EBB):* The prepared E-HG, E-MG, and E-HG@MG (bio)inks were applied for EBB using an INKREDIBLE+ bioprinter (CELLINK, Sweden). First, GFP^+^ MDA-MB-231-laden E-HG (3.5 × 10⁶ cells/mL) was applied, a PSU Nittany Lion logo (L × W × H =32 × 24 × 0.4 mm) was bioprinted with a 27G nozzle at an air pressure of 6–15 kPa. The E-HG@MG (bio)inks, where acellular EggMA hydrogel and acellular/GFP^+^ MDA-MB-231-laden microgels (3.5 × 10⁶ cells/mL) were mixed at a 1:1 (v/v) ratio, respectively, were (bio)printed into calycinal tube and branch structures with a 21G nozzle at an air pressure of 30–35 kPa. Moreover, the GFP^+^ MDA-MB-231-laden E-MG with low-packing and acellular E-MG with high-packing were utilized for (bio)printing macroporous gride constructs using a 24G nozzle with an air pressure of 15-35 kPa. After (bio)printing, all samples were crosslinked under 405 UV light for 2 min.

*Aspiration-assistant Bioprinting (AAB):* Prepared E-MG and E-HG@MG were employed as supporting matrices for AAB. Briefly, each of the microgel-based matrix was loaded into a 3D-printed PVA mold (4 mm in diameter, 2 mm in height) as described above, and positioned on the bioprinting platform. Prefabricated spheroids, including vascular and hMSC spheroids, were suspended in DMEM culture medium within a 50 mm Petri dish serving as the spheroid reservoir. An in-house established AAB system was utilized for spheroid manipulation and placement (*37*). The spheroids were aspirated using a 32 G blunt-end nozzle (inner diameter: 110 μm) under a vacuum pressure of 15–20 mmHg, transferred to the bioprinting platform, and precisely deposited into the microgel-based matrices by releasing the vacuum pressure. The spheroids were sequentially patterned within the matrix according to the predefined structural design by repeating the aforementioned procedure. In addition, the bioprinting accuracy within microgels was evaluated by quantifying the positional deviation of deposited spheroids from predefined reference coordinates, as described previously (*69*). For each type of microgel-based supporting matrix, fourteen spheroids were bioprinted, and their positions were quantified from microscopic images using ImageJ. The root mean square deviation (*RMSD*) was calculated using the following equation to quantify the AAB accuracy of spheroids within these supporting matrices:

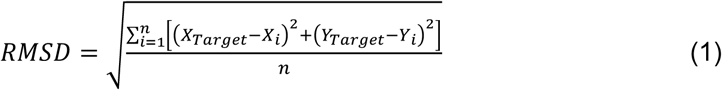

where, *X_Target_* and *Y_Target_* are the *X* and *Y* coordinates of the predefined target position, and *X_i_* and *Y_i_* are the actual X and Y position of bioprinted spheroids respectively; and *n* is the size of tested samples. Lower *RMSD* values indicate higher printing accuracy and reproducibility of the bioprinting process.

### 4.9. Assessment of Inflammatory Immune Response

THP-1 human monocytes were cultured in RPMI-1640 medium supplemented with 10% FBS, 0.05 mM β-mercaptoethanol, and P/S (10 U/mL and 10 μg/mL). The reagents, medium components were sourced from Thermo-Fisher, while the cytokines used were sourced from PeproTech, USA. Differentiation into adherent, resting macrophages (M0) was initiated by treating the suspended monocytes with 160 nM phorbol 12-myristate 13-acetate (PMA) overnight. M0 macrophages were polarized toward either an M1 (pro-inflammatory) phenotype using 20 ng/mL IFN-γ or an M2 (anti-inflammatory) phenotype using 20 ng/mL IL-4 for 72 h (*70*). These populations served as the baseline (control) macrophage conditions. The effect of the material properties to prime the naïve M0 macrophages were tested by encapsulating them in E-HG, E-MG or E-HG@MG in prepolymer samples at a cell density of 6 × 10^6^ cells/mL (*71*) after which the encapsulated samples were crosslinked following the same light-curing regimen mentioned previously in Section 4.5. As control matrices, GelMA was used which was synthesized based on our previously published protocol (*67*) from gelatin (derived from porcine skin, Sigma-Aldrich, USA) (ref). 10 % w/v prepolymer solutions of GelMA-HG or GelMA-HG@MG were encapsulated with the same cell density of 6 × 10^6^ cells/mL. The GelMA-MG group did not form interparticle crosslinking using photocuring regimen followed using LAP and hence they were omitted from the study.

*Quantitative Analysis using Flow Cytometry:* The polarized macrophages (IFN-γ or IL-4 treated) were trypsinized and characterized for the population distribution using flow cytometry. The cells embedded within the constructs were harvested by digesting the gels for 30 min in enzyme cocktail of 1:1 v/v of 0.1 % w/v Collagenase, Type I (Sigma-Aldrich; 125 CDU mg-^1^) and 0.2% w/v Dispase (Gibco, USA; 0.5 U mg^-1^). The harvested cells were blocked with 10 % (v/v) normal donkey serum (Abcam, USA) for 30 min at RT, followed by incubation with sequential incubation of primary antibodies: firstly with surface markers rat monoclonal against human CD 197 (CCR7) (M1 macrophage marker) (dilution 1:1000, Thermo-Fisher eBioscience™), goat polyclonal against human CD163 (M2c macrophage marker) (dilution 1:200, R&D Systems, USA), rabbit polyclonal against human CD206 (M2a marker) (dilution 1:1000, Abcam, USA) in DPBS with 1% normal donkey serum for 1 h in ice. Cells were then washed with PBS, permeabilized with 0.1% Triton X-100 (Sigma Aldrich, USA) for 15 min in ice. After three washes with DPBS, cells were incubated for 1 h on ice with a mouse monoclonal anti-human CD68 antibody, a pan-macrophage marker (1:1,000; Thermo Fisher Scientific, eBioscience), diluted in DPBS containing 1% normal donkey serum. Cells collected by centrifugation with ice cold methanol (−20 °C) for 10 min to fix them. The cells were then washed with PBS (pH 7.4) with 1% (v/v) BSA and then incubated with secondary antibody cocktail consisting of Alexa Fluor™ 405 conjugated donkey-anti mouse IgG (H+L) (dilution 1:1000), Alexa Fluor™ 488 conjugated donkey-anti rat IgG (H+L) (dilution 1:1000), Alexa Fluor™ 647 conjugated donkey-anti goat IgG (H+L) (dilution 1:1000), Alexa Fluor™ 568 conjugated donkey-anti rabbit IgG (H+L) (dilution 1:1000), for 30 min at RT. The stained cells were washed, resuspended in PBS, and analyzed using a Cytek Aurora spectral flow cytometer (Cytek Biosciences, USA). Prior to analyses, fluorescence minus one (FMO) controls and unstained populations were used to determine a proper gating strategy.

*Qualitative Analysis using Confocal Microscopy:* Cell-laden constructs were fixed using 4% paraformaldehyde overnight at 4 °C, washed thrice with DPBS. The gels were subjected to 15% (w/v) sucrose for 6 h and 30% sucrose overnight, after which they were retrieved from sucrose solution and embedded in Cryomatrix (Fisher-Scientific, USA). The embedded constructs were sectioned (30 μm) using a Leica CM1950 cryostat (Leica Biosystems, USA). The sectioned samples were stained for CCR7, CD206, CD68 using the same primary and secondary antibody combination. The immunostained samples were mounted using ProLong Gold Antifade Reagent with DAPI (Cell Signaling Technology, USA) and imaged using the LSM 880 Zeiss confocal microscope. The acquired images were processed and visualized as maximum-intensity projections.

### 4.10. Assessment of angiogenic and vascularization potential in EggMA hydrogel and microgel systems

*Tube Formation Assay:* The prepared E-HG at concentrations of 10 and 15% (w/v) were tested, while gelatin methacryloyl (GelMA, 10% w/v) and Matrigel matrix (growth factor reduced (GFR), phenol red-free, LDEV-free, Corning) served as control groups. 500 μL of each hydrogel was dispensed into 24-well plates and fully crosslinked. Human umbilical vein endothelial cells (Passage 4-6, HUVECs; Lonza), transduced to express tdTomato according to a previously established protocol (*72*), were expanded in DMEM medium (10% FBS and 1% P/S) and seeded on the surface of hydrogels at a density of 1 × 10^5^ cells/mL, and then cultured in EGM-2-based endothelial cell growth medium (EGM, Lonza) at 37 °C in a humidified incubator with 5% CO₂. Cellular morphology and microvascular network formation were monitored over the culture period using an EVOS fluorescence microscope (Thermo Fisher Scientific, Waltham, MA, USA), followed by quantitative analysis using ImageJ.

*Cell-Microgel Interactions:* To examine the effects of EggMA microgels (E-MG, 10% w/v) on cellular viability and behavior, non–labeled HUVECs were utilized and seeded onto individual microgels at a density of 1 × 10⁵ cells/mL. The prepared E-MG were distributed into 24-well suspension culture plates, and 500 μL of the HUVEC suspension was added to each well to allow cell attachment. After 3h incubation for cell attachment, additional 2 mL fresh EGM medium was added in each well. Cell viability, morphology, and functional marker expression on the microgel surfaces were systematically evaluated over the cultivation period, and cellular coverage on the microgel surfaces was further quantified using ImageJ.

*Spheroid-Microgel Interactions:* Human dermal fibroblasts (HDFs; Lonza) were expanded in DMEM medium (10% FBS and 1% P/S). The harvested HDFs (Passage 5-8) were mixed with HUVECs at a 1:2 ratio to obtain a total of 6,000 cells per spheroid for vascular spheroid formation. Briefly, the HUVEC/HDF cell suspension was dispensed into a U-bottom 96-well plate (Cat. No. 650970, Greiner Bio-One) at 6,000 cells per well and incubated in a humidified incubator at 37 °C with 5% CO₂ for 24 h to allow the formation of HUVEC/HDF spheroids (designated as vascular spheroids) with spheroid diameter around 280-350 µm. To investigate the interaction between spheroids and microgel matrix, vascular spheroids were aspirated using in-house established AAB system under a vacuum pressure of 10–18 mmHg, and bioprinted into E-MG and E-HG/MG matrices and crosslinked under 405 nm UV light for 2 min. Spheroid-contained constructs were cultivated in EGM medium for designated time points. Cellular proliferation, migration, and functional responses were assessed via immunofluorescence staining and quantitative real-time polymerase chain reaction (qPCR). Quantitative analyses were performed using ImageJ and Origin, and the reconstructed vascular networks were visualized and analyzed using Imaris.

*CAM Assay:* To evaluate the angiogenic potential of EggMA microgel-based constructs, the prepared E-HG@MG and E-MG scaffolds with low- and high-packing were assessed using an *in-ovo* CAM assay, as described in a previous protocol (*73*). E-HG scaffolds were used as a microgel-free control group. Briefly, fertilized chicken eggs were incubated under standard hatching conditions using a universal egg incubator (Peatend, China) with automatic rotation. After 7 days of incubation, a window was carefully opened in the eggshell above the air chamber under sterile conditions, and the scaffolds were implanted onto the CAM. The opened window was then sealed with sterilized transparent Parafilm, and the eggs were further incubated under standard conditions until embryonic Day 14. The implanted samples were collected and imaged using a camera (Pexil, Google), and the embryos were euthanized by prolonged exposure of carbon dioxide (∼25 min). The angiogenic potential and degree of vascular invasion were quantitatively analyzed using the AngioTool (*74*). Moreover, the explanted constructs together with surrounding CAM tissue, were fixed in 4% paraformaldehyde, cryosectioned, and subjected to histological and vascular analyses. Hematoxylin and eosin (H&E) staining was performed to assess tissue integration and vessel ingrowth within the scaffold interior. Host microvasculature was further visualized by fluorescence staining with Lycopersicon esculentum (tomato) lectin (10 µg/mL; Invitrogen, Thermo Fisher Scientific, Waltham, MA, USA) according to the manufacturer’s instructions, enabling assessment of the spatial distribution and depth of vascular infiltration within the constructs.

### 4.11. Characterization of Osteogenic Function

*Bioprinting of Bone Spheroids and Cultivation:* To investigate the osteogenic potential of EggMA microgels and their ability to promote bone tissue formation, primary human bone marrow mesenchymal stem cells (hMSC, Lonza, Walkersville, MD) were tested and cultured in α-modified minimum essential medium (α-MEM; Corning) with 10% FBS, 2 mM glutamine (Thermo Fisher Scientific) and 1% P/S. Harvested hMSC (passages 4-6) were utilized to generate hMSC spheroids using U-bottom 96-well plates, with 8,000 cells seeded per well for 24-48 h to form spheroids with approximately 300–350 μm in diameter. Subsequently, these spheroids were bioprinted within microgel-based matrices, including E-MG (low packing) and E-HG@MG, using AAB with 32G nozzle under a vacuum pressure of 15–20 mmHg. After photocrosslinking, hMSC spheroids-loaded E-MG and E-HG@MG constructs were incubated in a basal α-MEM medium containing 15% FBS and 1% P/S for 14 days under standard cultivation conditions (37 °C, 5% CO_2_). Afterward, human osteogenic differentiation medium (Cell Applications, San Diego, CA, denoted as OS^+^ medium) was applied for an additional 14 days until a total of 28 days of culture. For long-term cultivation, after 14 days of culture in the growth medium, the samples were transferred to the OS⁺ medium and maintained for 56, 84, and 112 days to evaluate osteogenic maturation and mineralized matrix formation. Cellular morphology, migration, proliferation, and functional activity within the microgel-based matrices were systematically evaluated over the cultivation period using (immuno)histochemical (IHC) staining, mRNA-level gene expression analysis, protein-level functional assays, tissue-level micro computed tomography (μCT) imaging, and physicochemical characterization of elemental composition as described below.

*μCT Imaging Evaluation:* Samples collected after 56, 84, and 112 days of cultivation were fixed in 4% PFA overnight at 4 °C and subsequently analyzed using a μCT scanner (SkyScan 1176; Bruker Micro-CT, Belgium). Scans were performed at 50 kV and 72 μA using a 0.25 mm aluminum filter, an isotropic pixel size of 9.13 μm, and a rotation step of 0.2° to evaluate mineralization and hardness of formed mini-bones. A hydroxyapatite (HA) calibration phantom (Micro-CT HA, QRM, Germany) was included in each scan to establish a standard curve for converting Hounsfield units to mg HA/ccm. Projection images were reconstructed using Nrecon Reconstruction software (version 2.2.0.6, Bruker Micro-CT) with ring artefact correction set to 20%, beam-hardening correction set to 12%, and a Hamming reconstruction filter (α = 0.54). Reconstructed datasets were visualized and analyzed using Avizo software (Thermo Fisher Scientific, Waltham, MA, USA). Mineralized regions were visualized using a grayscale threshold of 61–225 and rendered in purple; quantitative mineral analysis was performed using an HA-calibrated threshold of 200 mg HA/ccm.

*Evaluation of X-ray Diffraction (XRD):* The mineralized crystal structures of E-MG samples after 84 days of cultivation were analyzed using XRD to characterize bone-like nanocomposite formation. The samples were fixed, freeze-dried, and ground into fine powders prior to analysis. The powders were front-loaded onto a silicon zero-background sample holder, and diffraction data were collected over a 2θ range of 5–70° using a Malvern Panalytical Empyrean® diffractometer equipped with a copper (Kα₁ = 1.540598 Å, Kα₂ = 1.544426 Å) long-fine-focus X-ray tube operated at 45 kV and 40 mA. The incident beam path employed iCore® optics fitted with a BBHD® optic, 0.03-radian Soller slits, 14 mm primary and secondary beam masks, and a fixed 0.25° divergence slit. The diffracted beam path utilized dCore® optics with a 0.25° fixed anti-scatter slit and 0.04-radian Soller slits. A PIXcel® detector operated in scanning line (1D) mode with an active length of 3.347° 2θ was used, with data collected at a nominal step size of 0.026° 2θ. Phase identification was conducted using Jade® software (version 9.1, Materials Data Inc., Livermore, CA, USA) and the International Centre for Diffraction Data (ICDD) PDF-5® database. Acellular E-MG samples served as control group.

*Analysis of X-ray Photoelectron Spectroscopy (XPS):* E-MG samples after 84 days of cultivation were fixed and analyzed via XPS to determine the elemental composition and chemical states within the mineralized bone matrix. Acellular E-MG samples were analyzed in parallel as a control group. XPS measurements were conducted using a PHI VersaProbe III instrument (Physical Electronics, MN, USA) equipped with a monochromatic Al kα x-ray source (hν = 1,486.6 eV) and a concentric hemispherical analyzer. Charge neutralization was performed using both low energy electrons (<5 eV) and argon ions. The binding energy axis was calibrated using sputter cleaned Cu (Cu 2p3/2 = 932.62 eV, Cu 3p3/2 = 75.1 eV) and Au foils (Au 4f7/2 = 83.96 eV) reference foils, and spectra were charge-referenced to the C–Hₓ peak in the C 1s region at 284.8 eV (*75*). Measurements were collected at a take-off angle of 45° relative to the sample surface, corresponding to an approximate sampling depth of 3–6 nm (encompassing 95% of the detected signal). Elemental quantification was performed using instrument-specific relative sensitivity factor that account for X-ray cross sections and electron inelastic mean free paths. On homogeneous samples major elements (>5 atom%) tend to have standard deviations of <3% while minor elements can be significantly higher. The analyzed surface area was approximately 200 µm in diameter.

*Scanning Electron Microscopy (SEM) and Energy Dispersive X-ray Spectroscopy (EDS):* Cell-laden and acellular samples were collected at predetermined time points and lyophilized prior to analysis. The dried samples were mounted on double-sided carbon tape and sputter-coated with iridium to render the surface conductive. Surface morphology was examined using a scanning electron microscope (Apreo SEM, Thermo Fisher Scientific) operated under high vacuum at an accelerating voltage of 5–10 kV. Elemental composition was subsequently analyzed by an EDS detector integrated with the SEM at an accelerating voltage of 10-15 kV, and the data were processed using AZtec software (Oxford Instruments, UK).

### 4.12. Biochemical and Physicochemical Characterization

*LIVE/DEAD Assay:* Cell viability within EggMA bulk hydrogel and microgel-based matrices was evaluated using the LIVE/DEAD™ Viability/Cytotoxicity Kit (Thermo Fisher Scientific) following the manufacturer’s instructions. Briefly, a working solution was freshly prepared by adding 0.6 µL of calcein-AM (live-cell dye) and 1.2 µL of ethidium homodimer-1 (EthD-1, dead-cell dye) into 1 mL DPBS. At predetermined cultivation time points, the cell-laden constructs were rinsed with DPBS and incubated in the staining solution for 25 min on a shaker at RT, followed by three washes with DPBS. Z-stack images of live (green) and dead (red) cells in the constructs were captured using an EVOS fluorescence microscope and the LSM 880 Zeiss confocal microscope. The acquired images were processed and visualized as maximum-intensity projections or 3D reconstructions using Imaris. Quantitative analysis of cell viability was performed in ImageJ, where viability was calculated as the ratio of live cells to the total number of cells (i.e., the sum of living and dead cells).

*(Immuno)fluorescence Imaging:* At given time points of cultivation, cell/spheroid-laden constructs were fixed and permeabilized by using 4% PFA and 0.1% Triton X-100 (Merck KGaA, Darmstadt, Germany) in DPBS, respectively. Afterward, the nonspecific binding of antibodies/dyes was blocked by incubation in 2% bovine serum albumin (BSA) for 45 min in DPBS. To assess angiogenesis, the samples were incubated overnight at 4 °C with a mouse monoclonal anti-CD31 primary antibody (1:100 dilution, ab9498; Abcam). Following three washes with DPBS, the samples were incubated with a goat anti-mouse IgG (H+L) secondary antibody conjugated to Alexa Fluor 488 (1:200 dilution in PBS; A11017, Invitrogen) for 3h under dark conditions at RT. To evaluate osteogenic potential, the samples were similarly incubated overnight at 4 °C with rabbit polyclonal anti-COL1A1 (1:100 in 0.1% BSA; PA5-29569, Invitrogen) and mouse monoclonal anti-osteocalcin (OCN) (1:200 in 0.1% BSA; 33-5400, Invitrogen) primary antibodies. Following three washes with DPBS, the samples were incubated for 3 h of staining at RT in the dark with goat anti-mouse IgG (H+L) Alexa Fluor 488 (1:200 in DPBS; A11017, Invitrogen) and goat anti-rabbit IgG (H+L) Alexa Fluor 647 (1:200 in DPBS; A21245, Invitrogen) secondary antibodies. Moreover, to visualize cells and assess cellular morphology, nuclei and cytoskeletons were stained simultaneously using DAPI (0.1 µg/mL, Gibco) and Alexa Fluor™ 568 Phalloidin (1 µL/mL, Invitrogen). Post-staining, the z-stack images (thickness: 400–700 µm) were acquired using the LSM 880 Zeiss confocal microscope.

*Quantitative Real-Time Polymerase Chain Reaction (qPCR) Analysis:* At given time points of cultivation, cell/spheroid-laden constructs were collected, and total ribonucleic acid (RNA) content was extracted using previously published protocols (*76, 77*). Briefly, the total RNA content was extracted from constructs by Trizol™ reagent (Thermo Scientifi) and 1 µg of RNA was used for synthesis of cDNA using AccuPower® CycleScript RT PreMix (Bioneer, USA) in a thermal cycler machine (Bio-Rad T100, Bio-Rad, USA). The expression levels were estimated by Delta CT Method (−ΔCT), and represented as heat map, where relative expression level of gene of interest is calculated based on the equation, Relative expression, *R = 2 ^−(C^_T (GOI)_ ^− C^_T (ACTB)_*^)^. The basal expression levels of these genes (listed in Table S1) in free-standing spheroids with reference to endogenous house-keeping gene was used as baseline reference against the spheroid-laden constructs. The fluorescence signal was detected using Power SYBR PCR master mix (Applied Biosystems, Invitrogen, USA) in a real-time PCR machine (QuantStudio™ 5, Thermo Fisher Scientific).

*Western Blot Analysis:* At given time points of cultivation, spheroid-laden constructs were collected, and total protein from them were lysed using RIPA Lysis buffer (Thermo-Scientific) supplemented with 1× Protease/Phosphatase Inhibitor Cocktail (Cell Signaling Technologies, USA) and 1× LDS sample buffer (Invitrogen, USA). Samples were thawed, sonicated at 50% amplitude for 10 s in 2 s ON/OFF intervals, incubated at 95 °C for 5 min, and vortexed. Proteins were separated on 4–20% Mini-PROTEAN® TGX™ (Tris-Glycine eXtended) Gels (Bio-Rad, USA) and transferred using the Mini-PROTEAN® Tetra electrophoresis wet transfer system (Bio-Rad) onto an Immun-Blot PVDF Membrane (Bio-Rad). Blots were blocked in 5% nonfat dry milk (Bio-Rad) made in Tris-buffered saline containing 0.1% Tween 20 (TBST). Blots were incubated overnight at 4 °C with primary antibodies (listed in Supplementary Table 1) diluted in 1X TBST (from 10X TBST, Cell Signaling Technology, USA) containing 2% nonfat dry milk. Following primary incubation, blots were washed and incubated for 2 h at RT with horseradish peroxidase (HRP)-conjugated secondary antibodies (listed in Table S2) diluted in TBST with 2% nonfat dry milk. Blots were imaged using the Bio-Rad ChemiDoc MP Imaging System. Prestained ladder (10–245 kDa) (Abcam), Precision Plus Protein™ WesternC™ Blotting Standards, 10–250 kDa (Bio-Rad), Precision Protein™ StrepTactin-HRP Conjugate (dilution:1:10,000), Bio-Rad) were used as reference markers for determining the molecular weights of proteins of interest. When required, blots were stripped using Restore™ PLUS Western Blot Stripping Buffer (Thermo Scientific, USA) according to manufacturer’s instructions, thoroughly washed in TBST, and then reprobed with additional primary antibodies. After acquisition, densitometric analysis was performed using Image Lab (Bio-Rad, v5.1, build 8). Image brightness and contrast were uniformly adjusted in Image Lab. Band intensities were quantified after local background subtraction done through Image Lab and normalized to GAPDH loading controls. Only linear-range exposures were used for quantification, and adjustments to image brightness/contrast were applied uniformly across the entire blot without altering quantitative values. Normalized intensities were exported from Image Lab and were reported as fold-change relative to the indicated control.

### 4.13. Statistics and Reproducibility

All quantitative data were depicted as mean ± standard deviation (SD) from at least three independent biological replicates, unless otherwise specified. Statistical analyses were performed using one-way analysis of variance (ANOVA) followed by Tukey’s post hoc test, while a value of *p* ≤ 0.05 was considered statistically significant. All analyses were conducted using Origin 2017 software (OriginLab, USA). Details of specific statistical methods, sample size, and *p*-value results were included within the article and corresponding figure captions. Gene expression data were expressed as fold-changes (2⁻ΔΔCT ± upper/lower limit), with statistical comparisons performed on ΔCT values.

## Reference

1. S. V. Murphy, A. Atala, 3D bioprinting of tissues and organs. Nature Biotechnology 32, 773–785 (2014).

2. S. V. Murphy, P. De Coppi, A. Atala, Opportunities and challenges of translational 3D bioprinting. Nature Biomedical Engineering 4, 370–380 (2020).

3. Y. S. Zhang, A. Dolatshahi-Pirouz, G. Orive, Regenerative cell therapy with 3D bioprinting. Science 385, 604–606 (2024).

4. A. Schwab et al., Printability and Shape Fidelity of Bioinks in 3D Bioprinting. Chemical Reviews 120, 11028–11055 (2020).

5. G. Miklosic, S. J. Ferguson, M. D’Este, Engineering complex tissue-like microenvironments with biomaterials and biofabrication. Trends in Biotechnology 42, 1241–1257 (2024).

6. Y. Guan, L. Racioppi, S. Gerecht, Engineering biomaterials to tailor the microenvironment for macrophage–endothelium interactions. Nature Reviews Materials 8, 688–699 (2023).

7. N. E. Gregorio, C. M. Haas, N. P. King, C. A. DeForest, Engineering complexity into protein-based biomaterials for biomedical applications. Nature Reviews Materials 11, 328–342 (2026).

8. Y. Liu, A. E. Gilchrist, S. C. Heilshorn, Engineered Protein Hydrogels as Biomimetic Cellular Scaffolds. Advanced Materials 36, 2407794 (2024).

9. G. S. Hussey, J. L. Dziki, S. F. Badylak, Extracellular matrix-based materials for regenerative medicine. Nature Reviews Materials 3, 159–173 (2018).

10. X. Lin et al., Marine-Derived Hydrogels for Biomedical Applications. Advanced Functional Materials 33, 2211323 (2023).

11. Y. Li et al., Marine materials: New hope for biomedical applications. Cell Biomaterials.

12. T. Li et al., Plant-derived biomass-based hydrogels for biomedical applications. Trends in Biotechnology 43, 802–811 (2025).

13. M. M. Hasan, A. Ahmad, M. Z. Akter, Y. J. Choi, H. G. Yi, Bioinks for bioprinting using plant-derived biomaterials. Biofabrication 16, (2024).

14. C. Tuftee, E. Alsberg, I. T. Ozbolat, M. Rizwan, Emerging granular hydrogel bioinks to improve biological function in bioprinted constructs. Trends in Biotechnology 42, 339–352 (2024).

15. A. J. Graham et al., Stress-relaxing granular bioprinting materials enable complex and uniform organoid self-organization. Nature Materials, (2026).

16. Q. Feng, D. Li, Q. Li, X. Cao, H. Dong, Microgel assembly: Fabrication, characteristics and application in tissue engineering and regenerative medicine. Bioactive Materials 9, 105–119 (2022).

17. A. C. Daly, L. Riley, T. Segura, J. A. Burdick, Hydrogel microparticles for biomedical applications. Nature Reviews Materials 5, 20–43 (2020).

18. M. S. Huang, F. Christakopoulos, J. G. Roth, S. C. Heilshorn, Organoid bioprinting: from cells to functional tissues. Nature Reviews Bioengineering 3, 126–142 (2025).

19. K. J. Wolf, J. D. Weiss, S. G. M. Uzel, M. A. Skylar-Scott, J. A. Lewis, Biomanufacturing human tissues via organ building blocks. Cell Stem Cell 29, 667–677 (2022).

20. J. Schimelman et al., Biofabrication of Vascularized Tissues. Chemical Reviews, (2026).

21. K. D. Janson et al., Strategies for the vascular patterning of engineered tissues for organ repair. Nature Biomedical Engineering 9, 1007–1025 (2025).

22. Z. A. Sexton et al., Rapid model-guided design of organ-scale synthetic vasculature for biomanufacturing. Science 388, 1198–1204 (2025).

23. T. Xie et al., Microgels for 3D Biofabrication. Aggregate 6, e70166 (2025).

24. D. R. Griffin, W. M. Weaver, P. O. Scumpia, D. Di Carlo, T. Segura, Accelerated wound healing by injectable microporous gel scaffolds assembled from annealed building blocks. Nature Materials 14, 737–744 (2015).

25. K. L. Wilson, S. C. L. Pérez, M. M. Naffaa, S. H. Kelly, T. Segura, Stoichiometric Post-Modification of Hydrogel Microparticles Dictates Neural Stem Cell Fate in Microporous Annealed Particle Scaffolds. Advanced Materials 34, 2201921 (2022).

26. V. G. Muir et al., Influence of Microgel and Interstitial Matrix Compositions on Granular Hydrogel Composite Properties. Advanced Science 10, 2206117 (2023).

27. K. Yue et al., Structural analysis of photocrosslinkable methacryloyl-modified protein derivatives. Biomaterials 139, 163–171 (2017).

28. H.-B. Yang et al., Multiscale integral synchronous assembly of cuttlebone-inspired structural materials by predesigned hydrogels. Nature Communications 16, 62 (2025).

29. T. M. Squires, S. R. Quake, Microfluidics: Fluid physics at the nanoliter scale. Reviews of Modern Physics 77, 977–1026 (2005).

30. P. Garstecki, M. J. Fuerstman, H. A. Stone, G. M. Whitesides, Formation of droplets and bubbles in a microfluidic T-junction—scaling and mechanism of break-up. Lab on a Chip 6, 437–446 (2006).

31. T. Moragues et al., Droplet-based microfluidics. Nature Reviews Methods Primers 3, 32 (2023).

32. F. Nie, D. Yan, Bio-sourced flexible supramolecular glasses for dynamic and full-color phosphorescence. Nature Communications 15, 9491 (2024).

33. D. B. Emiroglu et al., Building block properties govern granular hydrogel mechanics through contact deformations. Science Advances 8, eadd8570.

34. Y. Xing et al., Origin of the critical state in sheared granular materials. Nature Physics 20, 646–652 (2024).

35. A. C. Daly, Granular Hydrogels in Biofabrication: Recent Advances and Future Perspectives. Advanced Healthcare Materials 13, 2301388 (2024).

36. R. Merkel, P. Nassoy, A. Leung, K. Ritchie, E. Evans, Energy landscapes of receptor–ligand bonds explored with dynamic force spectroscopy. Nature 397, 50–53 (1999).

37. M. H. Kim et al., High-throughput bioprinting of spheroids for scalable tissue fabrication. Nature Communications 15, 10083 (2024).

38. B. Ayan et al., Aspiration-assisted bioprinting for precise positioning of biologics. Science Advances 6, eaaw5111 (2020).

39. K. Sadtler et al., Design, clinical translation and immunological response of biomaterials in regenerative medicine. Nature Reviews Materials 1, 16040 (2016).

40. C. Backlund, S. Jalili-Firoozinezhad, B. Kim, D. J. Irvine, Biomaterials-Mediated Engineering of the Immune System. Annual Review of Immunology 41, 153–179 (2023).

41. A. Salehi Moghaddam et al., Engineering the Immune Response to Biomaterials. Advanced Science 12, 2414724 (2025).

42. M. Peng et al., Macrophages: Subtypes, Distribution, Polarization, Immunomodulatory Functions, and Therapeutics. MedComm 6, e70304 (2025).

43. X. Shi et al., Biomaterial-mediated macrophage polarization remodeling and sequential regulation: a potential strategy in bone infections treatment. Bone Research 13, 96 (2025).

44. Y. Zhu et al., Regulation of macrophage polarization through surface topography design to facilitate implant-to-bone osteointegration. Science Advances 7, eabf6654.

45. J. Li, X. Jiang, H. Li, M. Gelinsky, Z. Gu, Tailoring Materials for Modulation of Macrophage Fate. Advanced Materials 33, 2004172 (2021).

46. Z. Chen et al., A Stone-Cottage-Inspired Printing Strategy to Build Microsphere Patterned Scaffolds for Accelerated Bone Regeneration. Advanced Functional Materials 35, 2417836 (2025).

47. A. E. Widener, A. Roberts, E. A. Phelps, Granular Hydrogels for Harnessing the Immune Response. Advanced Healthcare Materials 13, 2303005 (2024).

48. A. J. Wolf, D. M. Underhill, Peptidoglycan recognition by the innate immune system. Nature Reviews Immunology 18, 243–254 (2018).

49. S. Pinzón Martín, P. H. Seeberger, D. Varón Silva, Mucins and Pathogenic Mucin-Like Molecules Are Immunomodulators During Infection and Targets for Diagnostics and Vaccines. Frontiers in Chemistry Volume 7–2019, (2019).

50. A. C. Dudley, A. W. Griffioen, Pathological angiogenesis: mechanisms and therapeutic strategies. Angiogenesis 26, 313–347 (2023).

51. J. Rouwkema, A. Khademhosseini, Vascularization and Angiogenesis in Tissue Engineering: Beyond Creating Static Networks. Trends in Biotechnology 34, 733–745 (2016).

52. M. T. Collins et al., Skeletal and extraskeletal disorders of biomineralization. Nature Reviews Endocrinology 18, 473–489 (2022).

53. M. Zhao, X. Zhang, C. Xiao, X. Chen, Biomimetic Mineralized Hydrogels for Bone Regeneration. Advanced Materials 38, e20380 (2026).

54. G. L. Koons, M. Diba, A. G. Mikos, Materials design for bone-tissue engineering. Nature Reviews Materials 5, 584–603 (2020).

55. N. Shang, J. Wu, Egg White Ovotransferrin Shows Osteogenic Activity in Osteoblast Cells. Journal of Agricultural and Food Chemistry 66, 2775–2782 (2018).

56. Z. Hamidouche et al., Priming integrin α5 promotes human mesenchymal stromal cell osteoblast differentiation and osteogenesis. Proceedings of the National Academy of Sciences 106, 18587–18591 (2009).

57. W. L. Murphy, T. C. McDevitt, A. J. Engler, Materials as stem cell regulators. Nature Materials 13, 547–557 (2014).

58. M. Brunner et al., Osteoblast mineralization requires β1 integrin/ICAP-1–dependent fibronectin deposition. Journal of Cell Biology 194, 307–322 (2011).

59. C. Liu et al., Exploring the potential of hydroxyapatite-based materials in biomedicine: A comprehensive review. Materials Science and Engineering: R: Reports 161, 100870 (2024).

60. A. L. Boskey, E. Villarreal-Ramirez, Intrinsically disordered proteins and biomineralization. Matrix Biology 52-54, 43–59 (2016).

61. S. Akkineni et al., Amyloid-like amelogenin nanoribbons template mineralization via a low-energy interface of ion binding sites. Proceedings of the National Academy of Sciences 119, e2106965119 (2022).

62. A. George, A. Veis, Phosphorylated Proteins and Control over Apatite Nucleation, Crystal Growth, and Inhibition. Chemical Reviews 108, 4670–4693 (2008).

63. D. Qin et al., Collagen-based biocomposites inspired by bone hierarchical structures for advanced bone regeneration: ongoing research and perspectives. Biomaterials Science 10, 318–353 (2022).

64. A. Saraswathibhatla, D. Indana, O. Chaudhuri, Cell–extracellular matrix mechanotransduction in 3D. Nature Reviews Molecular Cell Biology 24, 495–516 (2023).

65. R. Levato et al., Light-based vat-polymerization bioprinting. Nature Reviews Methods Primers 3, 47 (2023).

66. C.-F. He, T.-H. Qiao, G.-H. Wang, Y. Sun, Y. He, High-resolution projection-based 3D bioprinting. Nature Reviews Bioengineering 3, 143–158 (2025).

67. V. Pal et al., Interparticle Crosslinked Ion-Responsive Microgels for 3D and 4D (Bio)Printing Applications. Small 21, e02262 (2025).

68. X. Hu, Q. Hu, S. Liu, H. Zhang, Synergy of engineered gelatin methacrylate-based porous microspheres and multicellular assembly to promote osteogenesis and angiogenesis in bone tissue reconstruction. International Journal of Biological Macromolecules 283, 137228 (2024).

69. M. H. Kim, D. Banerjee, N. Celik, I. T. Ozbolat, Aspiration-assisted freeform bioprinting of mesenchymal stem cell spheroids within alginate microgels. Biofabrication 14, (2022).

70. J. C. Moses et al., In Situ Nitric Oxide Generating Nano-Bioactive Glass-Based Coatings and Its Therapeutic Ion Release toward Attenuating Implant-Associated Fibrosis and Infection. Small 21, 2411984 (2025).

71. B.-H. Cha et al., Integrin-Mediated Interactions Control Macrophage Polarization in 3D Hydrogels. Advanced Healthcare Materials 6, 1700289 (2017).

72. M. Dey et al., Chemotherapeutics and CAR-T Cell-Based Immunotherapeutics Screening on a 3D Bioprinted Vascularized Breast Tumor Model. Advanced Functional Materials 32, 2203966 (2022).

73. S. Liu et al., Evaluation of different crosslinking methods in altering the properties of extrusion-printed chitosan-based multi-material hydrogel composites. Bio-Design and Manufacturing 6, 150–173 (2023).

74. E. Zudaire, L. Gambardella, C. Kurcz, S. Vermeren, A Computational Tool for Quantitative Analysis of Vascular Networks. PLOS ONE 6, e27385 (2011).

75. M. P. Seah, Summary of ISO/TC 201 Standard: VII ISO 15472 : 2001—surface chemical analysis—x-ray photoelectron spectrometers—calibration of energy scales. Surface and Interface Analysis 31, 721–723 (2001).

76. J. C. Moses, S. Dey, A. Bandyopadhyay, M. Agarwala, B. B. Mandal, Silk-Based Bioengineered Diaphyseal Cortical Bone Unit Enclosing an Implantable Bone Marrow toward Atrophic Nonunion Grafting. Advanced Healthcare Materials 11, 2102031 (2022).

77. T. D. Schmittgen, K. J. Livak, Analyzing real-time PCR data by the comparative CT method. Nature Protocols 3, 1101–1108 (2008).

