## Supplementary Information for "Bioinstructive Orthogonally-crosslinked Ovoprotein Microgels for Modular Bioprinting"

**This SI file includes:**

**Supplementary Tables 1-2**

**Supplementary Figs. 1-38**

**Table S1** Primer sequences of genes investigated in this study

| GENE | SEQUENCE | ACCESSION NUMBER |
|---|---|---|
| Human β-actin (*ACTB*) | **F** 5′- GGCATCCTCACCCTGAAGTA-3′<br>**R** 5′- GGGGTGTTGAAGGTCTCAAA-3′ | NM_001101.5 |
| Human angiopoietin 2 (*ANGPT2*) | **F** 5′- GCAAAATAAGCAGCATCAGCCAAC-3′<br>**R** 5′- GCATCAAACCACCAGCCTCCT-3′ | NM_001386335.1 |
| Human CD34 (*CD34*) | **F** 5′- AGCTGTGCGGAGTTTAAGAAGGAC-3′<br>**R** 5′- TCACCTCAGACTGGGCAAGGAG-3′ | NM_001025109.2 |
| Human podocalyxin like 2 (PODXL2) | **F** 5′- GGAGGAGATTGGCATCCAGAACT-3′<br>**R** 5′- TGACCACCAGCACCACGAAGA-3′ | NM_015720.4 |
| Human Vascular endothelial growth factor receptor 2 (*VEGFR2)* | **F** 5′- CCAGATGACAACCAGACGGACAG-3′<br>**R** 5′- GGCACCATTCCACCAAAAGATG-3′ | NM_002253.4 |
| Human C-X-C motif chemokine receptor 4 (*CXCR4*) | **F** 5′- CGGTTACCATGGAGGGGATCAGTATATA-3′<br>**R** 5′- GCATTTTCTTCACGGAAACAGGGTTC-3′ | NM_003467.3 |
| Human C-X-C motif chemokine ligand 12 (*CXCL12*) | **F** 5′- GCTACAGATGCCCATGCCGATTC-3′<br>**R** 5′- CAGCCGGGCTACAATCTGAAG-3′ | NM_199168.4 |
| Human protocadherin 12 (*PCDH12*) | **F** 5′- AGGCAGCAGGAGTGCAATCC-3′<br>**R** 5′- CTGGGGTCCAGCAGACTTGACA-3′ | NM_016580.4 |
| Human alpha-2-macroglobulin (*A2M*) | **F** 5′- GAAGCCAACAGTGAAAATGCTTGA-3′<br>**R** 5′- CAAGCTCAGTGTCTGATTTGACACCT-3′ | NM_000014.6 |
| Human plexin D1 (*PLXND1*) | **F** 5′- GACCCCGACACCCTACACATCT-3′<br>**R** 5′- CCTGCGCGATGACTGAAAGG-3′ | NM_015103.3 |
| Human von Willebrand factor (*VWF*) | **F** 5′- CCGTGACTTGCCAGCCAGATG-3′<br>**R** 5′- AGCCACAGGTCTCTTCCACTTTAAC-3′ | XM_054373133.1 |
| Human platelet and endothelial cell adhesion molecule 1 (*PECAM or CD31*) | **F** 5′- TGCAGTGGTTATCATCGGAGTG-3′<br>**R** 5′- CGTTGTTGGAGTTCAGAAGTG-3′ | XM_054316431.1 |
| Human nitric oxide synthase 3 (*eNOS3*) | **F** 5′- GGAACCTGTGTGACCCTCA-3′<br>**R** 5′- CGAGGTGGTCCGGGTATCC-3′ | NM_000603.5 |
| Human thrombomodulin (*THBD*) | **F** 5′- TGCCGATGTCATTTCCTTGCTACTG-3′<br>**R** 5′- GTTGTTGTCTCCCGTAACCCACTG-3′ | NM_000361.3 |
| Human endoglin (ENG or CD105) | **F** 5′- CCTCAACATGGACAGCCTCTCTTTC-3′<br>**R** 5′- AGCAGGAACTCGGAGACGGATG-3′ | NM_001114753.3 |
| Human matrix metallopeptidase 2 (*MMP2*) | **F** 5′- ATCGAGACCATGCGGAAGC-3′<br>**R** 5′- GCCCGAGCAAAAGCATCAT-3′ | NM_001302508.1 |

| | | |
|---|---|---|
| Human matrix metallopeptidase 9 (*MMP9*) | **F** 5′- TTCTACGGCCACTACTGTGC-3′<br>**R** 5′- GAATCGCCAGTACTTCCCATC-3′ | NM_004994.3 |
| Human collagen 1A (*COL1A*) | **F** 5′- GCACCACGGCAGCAGGAG-3′<br>**R** 5′- TTTGACCAGGTTCACCAGGCTC-3′ | NM_000089.4 |
| Human Runt-related transcription factor 2 (*RUNX2*) | **F** 5′-GATGGGACTGTGGTTACTGTCA-3′<br>**R** 5′-CTCAGATCGTTGAACCTTGC-3′ | NM_001278478.1 |
| Human Osterix or SP7 (*OSX*) | **F** 5′- CCTCTGCGGGACTCAACAAC-3′<br>**R** 5′- AGCCCATTAGTGCTTGTAAAGG-3′ | NM_001300837.2 |
| Human osteocalcin or BGLAP (*OCN)* | **F** 5′- TCACACTCCTCGCCCTATTG-3′<br>**R** 5′- TCGCTGCCCTCCTGCTTG-3′ | NM_199173.2 |

**Table S2.** Antibodies used for Western blot

| Antibodies | Dilution | Manufacturer |
|---|---|---|
| GAPDH Mouse monoclonal | 1:2000 | Abcam |
| XP® Anti-YAP Rabbit Monoclonal | 1:1000 | Cell Signaling Technology |
| Sp7/Osterix Rabbit monoclonal | 1:1000 | Abcam |
| Runt-related transcription factor 2 (RUNX2) mouse monoclonal | 1:1000 | Abcam |
| Alkaline phosphatase (ALP) Rabbit monoclonal | 1:1000 | Abcam |
| Goat anti-Rabbit IgG (H+L) – HRP conjugated | 1:10,000 | Fisher Scientific |
| Goat anti-Mouse IgG (H+L) – HRP conjugated | 1:2000 | Fisher Scientific |

**A** Proteins that could enhance osteogenesis

| EggMA protein detected (Accession number) | Human ortholog | Human UniProt number |
|---|---|---|
| Ovotransferrin (A0A411G5W6, Q4ADJ7, A0A8V0ZZF1) | Transferrin | P02787 |
| Ovalbumin-related protein X (A0A8V0ZFT5) | Transferrin | P02787 |
| Annexin (A0A8V0YMS3) | No mapping rule matched | |
| Fibrillar collagen NC1 domain-containing protein (A0A8V0ZLQ9) | COL3A1 | P02461 |
| Ankyrin repeat and fibronectin type-III domain-containing protein 1-like (A0A8V0Z946) | FN1 | P02751 |
| Fibronectin type III and SPRY domain-containing protein 1 (A0A8V1AG81) | FN1 | P02751 |

**B** Proteins that could enhance angiogenesis

| EggMA protein detected (Accession number) | Human ortholog | Human UniProt number |
|---|---|---|
| Clusterin (A0A8V0Y0J2) | Clusterin | P10909 |
| Clusterin associated protein 1 (A0A8V0Z465) | Clusterin | P10909 |
| Ovotransferrin (A0A411G5W6, Q4ADJ7, A0A8V0ZZF1) | Transferrin | P02787 |
| Fibrillar collagen NC1 domain-containing protein (A0A8V0ZLQ9) | COL3A1 | P02461 |
| Ankyrin repeat and fibronectin type-III domain-containing protein 1-like (A0A8V0Z946) | FN1 | P02751 |

**C** Immunomodulatory proteins

| EggMA protein detected (Accession number) | Human ortholog | Human UniProt number |
|---|---|---|
| Ovalbumin (P01012) | No mapping rule matched | |
| Ovomucin, alpha subunit (A0A8V1A1T7) | MUC5B/ MUC6 | P10909 |
| Lysozyme C (P00698) | Lysozyme | P61626 |

**Figure S1. Human ortholog mapping of EggMA proteins associated with regenerative functions.** Among the 788 proteins identified in EggMA, 181 were mapped to human orthologs and screened using Reactome analysis for putative functional associations with **A,** osteogenesis, **B,** angiogenesis and **C,** immunomodulation.

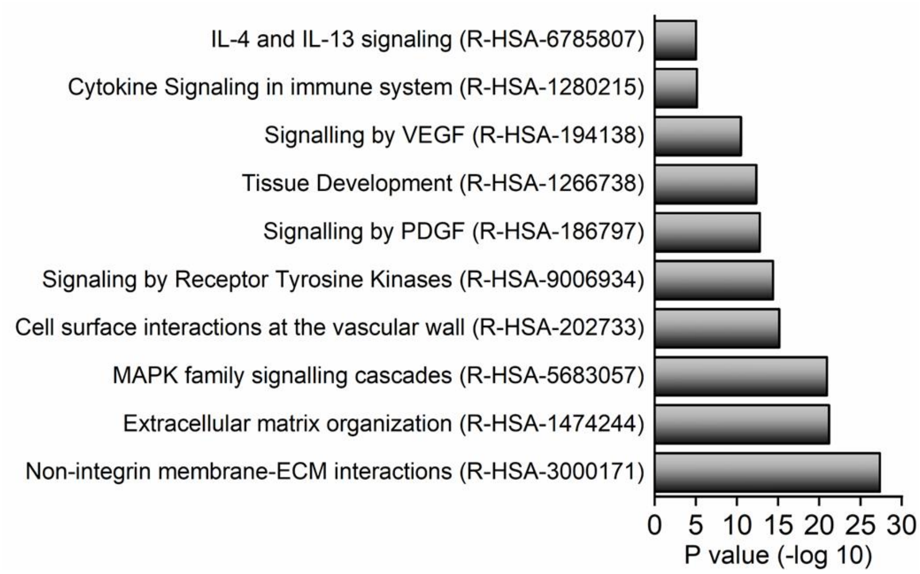


**Figure S2. Reactome pathway enrichment of human orthologs mapped from EggMA proteins.** Reactome pathway enrichment of human orthologs mapped from EggMA proteins, highlighting biological processes associated with ECM organization, membrane–ECM interactions, MAPK and receptor tyrosine kinase signaling, vascular regulation, tissue development and immune-related cytokine signaling.

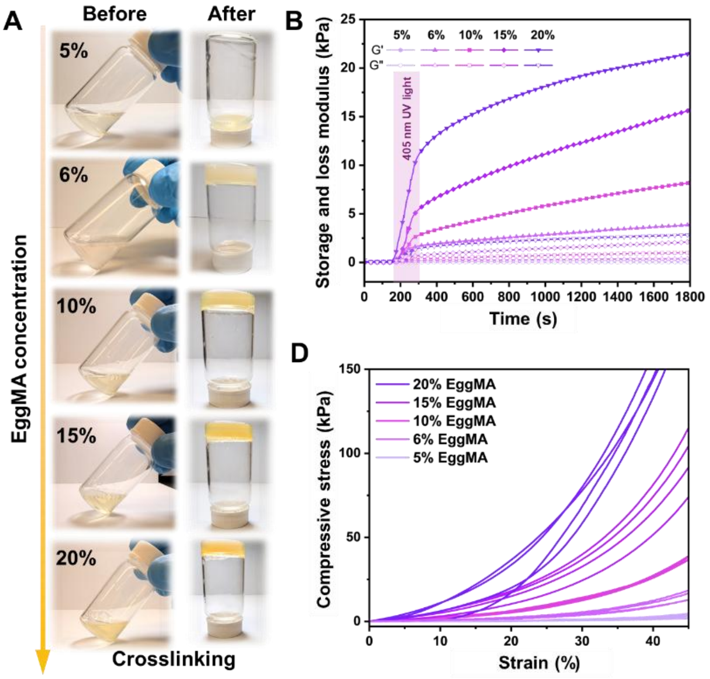


**Figure S3. Concentration-dependent photocrosslinking and mechanical properties of EggMA hydrogels.**

**A,** Vial-inversion test of EggMA precursor solutions before and after visible-light photocrosslinking at different concentrations. **B,** Time-sweep rheological analysis showing *G′* and *G″* modulus during 405 nm light-induced gelation for 2 min, following a 3-min baseline stabilization period. **C,** Frequency-sweep rheological analysis of photocrosslinked EggMA hydrogels. **D,** Representative compressive stress–strain curves of EggMA hydrogels with different concentrations. **E,** Quantification of compressive modulus as a function of EggMA concentration. Data were presented as mean ± SD ($n$ = 3); **$P$ < 0.01 and ***$P$ < 0.001.

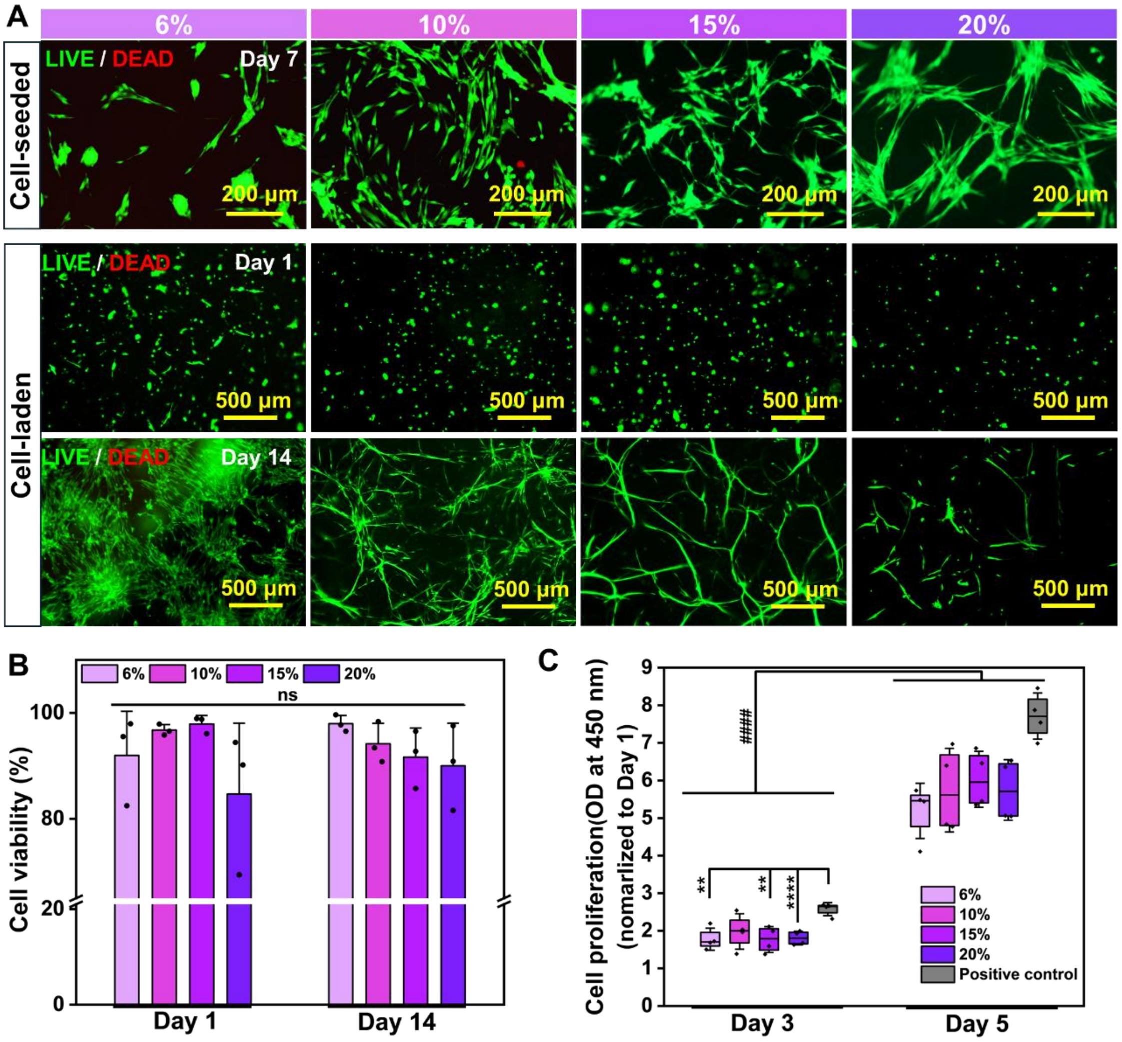


**Figure S4. Concentration-dependent NHLF response to the bulk EggMA hydrogel.** **A,** LIVE/DEAD fluorescence images of NHLFs seeded on or encapsulated within bulk EggMA of different concentrations, showing cell viability, spreading and morphology over culture. **B,** Quantification of NHLF viability in EggMA hydrogels at Days 1 and 14. Data were presented as mean ± SD ($n$ = 3); n.s., not significant. **C,** NHLF proliferation in EggMA assessed by CCK-8 assay, quantified by absorbance at 450 nm and normalized to Day 1. Data were presented as box plots with mean ± SD ($n$ = 4). $^{**}P < 0.01$ and $^{****}P < 0.0001$ denote comparisons among groups at the same time point; $^{\#\#\#\#}P < 0.0001$ denotes comparisons between time points within the same group.

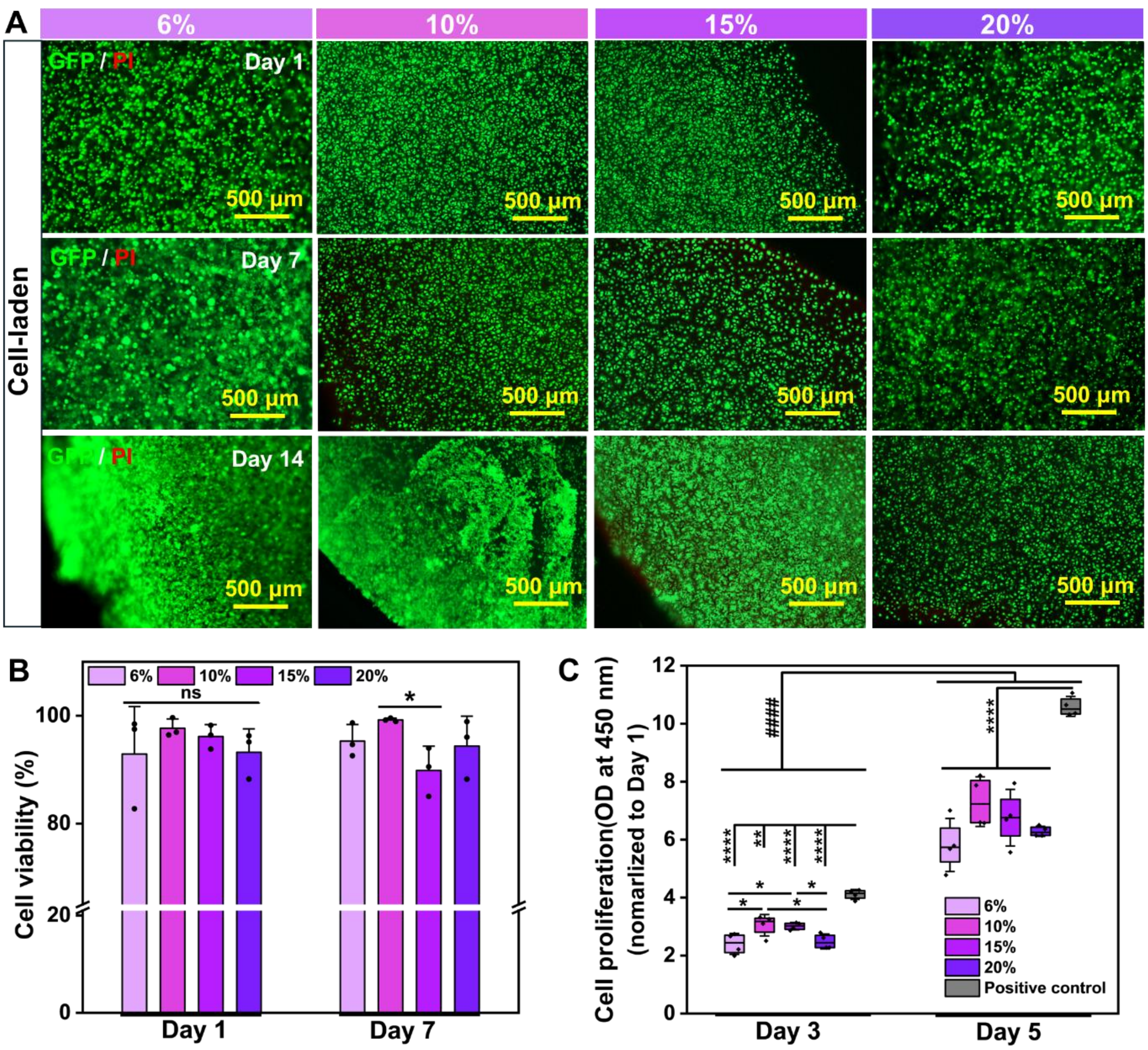


**Figure S5. Concentration-dependent MDA-MB-231 cell response to the bulk EggMA hydrogel.**

**A,** Fluorescence images of $GFP^+$ MDA-MB-231 cells encapsulated within EggMA of different concentrations, with propidium iodide (PI) staining dead cells, showing cell viability and proliferation over culture. **B,** Quantification of cell viability at Days 1 and 7. Data were presented as mean ± SD ($n$ = 3); n.s., not significant; $^*P < 0.05$. **C,** Proliferation of MDA-MB-231 in EggMA assessed by CCK-8 assay, quantified by measuring the optical density (OD) at 450 nm and normalized to Day 1. Data were presented as box plots with mean ± SD ($n$ = 4). $^*P < 0.05$, $^{**}P < 0.01$, and $^{****}P < 0.0001$ denote comparisons among groups at the same time point; $^{\#\#\#\#}P < 0.0001$ denotes comparisons between time points within the same group.

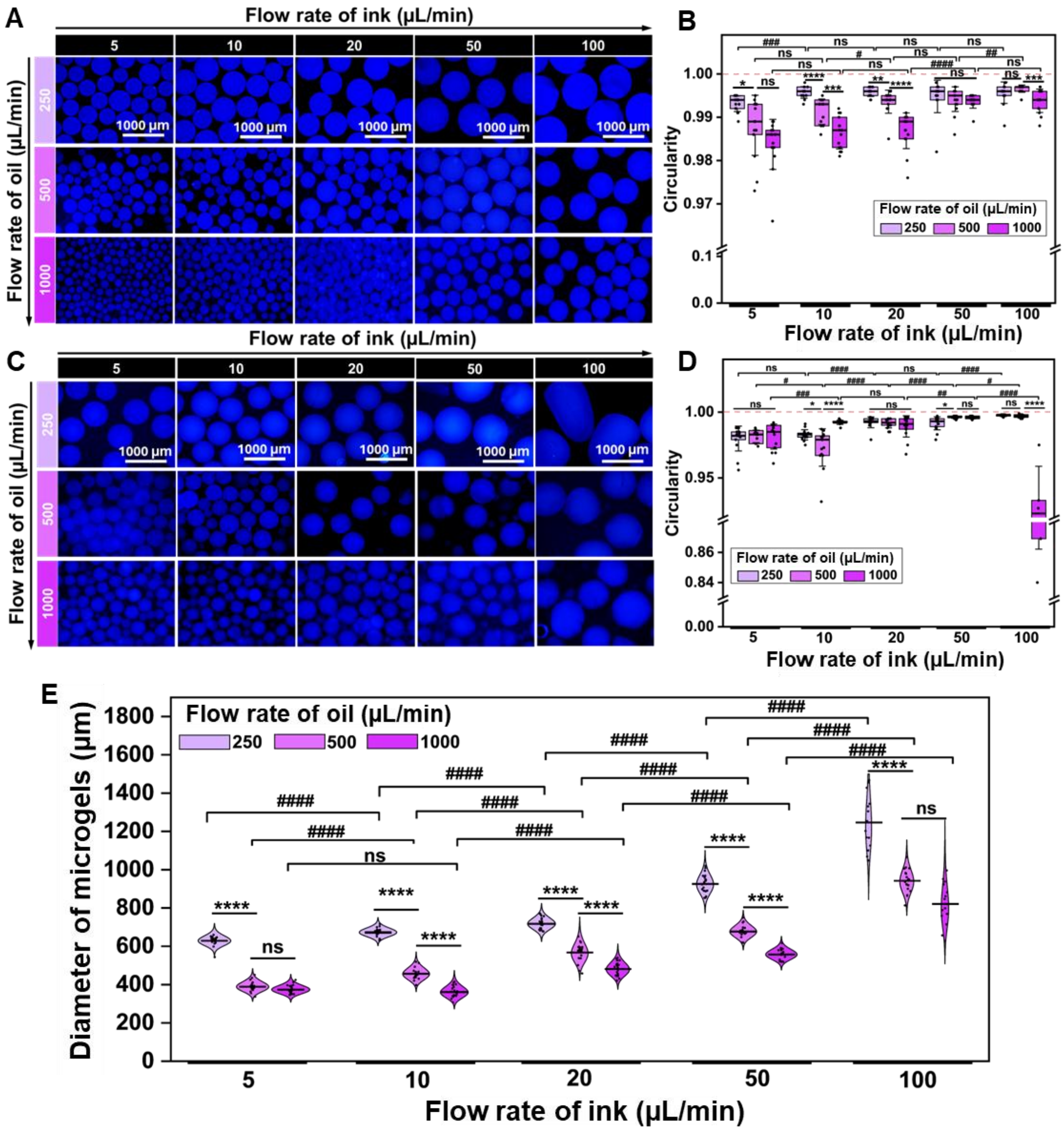


**Figure S6. Flow-rate-dependent control of EggMA microgel morphology and size.**
**A,** Autofluorescence images of 10% EggMA microgels generated under varying dispersed-phase (EggMA ink) and continuous-phase (oil) flow rates, showing flow-dependent changes in microgel morphology and size. **B,** Circularity analysis of 10% EggMA microgels under the corresponding flow conditions, related to Fig. 1F1. **C,** Autofluorescence images of 15% EggMA microgels generated under varying EggMA ink and oil flow rates, showing flow-dependent changes in microgel morphology and size. **D-E,** Corresponding circularity analysis and diameter distributions of 15% EggMA microgels. Data were presented as mean ± SD (*n* ≥ 100 microgels per condition). $^{*}P < 0.05$, $^{**}P < 0.01$, $^{***}P < 0.001$, and $^{****}P < 0.0001$ denote comparisons among oil flow rates at fixed ink flow rate; $^{\#}P < 0.05$, $^{\#\#}P < 0.01$, $^{\#\#\#}P < 0.001$, and $^{\#\#\#\#}P < 0.0001$ denote comparisons among ink flow rates at fixed oil flow rate; n.s., not significant.

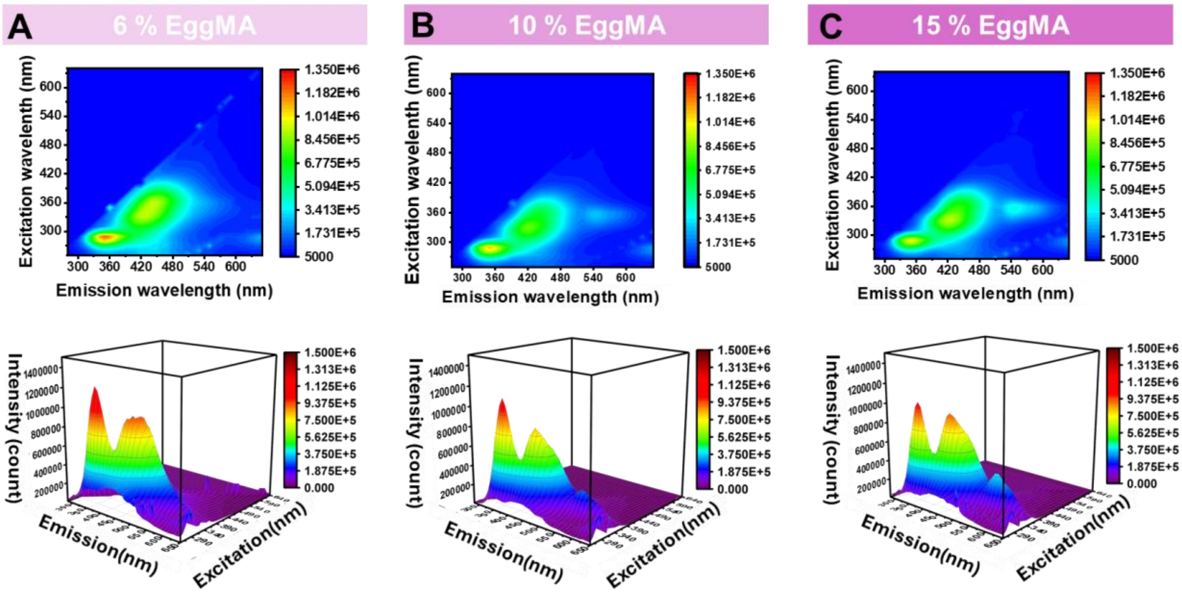


**Figure S7. Intrinsic autofluorescence of the photocrosslinked bulk EggMA hydrogel.** Photoluminescence excitation–emission mapping and corresponding 3D fluorescence intensity profiles of vinyl-crosslinked EggMA hydrogels at **A,** 6; **B,** 10; and **C,** 15% (w/v), revealing well-defined intrinsic autofluorescence across all tested concentrations. Distinct emission features were observed in the high-energy aromatic residue-associated region and the broader blue-emission region corresponding to the DAPI filter window used for E-MG visualization.

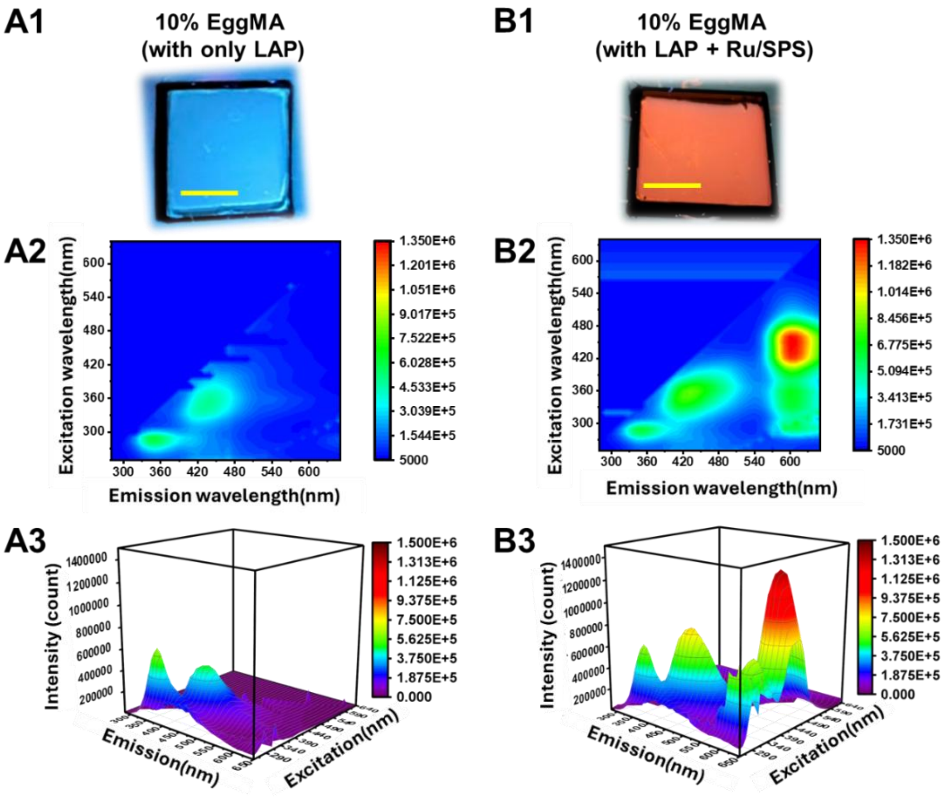


**Figure S8. Effect of orthogonal Ru/SPS crosslinking on EggMA autofluorescence. A1–A3,** A representative fluorescence image, photoluminescence excitation–emission map and 3D fluorescence intensity profile of 10% EggMA crosslinked with LAP only; scale bar, 500 μm. **B1–B3,** A representative fluorescence image, photoluminescence excitation–emission map and 3D fluorescence intensity profile of 10% EggMA orthogonally crosslinked with LAP + Ru/SPS. Scale bar, 500 μm. Ru/SPS-mediated dityrosine coupling enhanced autofluorescence intensity and shifted emission toward the ~500–600 nm range, accompanied by a visible transition from blue to orange fluorescence.

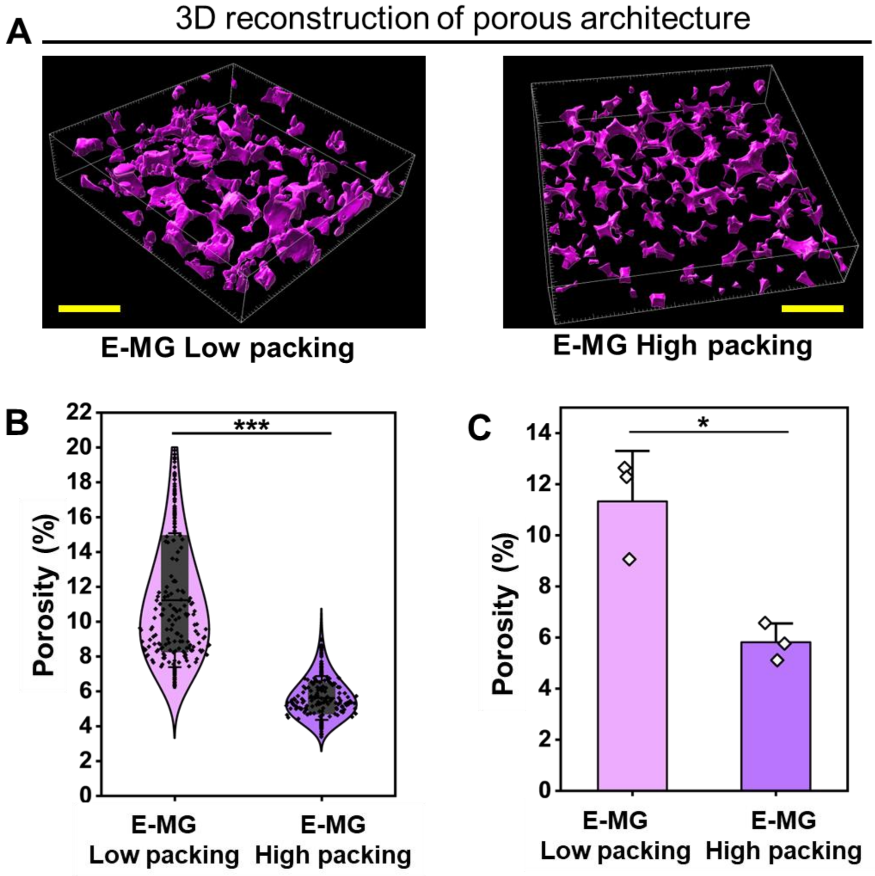


**Figure S9. Packing-density-dependent microporous architecture of E-MG.**
**A,** Representative 3D reconstructions of the interstitial pore architecture in low- and high-packing E-MG generated by centrifugation at 300 and 1300 mg, respectively. Scale bar. 300 μm. **B-C,** Quantification of porosity from **B,** 2D image projections and **C,** 3D volumetric reconstructions, showing reduced porosity with increased packing density. Data were presented as mean ± SD (*n*=3); **P* < 0.05 and ****P* < 0.001.

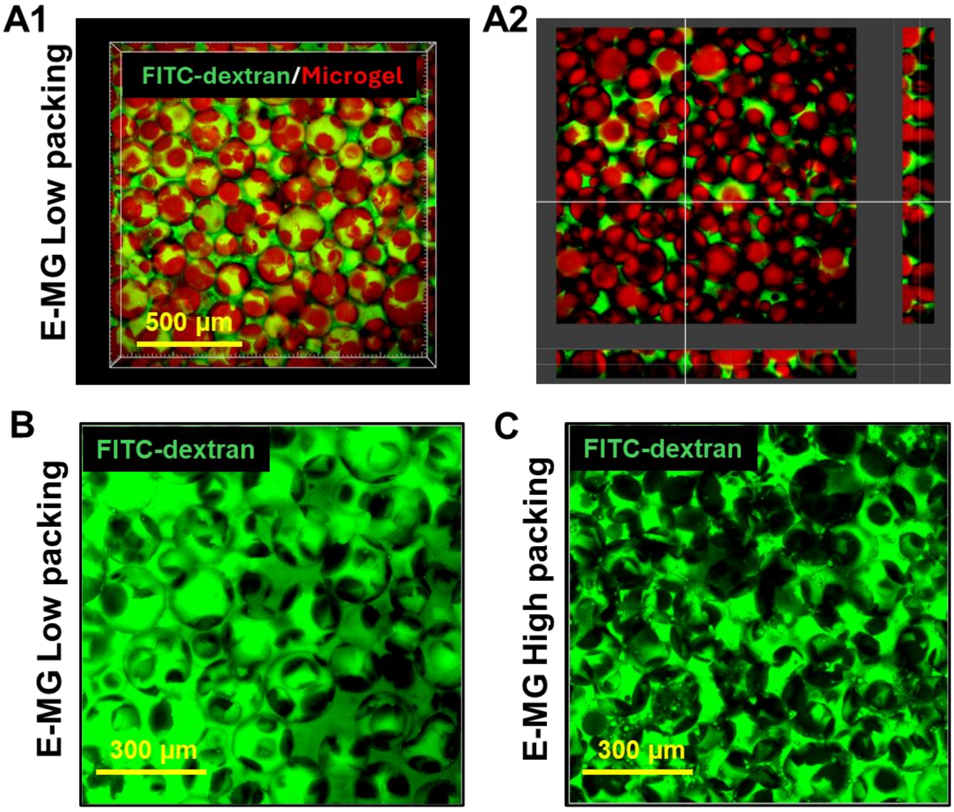


**Figure S10. Visualization of interstitial permeability in E-MG with different packing densities.** Representative fluorescent images of FITC–dextran diffusion within E-MG revealed the interconnected interstitial space between microgels. **A1,** 3D confocal reconstruction and **A2,** orthogonal views of a low-packing E-MG assembly, with FITC–dextran shown in green and autofluorescent microgels in red, illustrating the continuous permeable pore network. **B-C,** representative fluorescence images of FITC–dextran distribution in low- and high-packing E-MG, highlighting greater interstitial accessibility in the low-packing configuration.

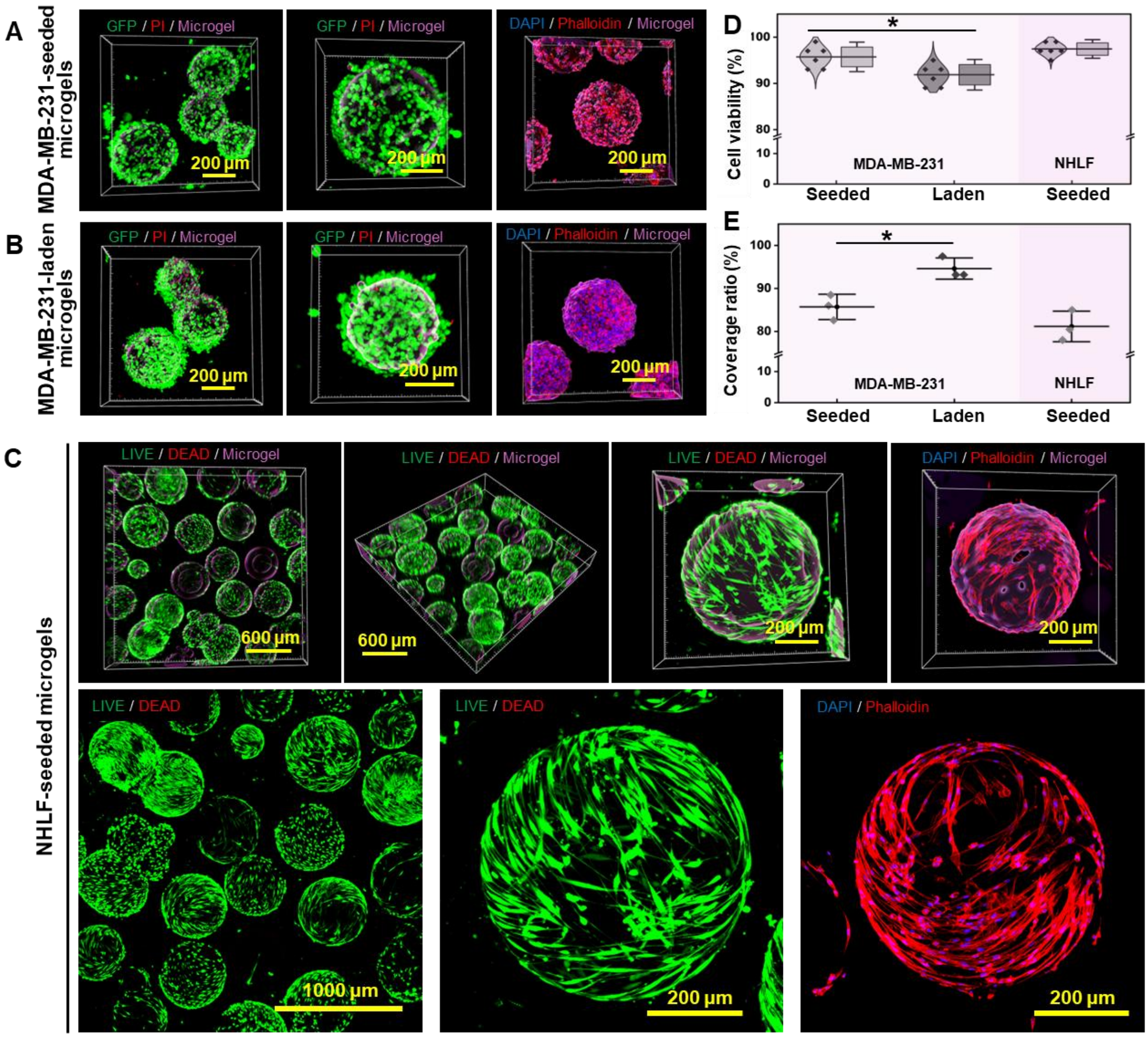


**Figure S11. Cytocompatibility and cellular colonization within ovoprotein microgels.** **A,** Representative confocal images of MDA-MB-231-seeded E-MG constructs after 7 days of culture, showing viable cells (GFP, green; PI, red) associated with the microgel matrix (3D rendering, purple) and exhibiting progressive surface attachment and spreading. Cytoskeletal organization was further visualized by phalloidin staining (red), with nuclei counterstained with DAPI (blue). **B,** Representative confocal images of MDA-MB-231-laden E-MG constructs at Day 7, showing high cell viability, pronounced proliferation and construct-wide colonization. **C,** Representative LIVE/DEAD and confocal images of NHLF-seeded E-MG at Day 7, demonstrating high cytocompatibility, extensive cell survival, and progressive fibroblast spreading and network formation along the microgel surfaces. **D,** Quantification of cell viability for MDA-MB-231-seeded, MDA-MB-231-laden, and NHLF-seeded microgel constructs. **E,** Quantification of microgel surface coverage ratio, indicating substantial cellular spreading and colonization within microgel matrices. Data were presented as mean ± SD ($n$=3); *$P < 0.05$.

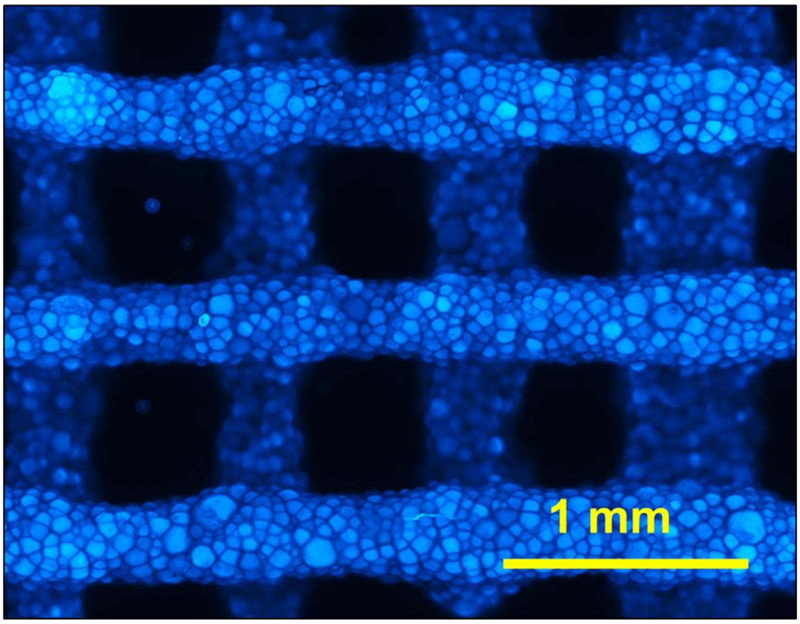


**Figure S12.** High-resolution fluorescent imaging of printed E-MG grid structure via EBB, showing densely packed microgels within continuous filaments and preserved macroporous architecture. This image corresponds to Fig. 2F.

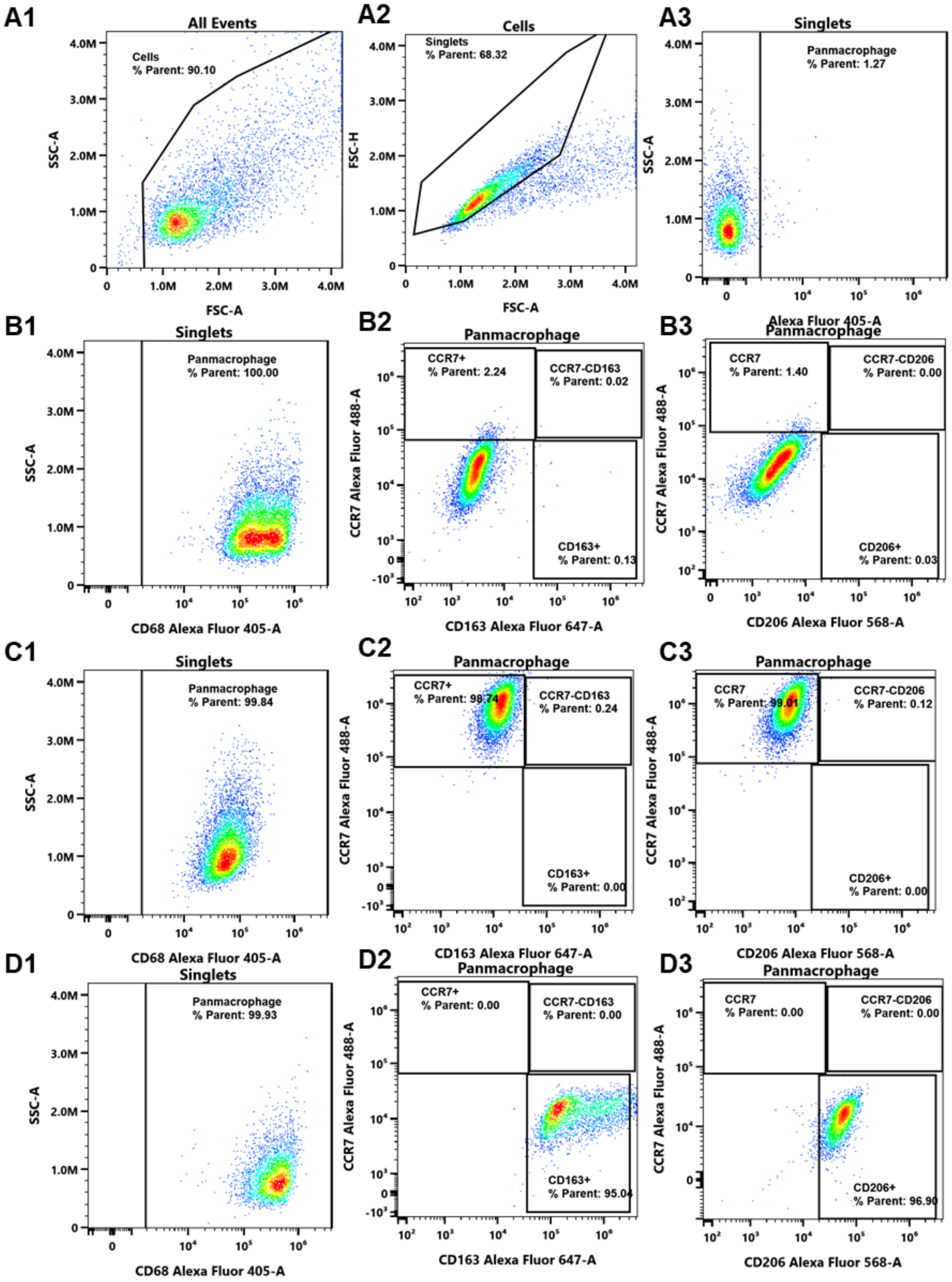


**Figure S13. Flow-cytometric gating strategy and validation of THP-1-derived macrophage polarization.**

**A1-A3,** Gating strategy followed to gate (**A1**) parent cells devoid of cell debris, (**A2**) gated parent population to doublet and proceed to (**A3**) gate singlets, which were CD68$^{+}$ (pan-macrophage) using non-stained cells. **B1-B3,** Naïve M0 macrophages without polarization treatment, showing **B1**, $CD68^+$ macrophage selection and baseline CCR7 (**B2**), CD163 and CD206 gates (**B3**). **C1-C3,** Naïve macrophages treated with inflammatory IFN-γ to yield M1 polarized macrophages exhibiting (**C1**) CD68+; (**C2-C3**) ≥ 98% $CCR7^+$, $CD163^-$, and $CD206^-$ cells. **D1-D3,** Naïve macrophages treated with anti-inflammatory IL-4 to induce M2 polarization, exhibiting (**D1**) $CD68^+$; (**D2-D3**) ≥ 95% $CCR7^-$, $CD163^+$, $CD206^+$ cells.

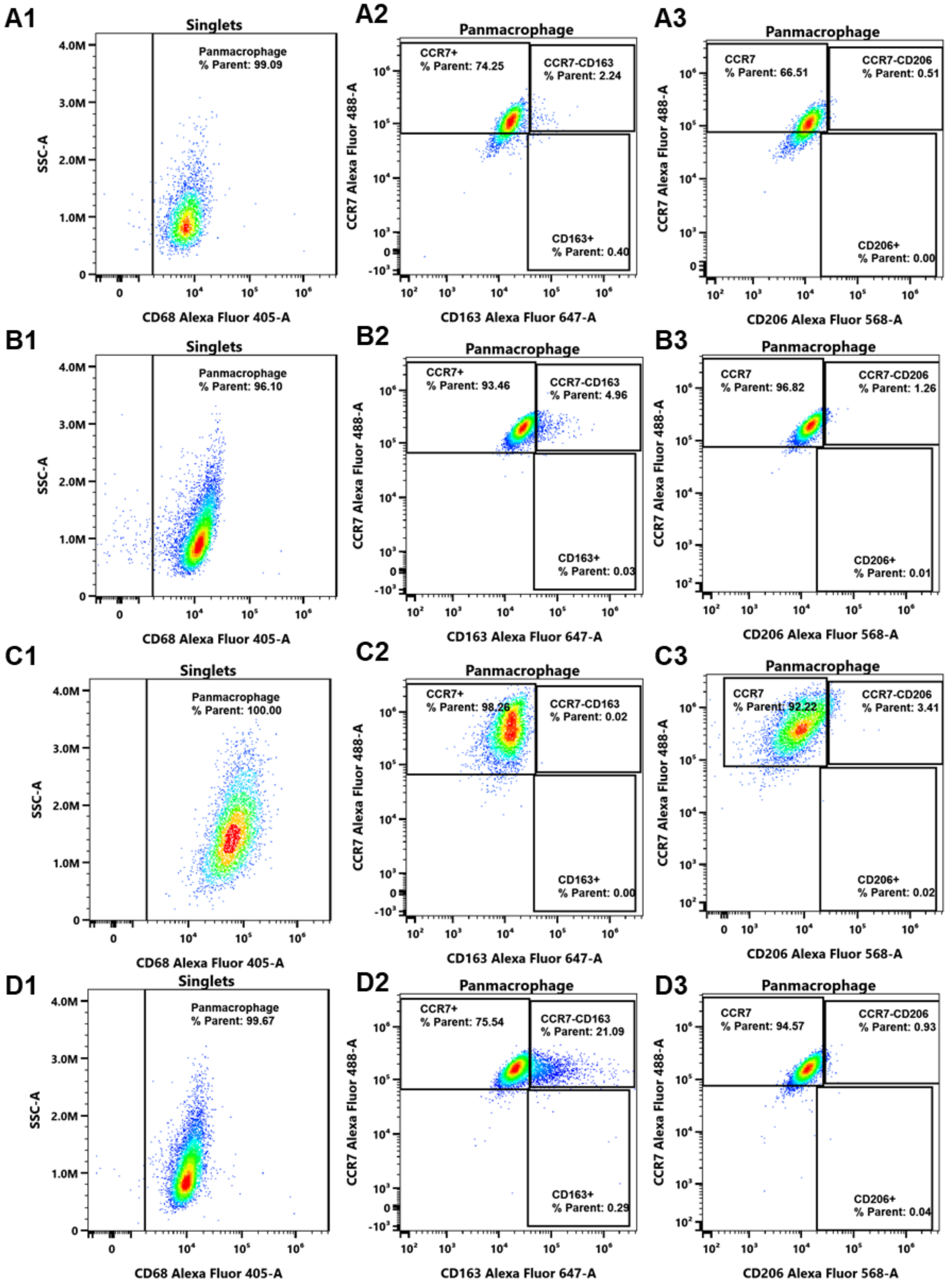


**Figure S14. Flow-cytometric analysis of macrophage responses in GelMA-based control matrices.**

THP1-monocytes derived macrophages encapsulated in GelMA-HG at **A1-A3,** Day 3 and **C1-C3,** Day 7 , and in GelMA-HG@MG at **B1-B3,** Day 3 and **D1-D3,** Day 7. Gating strategy followed for (**A1,B1,C1,D1**) CD68$^{+}$ population and proceeded to gate for (**A2,B2,C2,D2**)

CCR7/CD163, and (**A3,B3,C3,D3**) CCR7/CD206 cell population analysis.

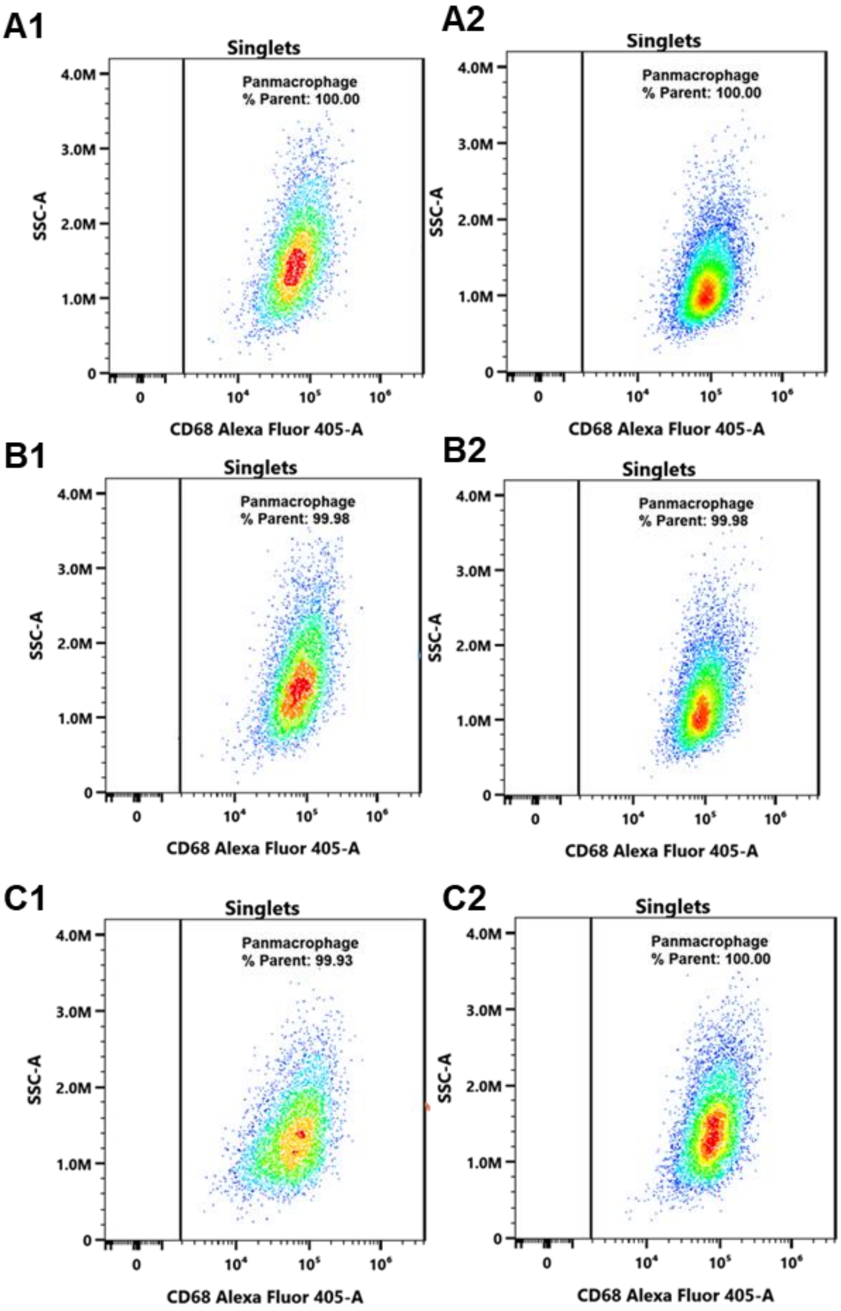


**Figure S15. CD68$^+$ macrophage gating for ovoprotein-based matrices.**

Flow cytometric analysis for immune response assessed by THP1-monocytes derived macrophages encapsulated in **A1-A2,** E-HG, **B1-B2,** E-MG, **C1-C2,** E-HG@MG, at (**A1, B1, C1**) Day 3 and (**A2, B2, C2**) Day 7 representing the CD68+ population, which was proceeded to gate for CCR7/CD163, and CCR7/CD206 cell populations represented in Figs. 3 A1, B1, and C1.

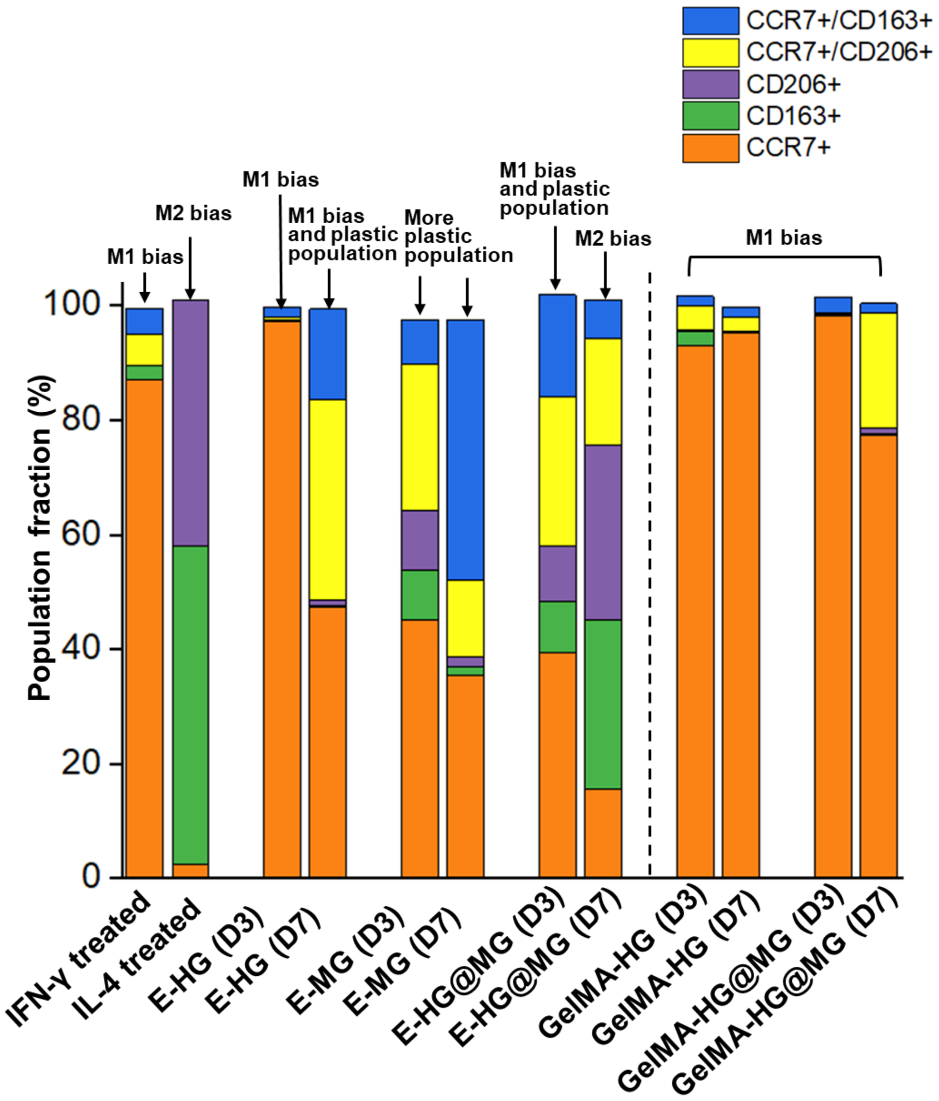


**Figure S16. Comparative macrophage polarization profiles across ovoprotein- and GelMA-based matrices.**

Flow cytometric analysis representing the population distributions from the macrophages retrieved at Day 3 (D3) and Day 7 (D7) from the encapsulated samples and compared against controls IFN-γ and IL-4 treated naïve M0 macrophages. While GelMA-HG and GelMA-HG@MG samples represent predominantly M1-biased macrophage populations, the E-HG, E-MG and E-HG@MG groups showcased (D7) reduced M1-alone populations and more plastic populations of $CCR7^+/CD163^+$ and $CCR7^+/CD206^+$ states over time. E-HG@MG group exhibited predominant M2-bias, with enriched $CD163^+$ and $CD206^+$populations, indicating that ovoprotein microgel-containing matrices promote macrophage plasticity and reparative polarization while attenuating sustained pro-inflammatory activation.

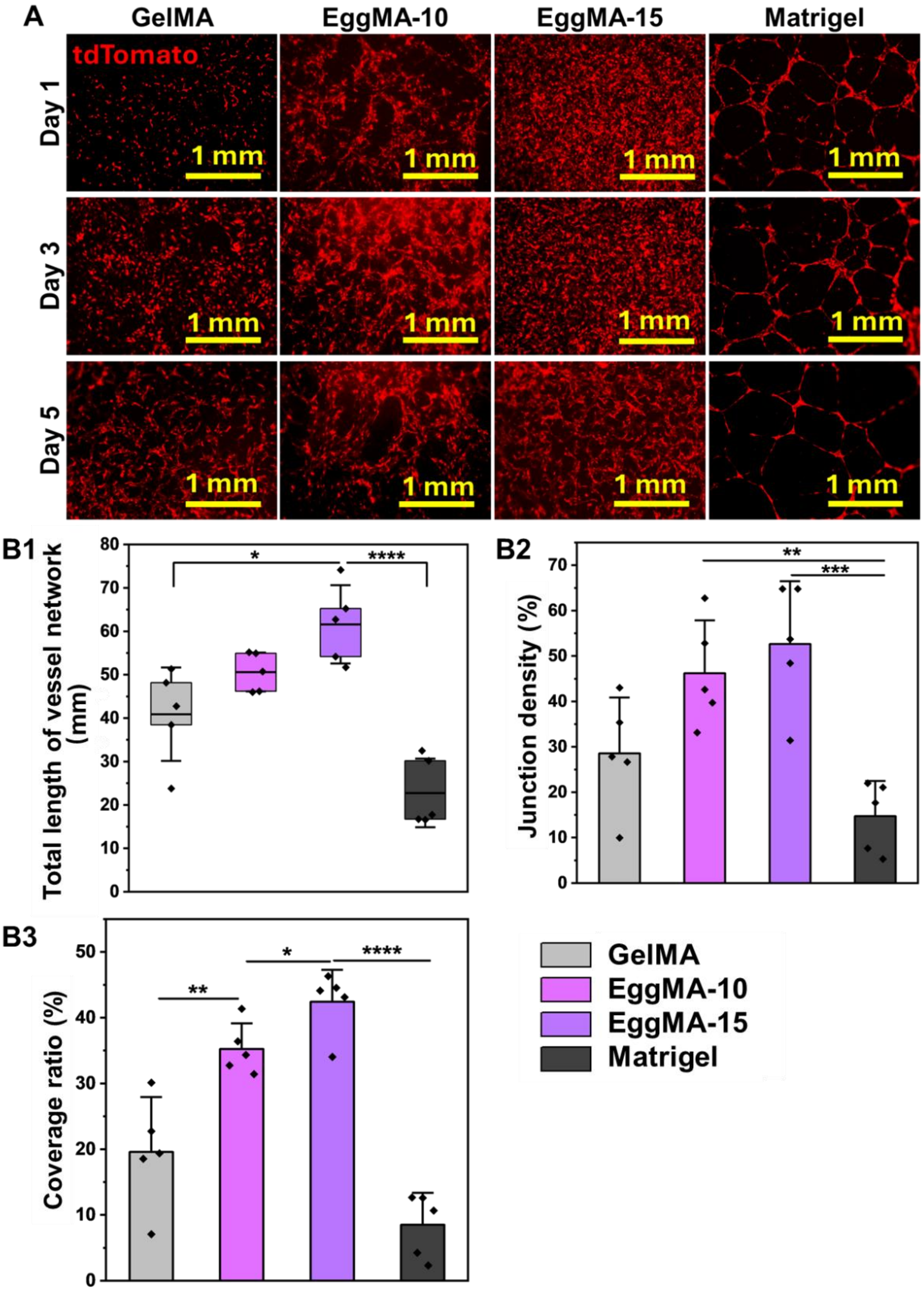


**Figure S17. Endothelial tube formation in bulk EggMA hydrogels.**

**A,** Representative fluorescence images of tdTomato-labelled HUVECs cultured on GelMA, EggMA-10, EggMA-15, and Matrigel matrices at Days 1, 3 and 5, showing matrix-dependent capillary-like network formation. **B1–B3,** Quantification of total vessel network length, junction density, and cellular coverage ratio after 5 days of cultivation, demonstrating enhanced endothelial network formation in bulk EggMA hydrogels compared with GelMA matrix. Data were presented as mean ± SD ($n$ = 5); *$P$ < 0.05, **$P$ < 0.01, ***$P$ < 0.001, and ****$P$ < 0.0001.

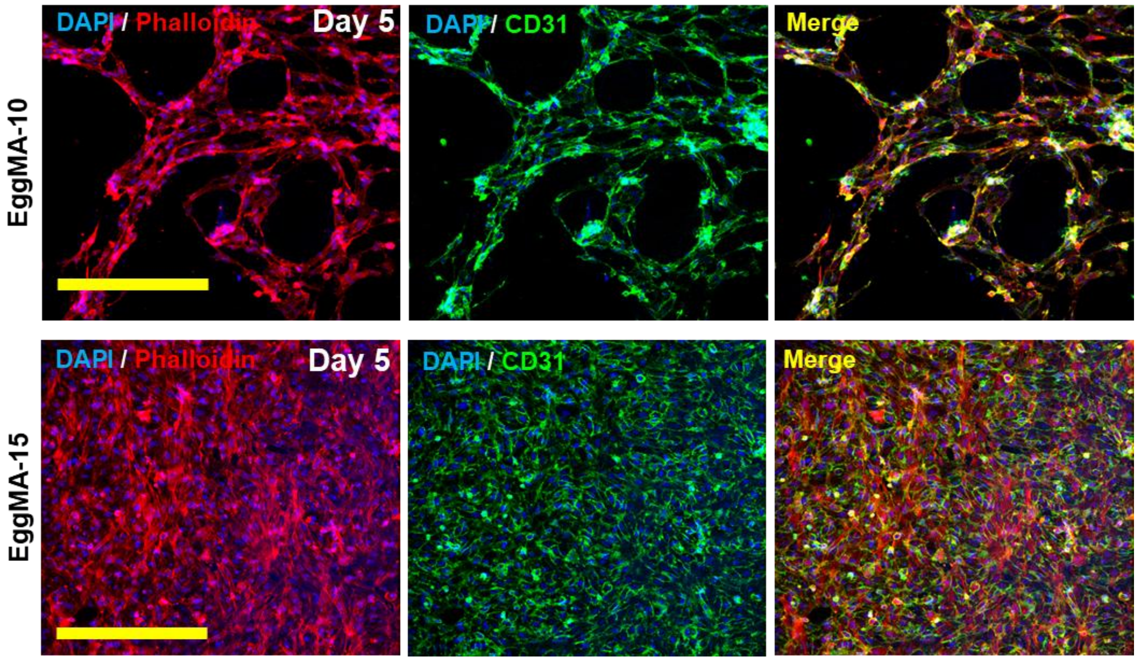


**Figure S18. CD31⁺ endothelial organization on EggMA-based matrices.** Representative immunofluorescence images of HUVECs cultured on EggMA-10 and EggMA-15 hydrogels at Day 5, stained for CD31 (green), phalloidin-labelled F-actin (red) and DAPI-counterstained nuclei (blue), showing endothelial organization and CD31$^{+}$ network formation. Scale bars, 400 µm.

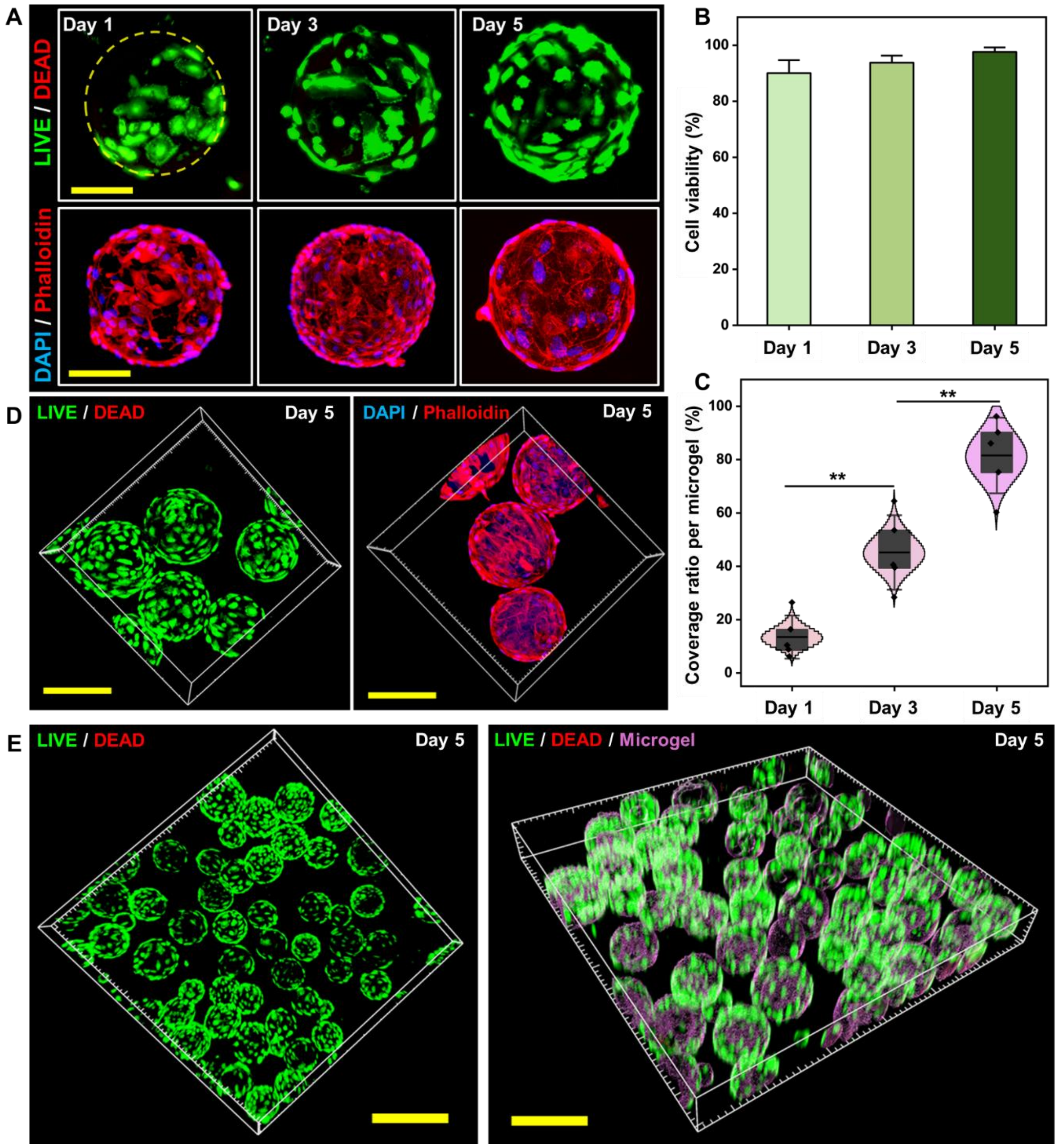


**Figure S19. Endothelial viability, spreading and construct-scale colonization within ovoprotein microgels.**

**A,** Representative fluorescent images of HUVECs cultured on E-MG microgels at Days 1, 3 and 5, stained for LIVE/DEAD (green/red) and DAPI/phalloidin (blue/red), showing sustained viability together with progressive cell spreading and cytoskeletal organization over time, Scale bar, 100 μm. **B,** Quantification of HUVEC viability during culture, confirming high cell vaibility maintained on microgels. Data were presented as mean ± SD (*n*=3). **C,** Quantification of endothelial coverage ratio per microgel over time, showing a significant increase in microgel surface colonization during culture. Data were presented as mean ± SD (*n*=5); ***P* < 0.01. **D-E,** Representative large-area 3D confocal views of HUVEC-populated E-MG at Day 5, showing uniform cytocompatibility, extensive cell survival, attachment and spreading, as well as construct-wide endothelial colonization and interconnected multicellular organization. Scale bar, **D,** 200 μm and **E,** 600 μm.

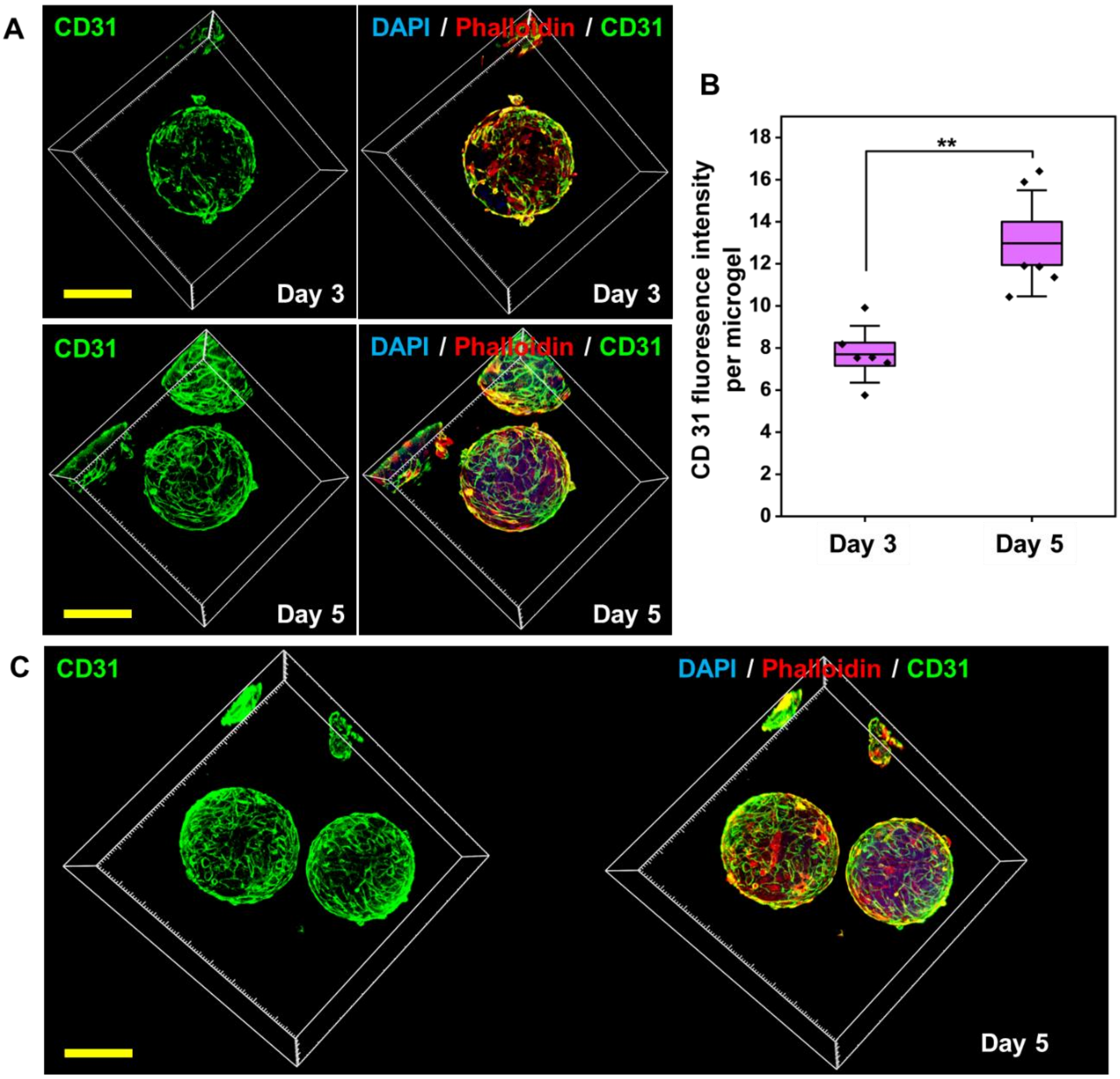


**Figure S20. CD31⁺ endothelial maturation within HUVEC-seeded ovoprotein microgels. A,** Representative 3D flourencent images of HUVEC-seeded ovoprotein microgels at Days 3 and 5, stained for CD31 (green), phalloidin-labelled F-actin (red) and DAPI-counterstained nuclei (blue), showing progressive endothelial organization on microgels. Scale bar, 200 μm. **B,** Quantification of CD31 fluorescence intensity per microgel demonstrates increased endothelial marker expression over time. Data were presented as mean ± SD; $n$ = 6 microgels per group; $^{**}P < 0.01$. **C,** Representative 3D fluorescence images showing uniform CD31⁺ endothelial coverage across multiple microgels at Day 5. Scale bar, 200 μm.

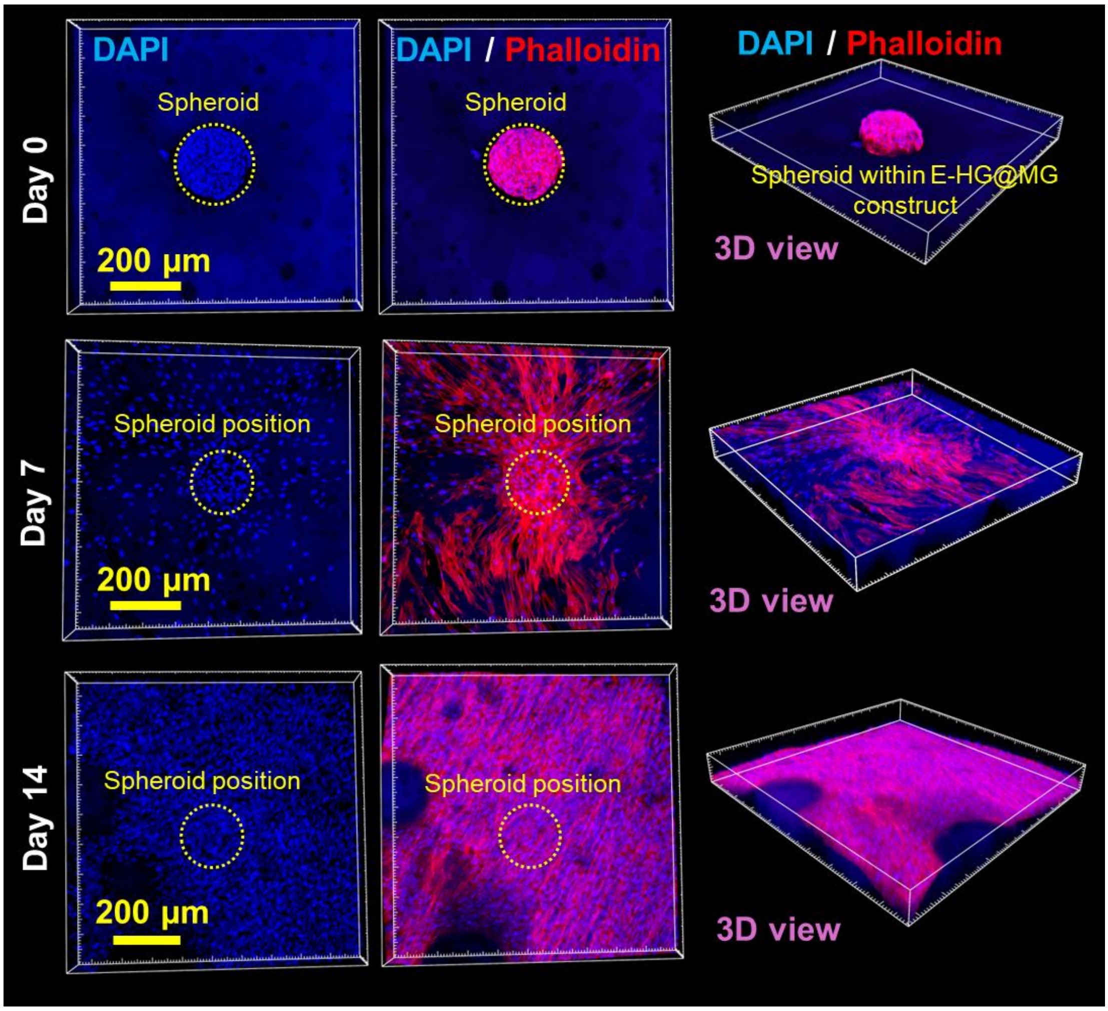


**Figure S21. Time-dependent outgrowth and dispersion of vascular spheroids within E-HG@MG.** Representative fluorescent images and 3D view of vascular spheroids cultured within E-HG@MG constructs at Days 0, 7 and 14. Nuclei were stained with DAPI (blue) and the actin cytoskeleton with phalloidin (red). The images show the initially confined spheroid position post bioprinting (Day 0), followed by progressive cellular outgrowth, proliferative expansion and radial invasion into the surrounding microgel matrix during culture, accompanied by gradual loss of the original spheroid boundary and increasing cellular dispersion.

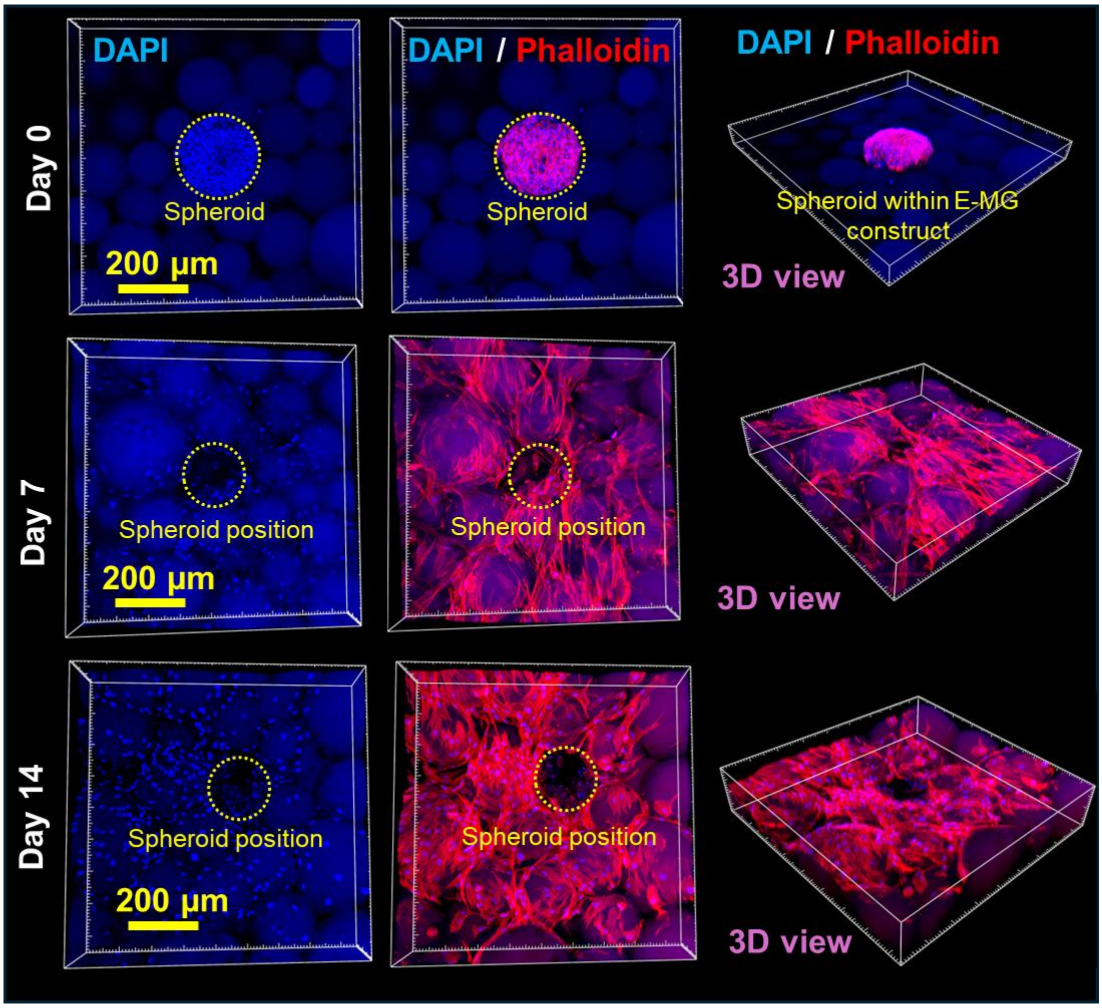


**Figure S22. Time-dependent outgrowth and dispersion of vascular spheroids within E-MG.** Representative fluorescence images and 3D view of vascular spheroids cultured within E-MG constructs at Days 0, 7 and 14. Nuclei were stained with DAPI (blue) and the actin cytoskeleton with phalloidin (red), with ovoprotein microgels visualized by intrinsic autofluorescence. The images show the initially localized spheroidal position immediately after bioprinting (Day 0), followed by progressive cellular outgrowth, proliferative expansion and invasion into the surrounding E-MG matrix during culture, accompanied by gradual loss of the original spheroid boundary and increasing cellular dispersion throughout the construct.

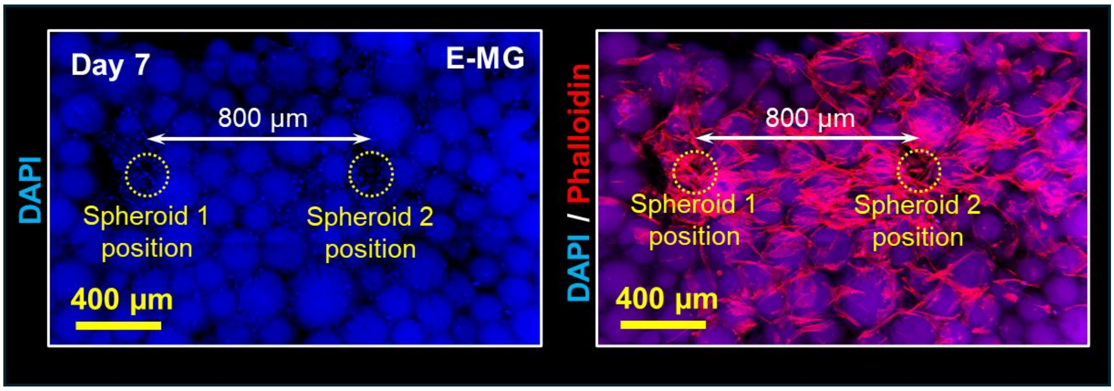


**Figure S23. Interspheroid cellular interaction and bridging within E-MG.** Representative fluorescence images of two vascular spheroids bioprinted within E-MG at an initial separation distance of 800 µm and cultured for 7 days. Nuclei were stained with DAPI (blue) and the actin cytoskeleton with phalloidin (red), with ovoprotein microgels visualized by intrinsic autofluorescence. The images show progressive cellular outgrowth from both spheroid positions and the establishment of a continuous interspheroid cellular bridge within the microgel matrix.

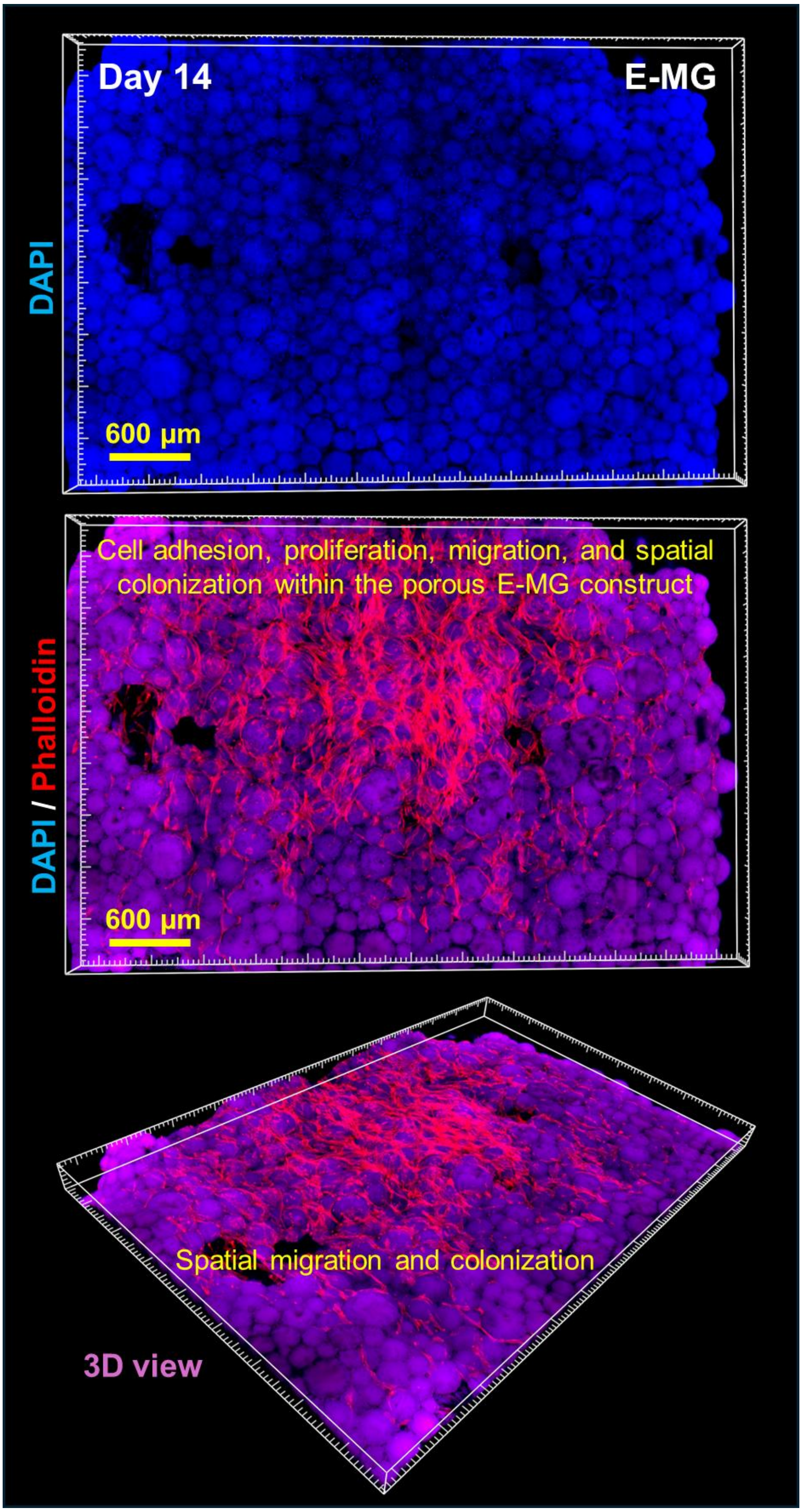


**Figure S24. Construct-scale cellular migration and colonization within porous E-MG.** Representative large-area fluorescence images and 3D view of bioprinted vascular spheroids within E-MG after 14 days of culture. Nuclei were stained with DAPI (blue) and

F-actin with phalloidin (red), with ovoprotein microgels visualized by intrinsic autofluorescence. The images show extensive cellular adhesion, migration and spatial colonization throughout the porous E-MG matrix, indicating construct-scale cell infiltration and multicellular integration.

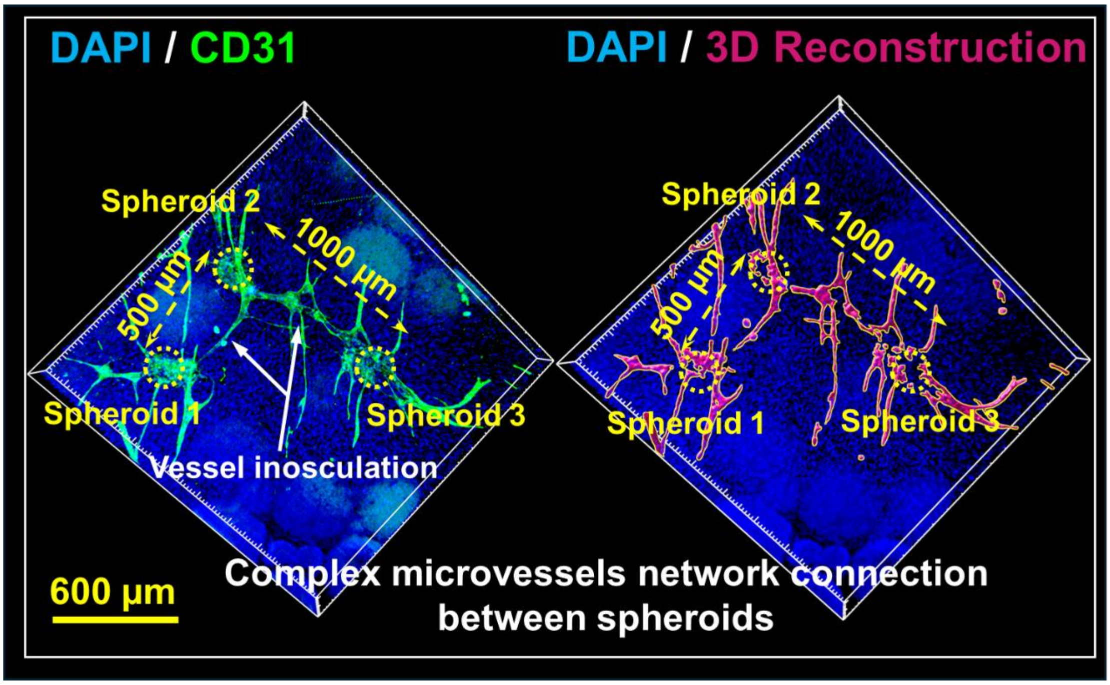


**Figure S25. Long-range microvascular interconnection among multiple spheroids within E-HG@MG.** Representative 3D fluorescence images and CD31-based reconstructions of three vascular spheroids bioprinted within E-HG@MG at predefined separation distances of 500–1000 μm after 21 days of culture. CD31 staining (green) and 3D-rendered vascular networks (magenta) show endothelial outgrowth, interspheroid vessel connection and spontaneous microvascular inosculation, forming an integrated microvascular network across the microgel-based construct.

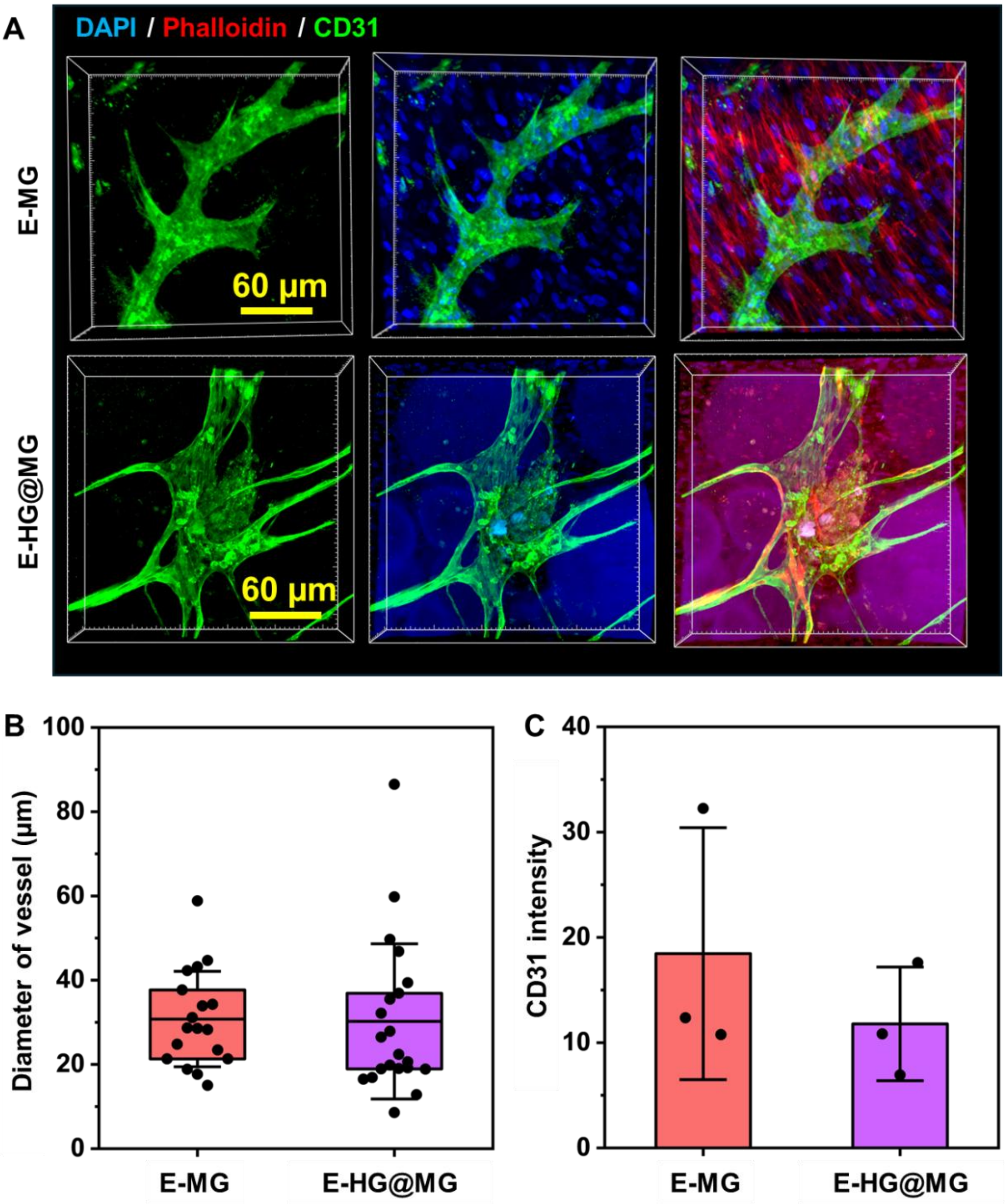


**Figure S26. Comparative microvessel formation within E-MG and E-HG@MG. A,** Representative high-resolution confocal images of $CD31^+$ vessel-like structures formed within E-MG and E-HG@MG constructs at Day 21, stained for nuclei (DAPI, blue), F-actin (phalloidin, red) and CD31 (green). Both matrices supported branched microvascular morphologies, with comparable vessel caliber and CD31 fluorescence intensity, indicating effective endothelial organization and vascular maturation within the ovoprotein microgel-based environments. **B-C,** Quantification of vessel diameter and CD31 intensity at Day 21. Data were presented as mean ± SD ($n$ = 3).

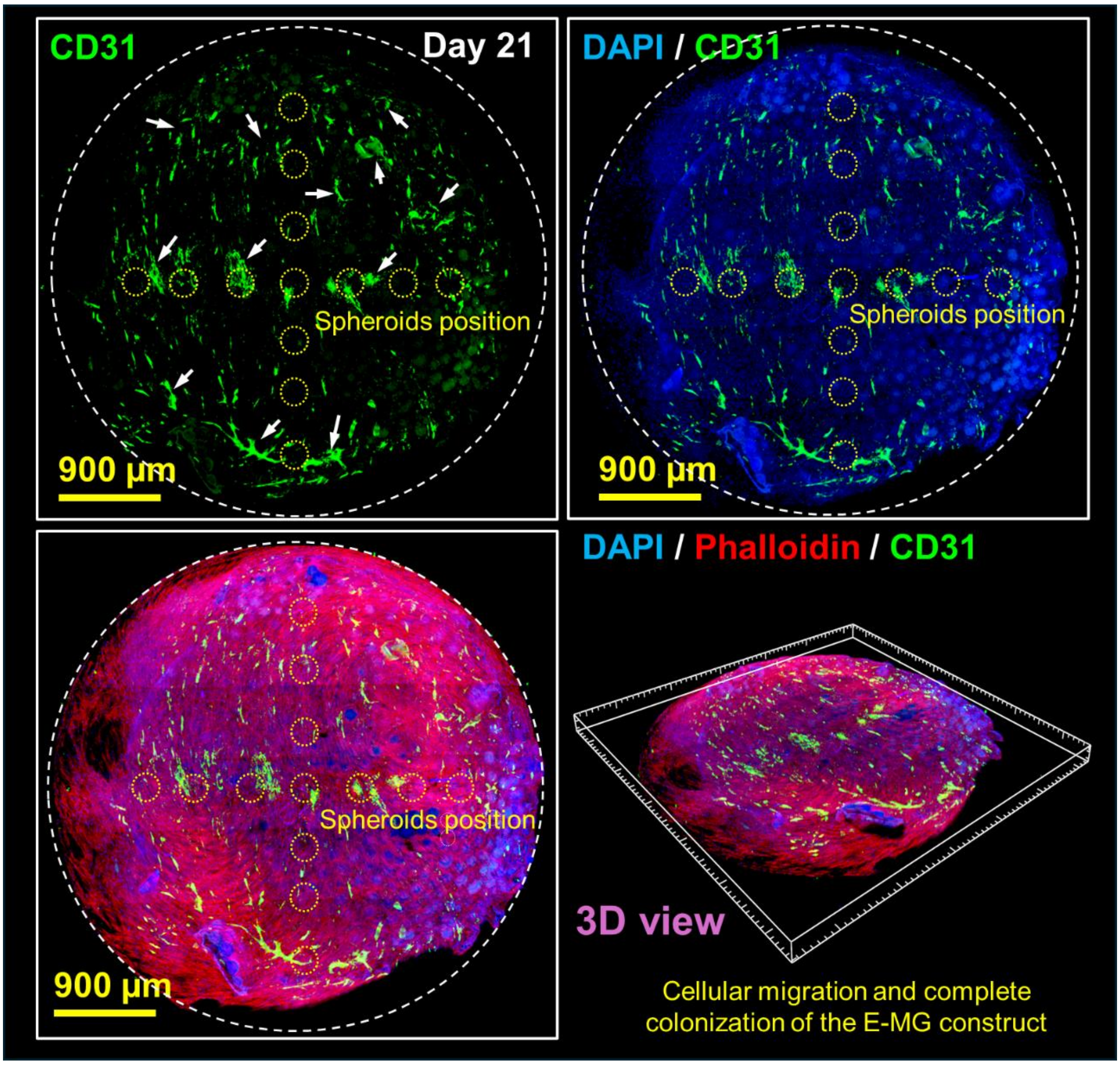


**Figure S27. Construct-scale vascular integration within the large E-MG matrix.** Representative large-area fluorescence images and 3D view of vascular spheroid-laden E-MG constructs after 21 days of culture, stained for CD31 (green), nuclei (DAPI, blue) and F-actin (phalloidin, red), with ovoprotein microgels visualized by intrinsic autofluorescence. The images reveal extensive endothelial outgrowth from the patterned spheroids, widespread cellular migration, progressive construct-wide colonization, and broad microvascular network formation throughout the ovoprotein microgel matrix, underscoring its capacity to support vascular integration across tissue-scale volumes.

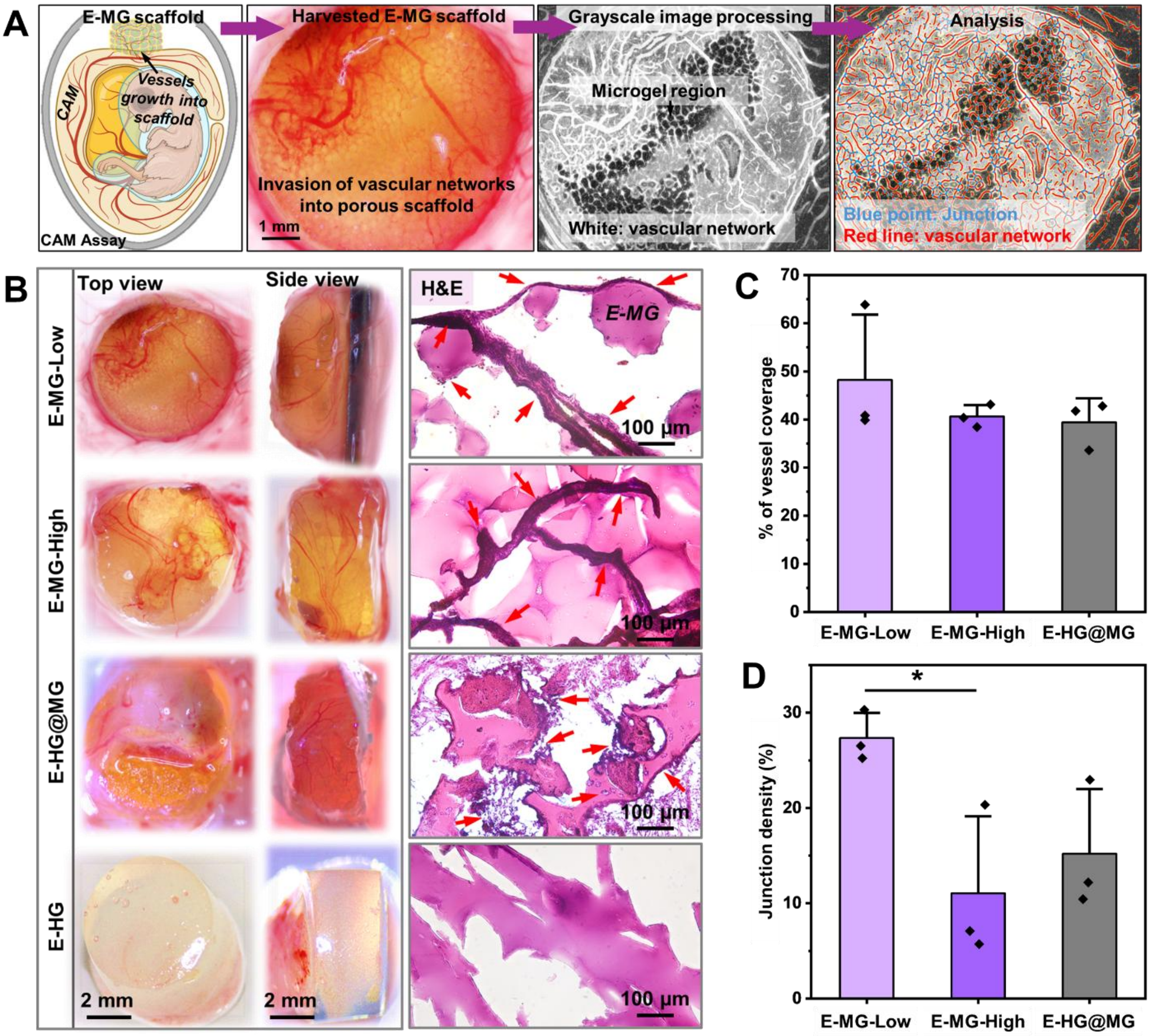


**Figure S28.** ***In ovo*** **CAM assay reveals host vascular recruitment and ingrowth into ovoprotein microgel scaffolds. A,** Schematic of the CAM assay used to evaluate the pro-angiogenic response to implanted E-MG scaffolds, together with representative images of harvested constructs, grayscale image processing and vascular network analysis, in which vessel networks and junctions were extracted for quantification. **B,** Representative top-view and side-view gross images, together with H&E-stained sections, of explanted E-MG constructs with low- and high-packing, E-HG@MG, and E-HG constructs after 7 days of incubation on the CAM. Red arrows indicate host vascular structures infiltrating the scaffold interior. Microgel-based groups show pronounced vascular recruitment and penetration throughout the construct, whereas bulk E-HG exhibits comparatively limited vascular ingrowth. **C-D,** Quantification of vessel coverage (C) and junction density (D) within the implanted constructs, confirming enhanced vascular invasion and network formation in ovoprotein microgel-based scaffolds. Data were presented as mean ± SD ($n$ = 3); $*P$ < 0.05.

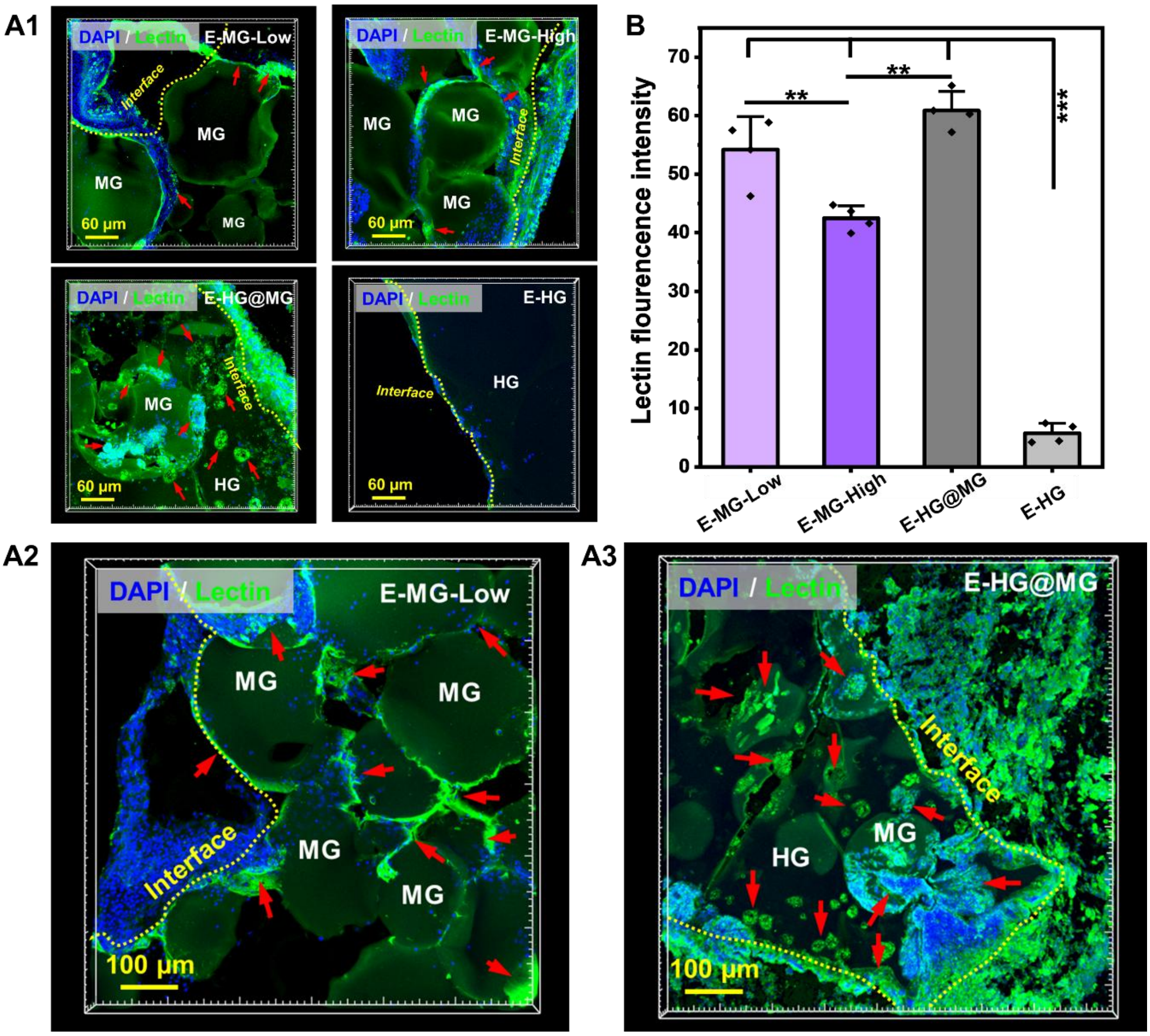


**Figure S29. Lectin staining reveals host vascular ingrowth and spatial distribution within ovoprotein microgel constructs. A1,** Representative confocal images of implanted E-MG-Low, E-MG-High, E-HG@MG and E-HG constructs stained for lectin (green) and nuclei (DAPI, blue), showing the spatial localization of host-derived vascular structures relative to the scaffold interface. Red arrows indicate lectin-positive host vascular infiltration within the construct. Compared with bulk E-HG, microgel-based groups display more extensive vessel penetration and deeper vascular engagement throughout the scaffold interior. **A2-A3,** Large-area views of low-packing E-MG (**A2**) and E-HG@MG (**A3**) constructs, highlighting host vascular extension along microgel interfaces, within interparticle void spaces, and deep into the scaffold interior. **B,** Quantification of lectin fluorescence intensity, confirming significantly enhanced host vascular infiltration in ovoprotein microgel-based constructs relative to bulk E-HG. Data were presented as mean ± SD ($n$ = 3); **$P < 0.01$, ***$P < 0.001$.

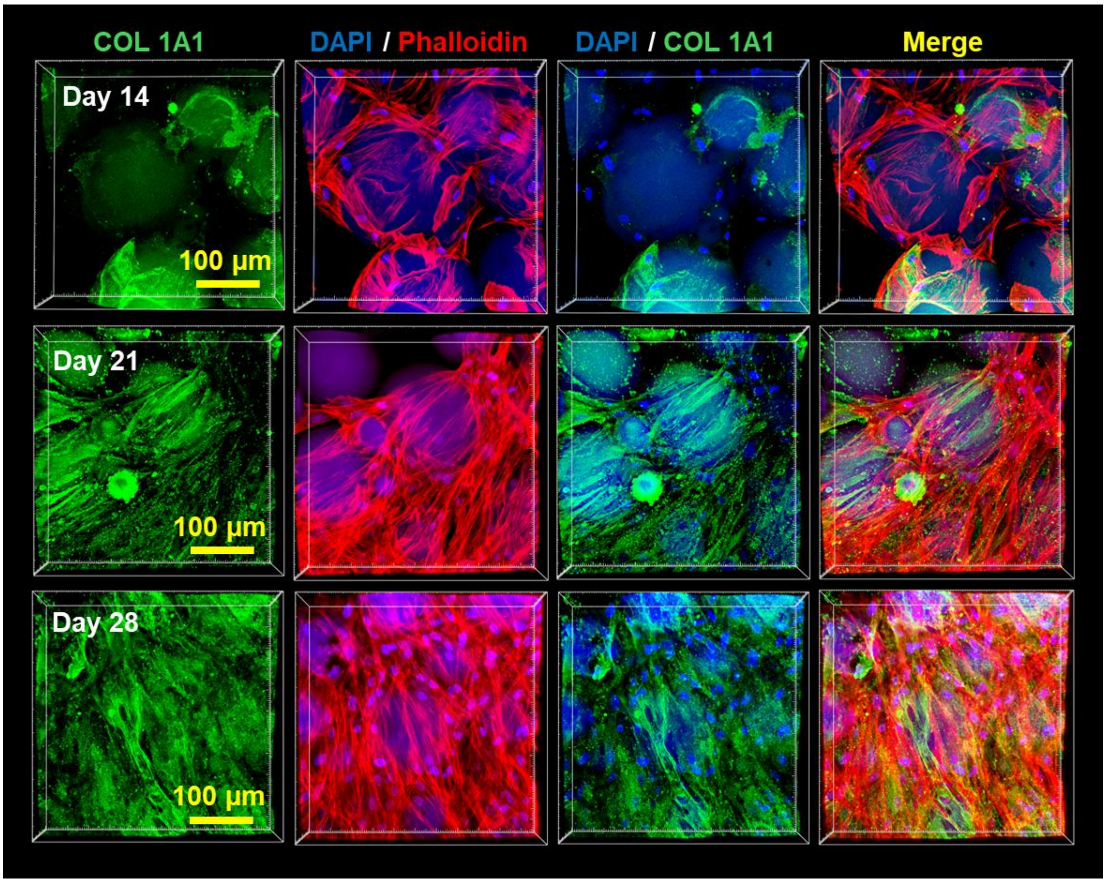


**Figure S30. Time-dependent COL1A1 deposition by bioprinted osteogenic spheroids within E-MG.** Representative immunofluorescence images of hMSC spheroid-loaded E-MG constructs at Days 14, 21 and 28, stained for COL1A1 (green), F-actin (phalloidin, red) and nuclei (DAPI, blue). Images show progressive collagenous matrix deposition and organization throughout the cellularized E-MG architecture during osteogenic culture.

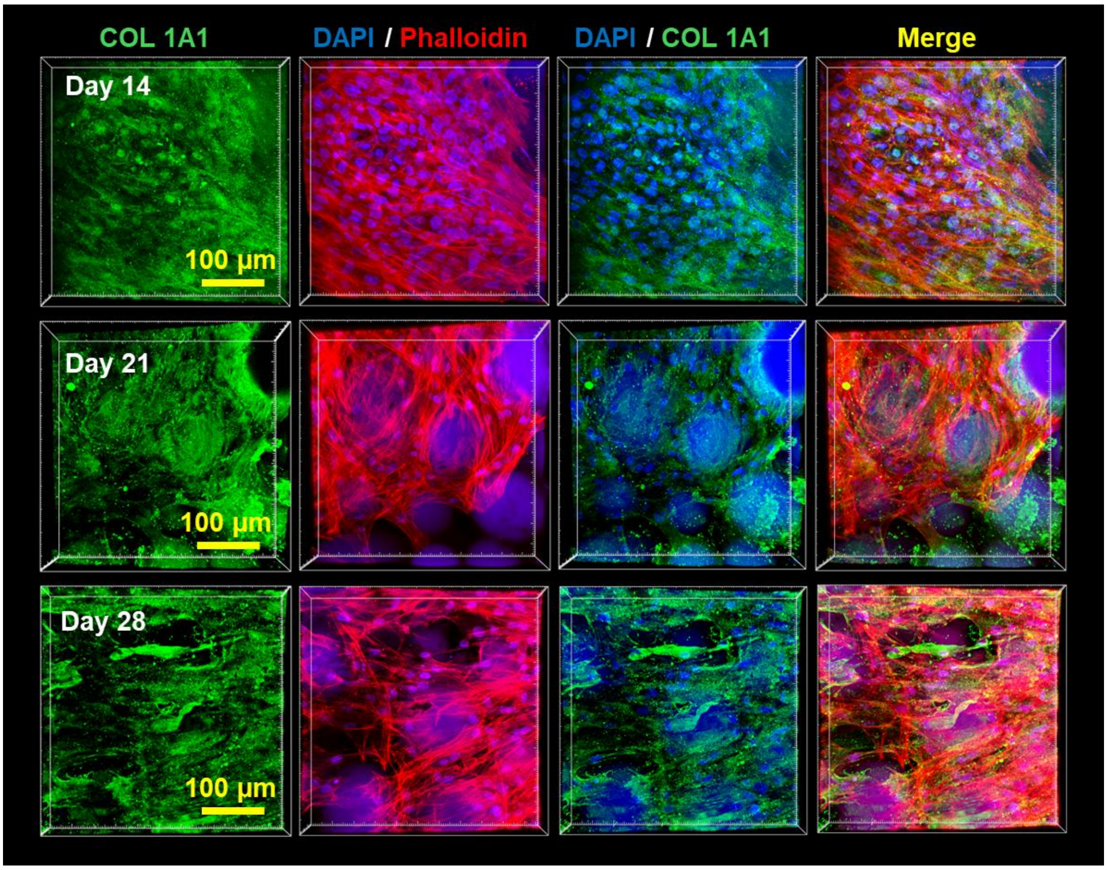


**Figure S31. Time-dependent COL1A1 deposition by bioprinted osteogenic spheroids within E-HG@MG.** Representative immunofluorescence images of hMSC spheroid-loaded E-HG@MG constructs at Days 14, 21 and 28, stained for COL1A1 (green), F-actin (phalloidin, red) and nuclei (DAPI, blue). Images show progressive collagenous matrix deposition and organization throughout the cellularized E-HG@MG architecture during osteogenic culture.

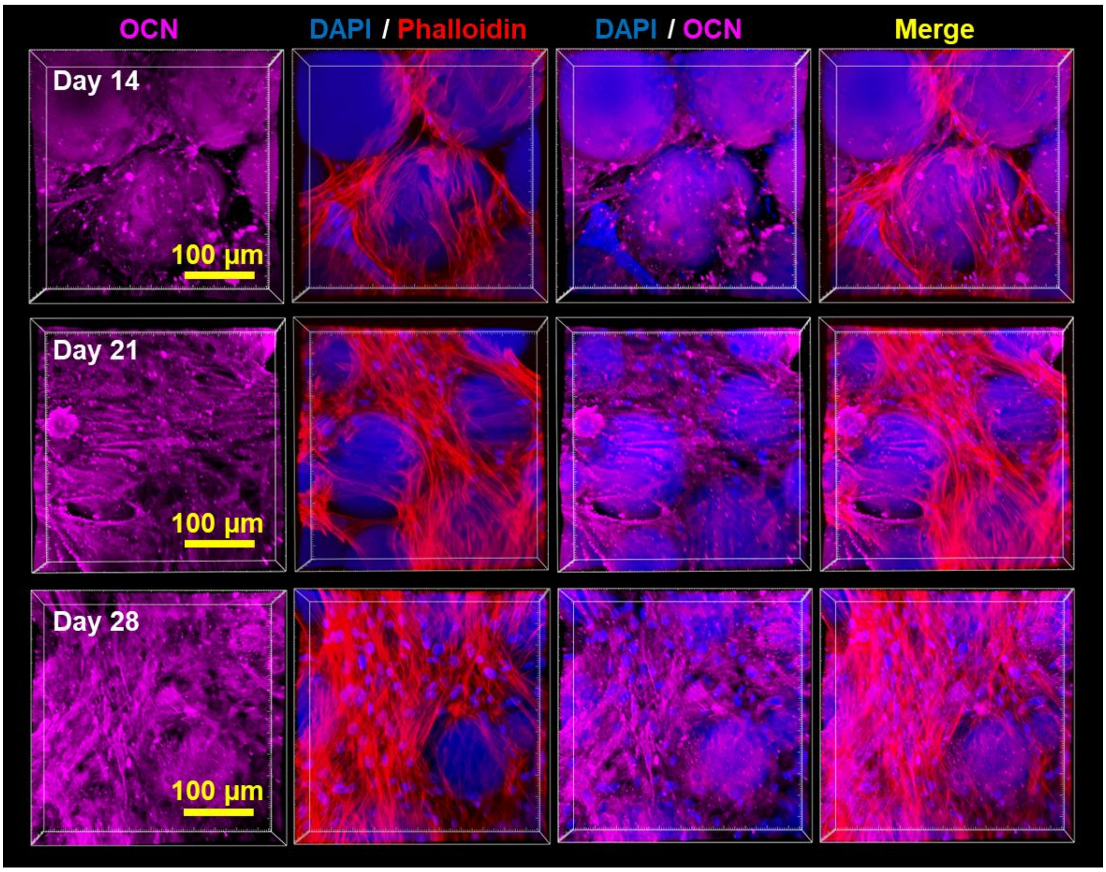


**Figure S32. Time-dependent OCN deposition by bioprinted osteogenic spheroids within E-MG.** Representative immunofluorescence images of hMSC spheroid-loaded E-MG constructs at Days 14, 21 and 28, stained for OCN (violet), F-actin (phalloidin, red) and nuclei (DAPI, blue). Images show progressive osteogenic matrix maturation and OCN deposition throughout the cellularized E-MG architecture during osteogenic culture.

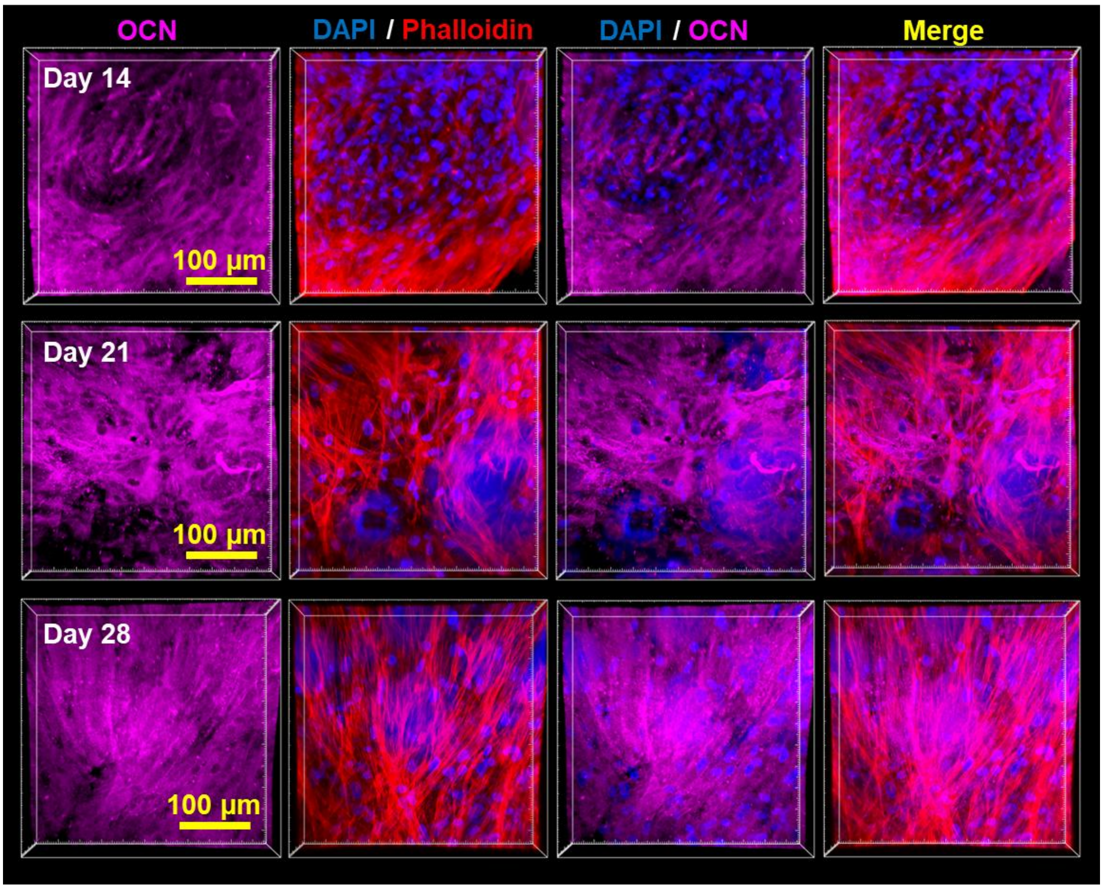


**Figure S33. Time-dependent OCN deposition by bioprinted osteogenic spheroids within E-HG@MG.** Representative immunofluorescence images of hMSC spheroid-loaded E-HG@MG constructs at Days 14, 21 and 28, stained for OCN (violet), F-actin (phalloidin, red) and nuclei (DAPI, blue). Images show pronounced early OCN deposition at Day 14, followed by progressive matrix organization and sustained OCN accumulation throughout the cellularized E-HG@MG architecture during osteogenic culture.

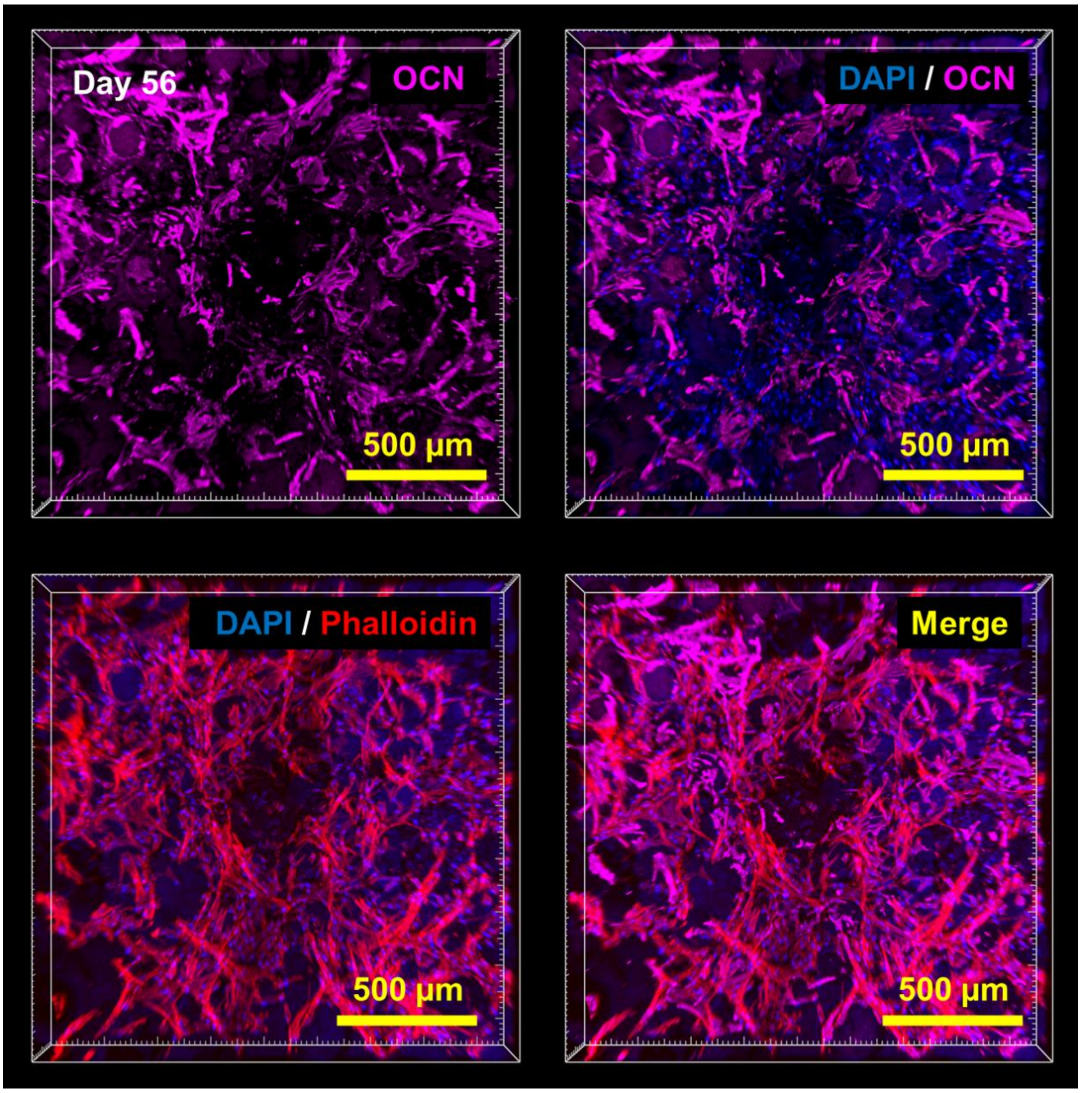

**Figure S34. Long-term OCN deposition by bioprinted osteogenic spheroids within E-MG.** Representative immunofluorescence images of hMSC spheroid-loaded E-MG constructs after 56 days of osteogenic culture, stained for OCN (violet), F-actin (phalloidin, red) and nuclei (DAPI, blue). Images show sustained OCN-rich matrix deposition and extensive cellular organization throughout the E-MG construct during long-term osteogenic maturation.

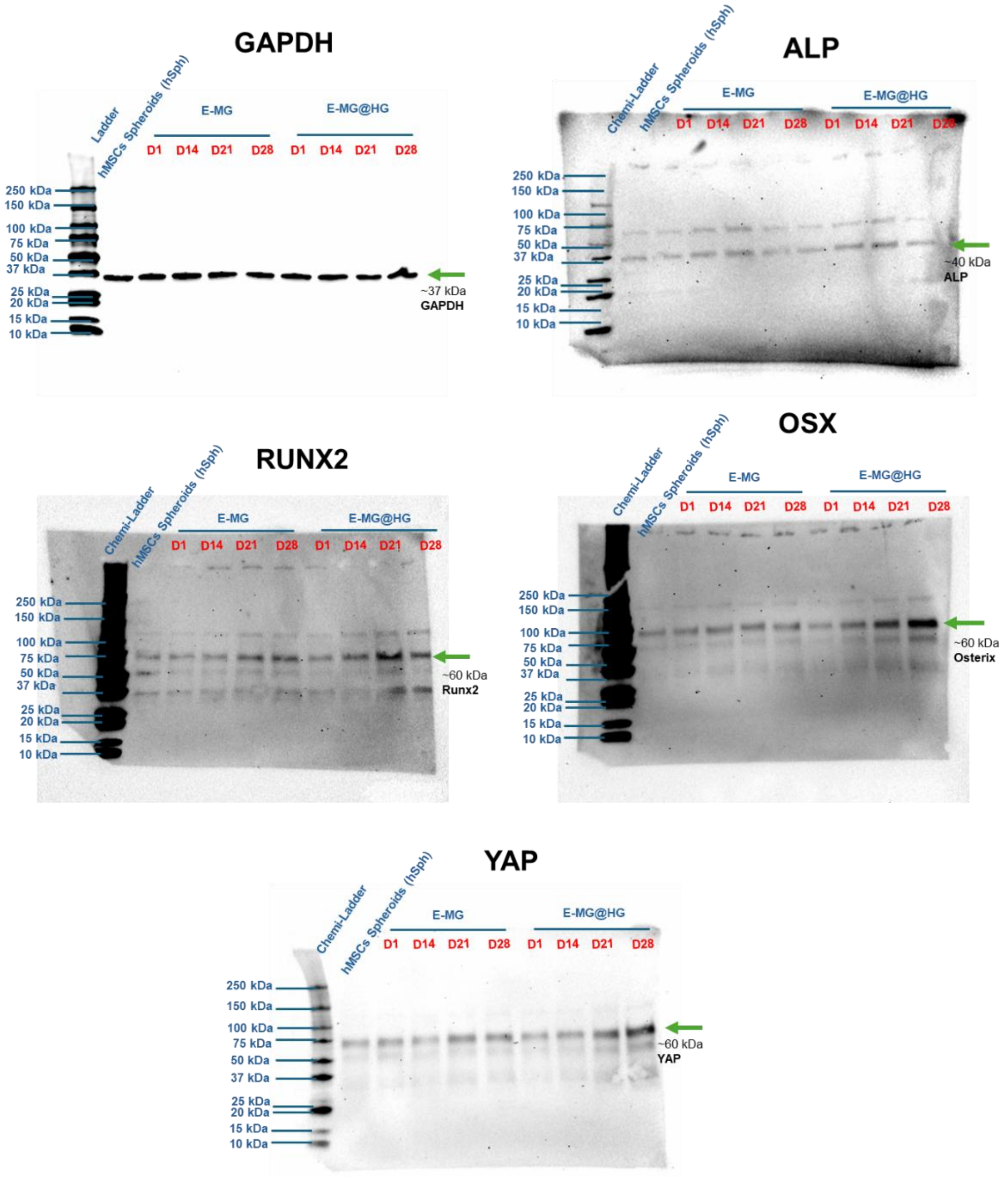


**Figure S35. Uncropped western blots for osteogenic and mechanotransductive protein analysis.** Uncropped immunoblots corresponding to Fig. 6B, showing GAPDH, ALP, RUNX2, OSX and YAP expression in hMSC spheroid-loaded E-MG and E-MG@HG constructs during culture, together with free-standing hMSC spheroid (hSph) controls. Green arrows indicate the expected molecular-weight bands used for densitometric quantification.

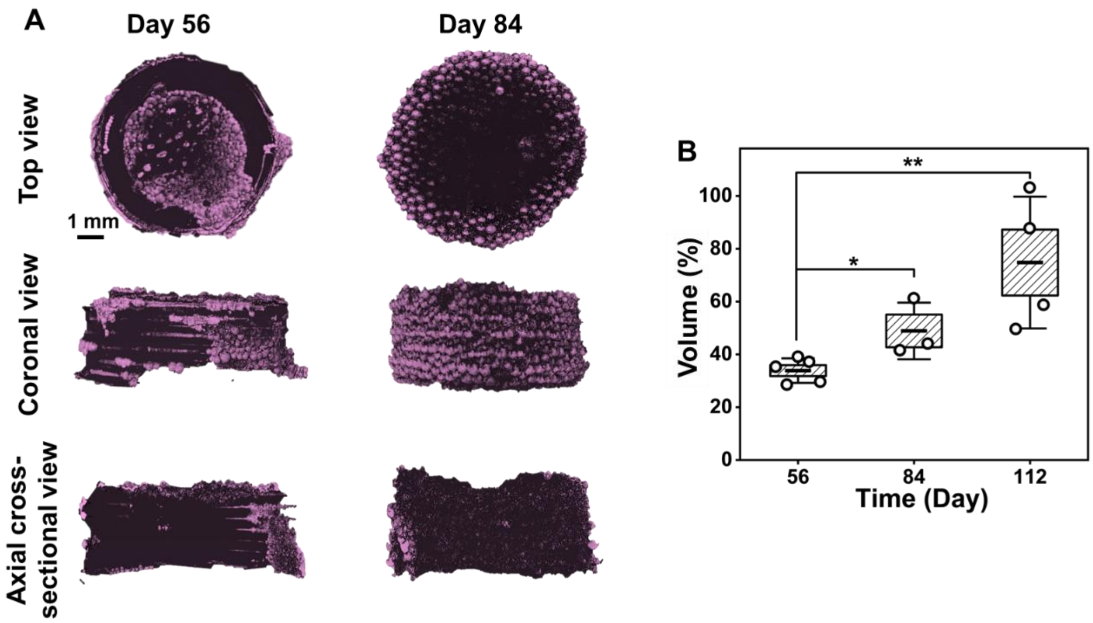


**Figure S36. Long-term μCT analysis of mineralized mini-bone constructs. A,** μCT-based 3D reconstructions of bioprinted mini-bone constructs after 56 and 84 days of osteogenic culture, shown in top, coronal and axial cross-sectional views. Mineralized regions were segmented using a grayscale threshold of 61–225 and rendered in purple, revealing progressive mineral accumulation and inward maturation over time. **B,** Quantification of mineralized volume fraction, confirming a time-dependent increase in detectable mineralized tissue. Data were presented as mean ± SD ($n \geq 3$); $^*P < 0.05$ and $^{**}P < 0.01$.

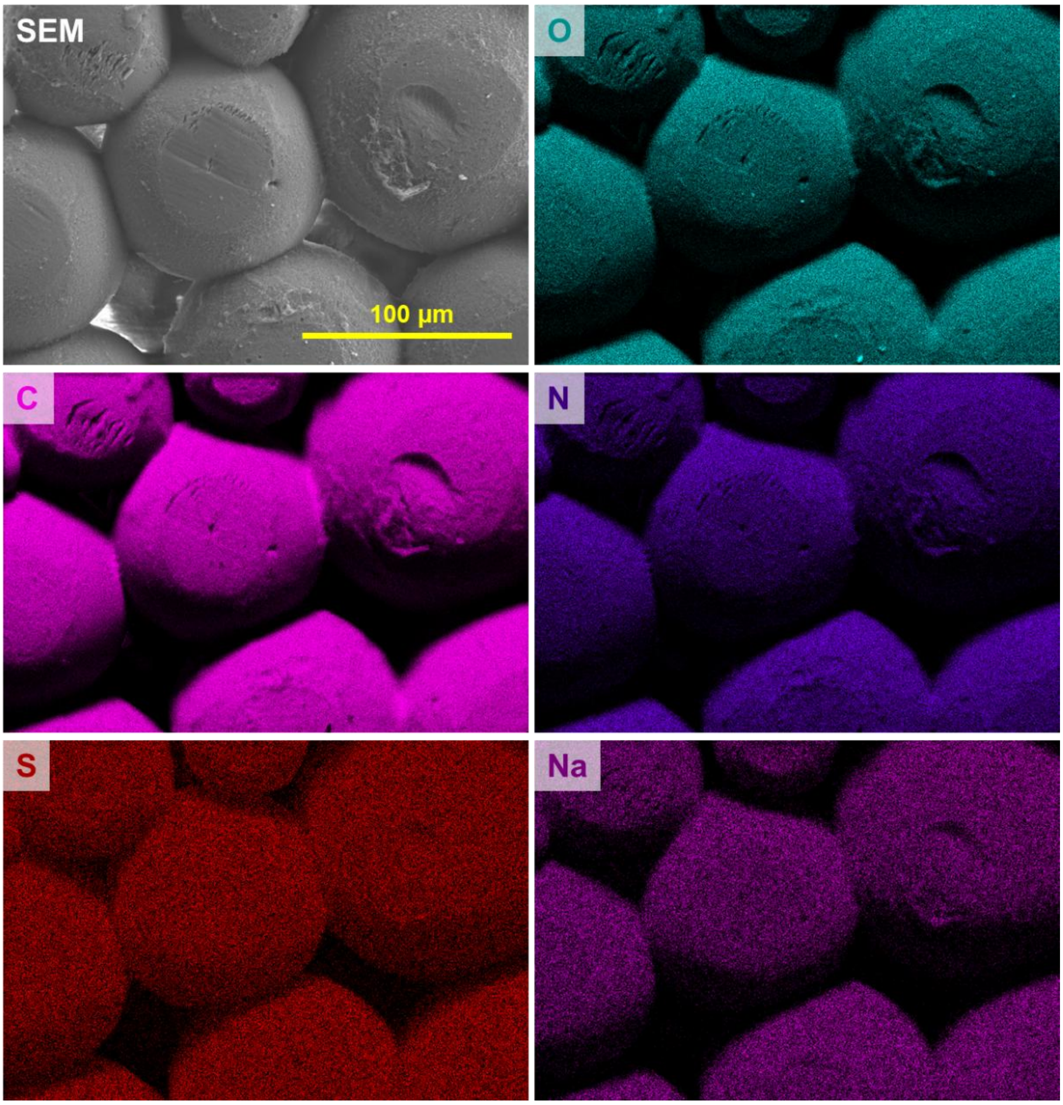


**Figure S37. SEM–EDX analysis of acellular E-MG controls.** Representative SEM image and elemental maps of cell-free E-MG constructs, showing the baseline elemental distribution of O, C, N, S and Na within the ovoprotein microgel architecture. No pronounced Ca or P enrichment was detected, supporting the cell-mediated origin of mineral-associated elemental signatures observed in spheroid-laden mini-bones.

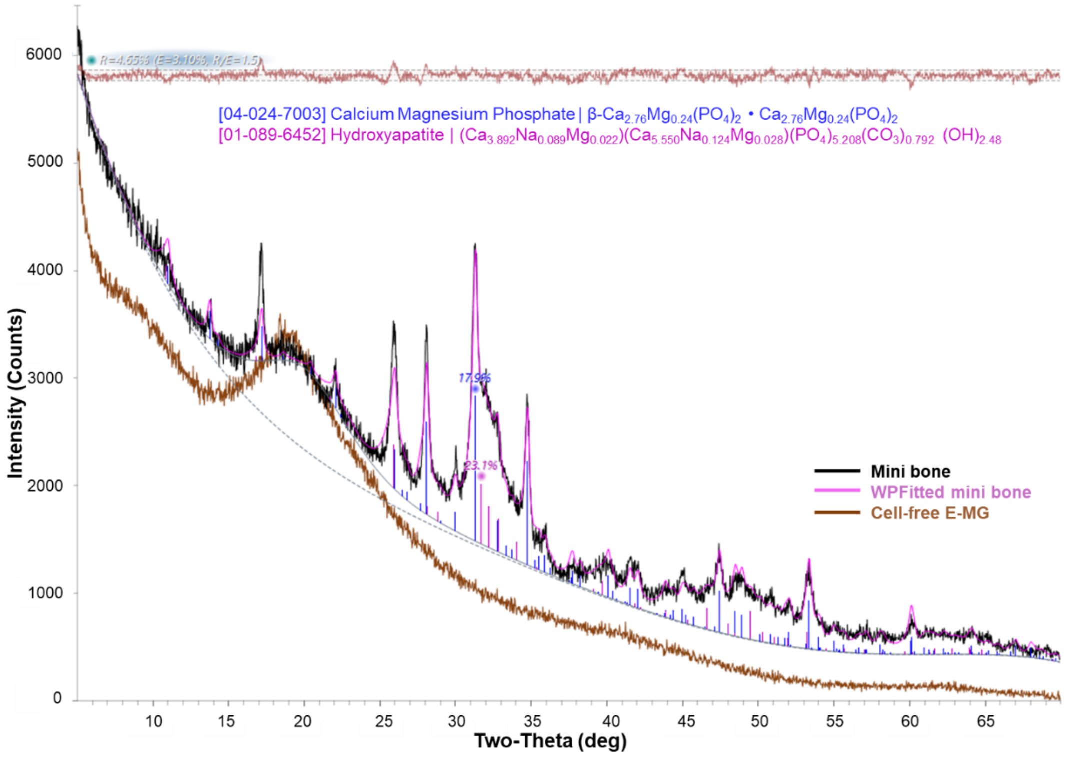


**Figure S38. XRD analysis of mineral crystalline phases in mini-bone constructs.** X-ray diffraction spectra of spheroid-loaded mini-bone after culture of 84 days and acellular E-MG controls. Whole-pattern fitting of the mini-bone spectrum identified crystalline phases assignable to hydroxyapatite-related calcium phosphate and calcium magnesium phosphate, whereas acellular E-MG showed no comparable mineral diffraction features.
