## Supplementary material for "Bioinstructive Orthogonally-crosslinked Ovoprotein Microgels for Modular Bioprinting": Description of Additional Supplementary Movies

### Description of Additional Supplementary Files

**File Name:** Supplementary Movie 1
**Description:** Interparticle photocrosslinking of E-MG, including a thin layer of E-MG matrix and one rectangular E-MG architecture.

**File Name:** Supplementary Movie 2
**Description:** Ru/SPS-mediated interparticle dityrosine photocrosslinking of E-MG assemblies.

**File Name:** Supplementary Movie 3
**Description:** DLP of acellular E-HG into a Penn State University Nittany Lion head model.

**File Name:** Supplementary Movie 4
**Description:** Aspiration-assisted bioprinting of vascular spheroids within E-MG matrices.

**File Name:** Supplementary Movie 5
**Description:** Vascularization within E-HG@MG constructs: confocal visualization and 3D reconstruction of $CD31^+$ vessel-like structures within E-HG@MG constructs at day 21, revealing hierarchical branching architectures and patent luminal compartments.

**File Name:** Supplementary Movie 6
**Description:** Mineralization of bioprinted mini-bone constructs: µCT-based 3D reconstructions of bioprinted mini-bone constructs after 56, 84 and 112 days of osteogenic culture, showing progressive mineral accumulation and inward mineral maturation over time.
